# Single-cell proteo-genomic reference maps of the hematopoietic system enable the purification and massive profiling of precisely defined cell states

**DOI:** 10.1101/2021.03.18.435922

**Authors:** Sergio H. Triana, Dominik Vonficht, Lea Jopp-Saile, Simon Raffel, Raphael Lutz, Daniel Leonce, Magdalena Antes, Pablo Hernández-Malmierca, Diana Ordoñez-Rueda, Beáta Ramasz, Tobias Boch, Johann-Christoph Jann, Daniel Nowak, Wolf-Karsten Hofmann, Carsten Müller-Tidow, Daniel Hübschmann, Theodore Alexandrov, Vladimir Benes, Andreas Trumpp, Malte Paulsen, Lars Velten, Simon Haas

**Affiliations:** Structural and Computational Biology Unit, European Molecular Biology Laboratory, 69117 Heidelberg, Germany; Collaboration for joint PhD degree between EMBL and Heidelberg University, Faculty of Biosciences, 69120 Heidelberg, Germany; Centre for Genomic Regulation (CRG), The Barcelona Institute of Science and Technology, Dr. Aiguader 88, Barcelona 08003, Spain; Heidelberg Institute for Stem Cell Technology and Experimental Medicine (HI-STEM gGmbH), 69120 Heidelberg, Germany; Division of Stem Cells and Cancer, Deutsches Krebsforschungszentrum (DKFZ) and DKFZ–ZMBH Alliance, 69120 Heidelberg, Germany; Faculty of Biosciences, Heidelberg University, 69117 Heidelberg, Germany; Department of Internal Medicine V, Heidelberg University Hospital, 69120 Heidelberg, Germany; Genome Biology Unit, European Molecular Biology Laboratory (EMBL), 69117 Heidelberg, Germany; Flow Cytometry Core Facility, European Molecular Biology Laboratory (EMBL), 69117 Heidelberg, Germany; Department of Hematology and Oncology, Medical Faculty Mannheim, University of Heidelberg, 68167 Mannheim, Germany; Computational Oncology, Molecular Diagnostics Program, National Center for Tumor diseases (NCT) Heidelberg and German Cancer Research Center (DKFZ), 69120 Heidelberg, Germany; German Cancer Consortium (DKTK), 69120 Heidelberg, Germany; Department of Pediatric Immunology, Hematology and Oncology, University Hospital Heidelberg, 69120 Heidelberg, Germany; Skaggs School of Pharmacy and Pharmaceutical Sciences, University of California, San Diego, La Jolla, CA 92093, USA; Genomics Core Facility, European Molecular Biology Laboratory (EMBL), 69117 Heidelberg, Germany; Novo Nordisk Foundation Center for Stem Cell Biology, DanStem. Faculty of Health and Medical Sciences Blegdamsvej, DK-2200 Copenhagen, Denmark; Universitat Pompeu Fabra (UPF), Barcelona, Spain; Berlin Institute of Health (BIH) at Charité – Universitätsmedizin Berlin, 10117 Berlin, Germany; Charité-Universitätsmedizin, 10117 Berlin, Germany; Berlin Institute for Medical Systems Biology, Max Delbrück Center for Molecular Medicine in the Helmholtz Association, 10115 Berlin, Germany

## Abstract

Single-cell genomics has transformed our understanding of complex cellular systems. However, excessive costs and a lack of strategies for the purification of newly identified cell types impede their functional characterization and large-scale profiling. Here, we have generated high content single-cell proteo-genomic reference maps of human blood and bone marrow that quantitatively link the expression of up to 197 surface markers to cellular identities and biological processes across all major hematopoietic cell types in healthy aging and leukemia. These reference maps enable the automatic design of cost-effective high-throughput cytometry schemes that outperform state-of-the-art approaches, accurately reflect complex topologies of cellular systems, and permit the purification of precisely defined cell states. The systematic integration of cytometry and proteo-genomic data enables measuring the functional capacities of precisely mapped cell states at the single-cell level. Our study serves as an accessible resource and paves the way for a data-driven era in cytometry.

## INTRODUCTION

Single-cell transcriptomic technologies have revolutionized our understanding of tissues (Giladi and Amit, 2018; Stuart and Satija, 2019; Tanay and Regev, 2017). The systematic construction of whole-organ and whole-organism single-cell atlases has revealed an unanticipated diversity of cell types and cell states, and has provided detailed insights into cellular development and differentiation processes (Baccin et al., 2020; Han et al., 2018, 2020; Schaum et al., 2018). However, strategies for the prospective isolation of cell populations, newly identified by single-cell genomics, are needed to enable their functional characterization or therapeutic use. Furthermore, single-cell genomics technologies remain cost-intense and scale poorly, impeding their integration into the clinical routine.

Unlike single-cell transcriptomics, flow cytometry offers a massive throughput in terms of samples and cells, is commonly used in the clinical routine diagnostics (Van Dongen et al., 2012) and remains unrivaled in the ability to prospectively isolate live populations of interest for downstream applications. However, flow cytometry provides low dimensional measurements and relies on predefined sets of surface markers and gating strategies that have evolved historically in a process of trial and error. Hence, single-cell transcriptomics (scRNA-seq) approaches have demonstrated that flow cytometry gating schemes frequently yield impure or heterogeneous populations (Paul et al., 2015; Velten et al., 2017), and flow strategies for the precise identification of cell types defined by scRNA-seq are lacking. Conversely, the precision and efficiency of commonly used cytometry gating schemes are largely unknown, and the exact significance of many surface markers remains unclear. Together, these findings highlight a disconnect between single-cell genomics-based molecular cell type maps and data generated by widely used cytometry assays.

The differentiation of hematopoietic stem cells (HSCs) in the bone marrow constitutes a particularly striking example for this disconnect (Haas et al., 2018; Jacobsen and Nerlov, 2019; Laurenti and Göttgens, 2018; Loughran et al., 2020). The classical model of hematopoiesis, which is mainly based on populations defined by flow cytometry (Akashi et al., 2000; Doulatov et al., 2010; Kondo et al., 1997) has recently been challenged in several aspects by single-cell transcriptomic (Giladi et al., 2018; Nestorowa et al., 2016; Paul et al., 2015; Tusi et al., 2018; Velten et al., 2017), functional (Notta et al., 2016; Perié et al., 2015) and lineage-tracing (Rodriguez-Fraticelli et al., 2018) approaches. These studies revealed that hematopoietic lineage commitment occurs earlier than previously anticipated, that putative oligopotent progenitors isolated by FACS consist of heterogeneous mixtures of progenitor populations, and that lineage commitment is most accurately represented by a continuous process of differentiation trajectories rather than by a stepwise differentiation series of discrete progenitor populations (Haas, 2020; Haas et al., 2018; Jacobsen and Nerlov, 2019; Laurenti and Göttgens, 2018). The frequency of functionally oligopotent progenitors in immunophenotypic hematopoietic stem and progenitor gates still remains controversial (Karamitros et al., 2018; Psaila et al., 2016; Velten et al., 2017). These discrepancies have contributed to conflicting results between studies that employ scRNA-seq for the definition of progenitor populations (Giladi et al., 2018; Paul et al., 2015; Pellin et al., 2019; Tusi et al., 2018; Velten et al., 2017) and studies that use FACS (Akashi et al., 2000; Kondo et al., 1997; Pei et al., 2017). As a consequence, flow-based assays that accurately reflect the molecular and cellular complexity of the hematopoietic system are urgently needed.

Recently, methods to simultaneously measure mRNA and surface protein expression in single cells have been developed (Shahi et al., 2017; Stoeckius et al., 2017). Here, we demonstrate that ultra-high content single-cell proteo-genomic reference maps, alongside appropriate computational tools, can be used to systematically design and analyze cytometry assays that accurately reflect scRNA-seq based molecular tissue maps at the level of cell types and differentiation states. For this purpose, we have generated proteo-genomic datasets encompassing 97-197 surface markers across 122,004 cells representing the cellular landscape of young, aged and leukemic human bone marrow and blood, as well as all states of hematopoietic stem cell differentiation. We demonstrate how such data can be used in an unbiased manner to evaluate and automatically design cytometry gating schemes for individual populations and entire biological systems without prior knowledge. We show that, compared to existing approaches, such optimized schemes are superior in the identification of cell types and more accurately reflect molecular cell states. Projecting datasets from malignant hematopoiesis on our reference atlases enables the fine-mapping of the exact stage of differentiation arrest in leukemias, the identification of leukemia-specific surface markers and an unsupervised classification of disease states. Finally, we demonstrate how such data resources can be used to project low-dimensional cytometry data on single-cell genomic atlases to enable functional analysis of precisely defined states of cellular differentiation. Our data resource and bioinformatic advances enable the efficient identification and isolation of any molecularly defined cell state from blood and bone marrow while laying the grounds for reconciling flow cytometry and single-cell genomics data across human tissues.

## RESULTS

### A comprehensive single-cell proteo-genomics reference map of young, aged and malignant bone marrow

To establish a comprehensive single-cell transcriptomic and surface protein expression map in the human bone marrow (BM), we performed a series of Abseq experiments, in which mononuclear BM cells from hip aspirates were labelled with 97-197 oligo-tagged antibodies, followed by targeted or whole transcriptome scRNAseq on the Rhapsody platform (Figure 1a). For targeted single-cell transcriptome profiling, we established a custom panel, consisting of 462 mRNAs covering all HSPC differentiation stages, cell type identity genes, mRNAs of surface receptors and additional genes that permit the characterization of cellular states. These genes were systematically selected to capture all relevant layers of RNA expression heterogeneity observed in this system (Supplementary Note 1 and Supplementary Table 1). Whole transcriptome single-cell proteo-genomics confirmed that no populations were missed due to the targeted nature of the assay (Supplementary Note 2). Using this panel, in combination with 97 surface markers (Supplementary Table 2), we analyzed the BM of three young healthy donors, three aged healthy donors, and three acute myeloid leukemia (AML) patients at diagnosis (Figures 1a, S1, Supplementary Table 3). For samples from healthy donors, CD34+ cells were enriched to enable a detailed study of hematopoietic stem cell (HSC) differentiation (Figure S2). For samples from AML patients, CD3+ cells were enriched in some cases to ensure sufficient coverage of T cells.

**Figure 1.**
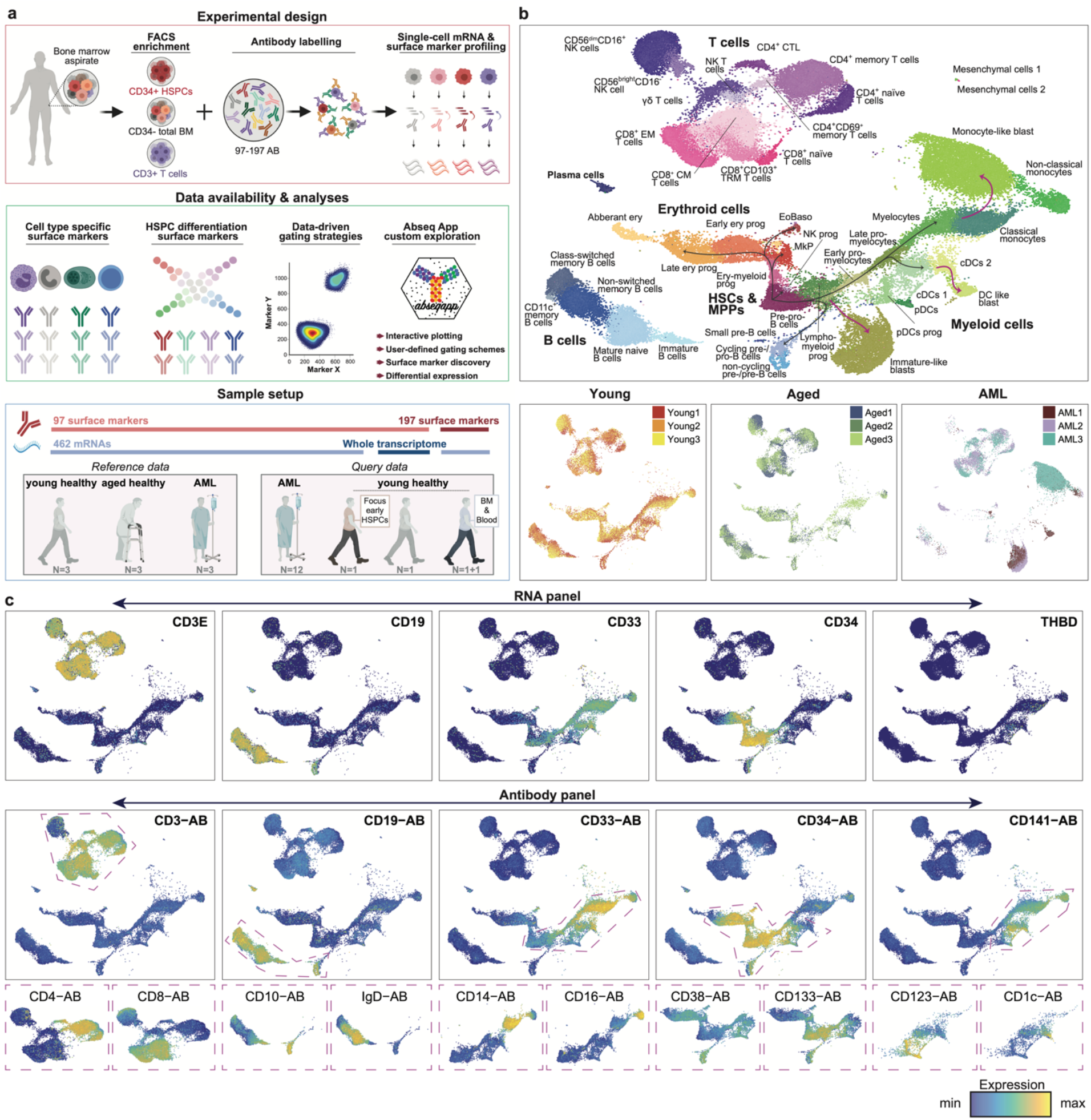
A comprehensive single-cell proteo-genomic map of young, aged and malignant bone marrow. **a**. Overview of the study. See methods and main text for details. **b**. Top: UMAP display of single-cell proteo-genomics data of human bone marrow from healthy young, healthy aged and AML patients (n=70,017 single cells, 97 surface markers), integrated across n=9 samples and data modalities. Clusters are color-coded. Bottom: UMAPs highlighting sample identities. See *Supplementary Note 5* for details on cluster annotation. The whole transcriptome Abseq data is presented in *Supplementary Note 2*, the Abseq experiments with measurements of 197 surface markers are presented in *Figure S4*. **c**. Normalized expression of selected mRNAs and surface proteins highlighted on the UMAP space from *b*. Top: Expression of mRNAs encoding surface markers widely used to identify major cell types. Middle: Expression of the corresponding surface proteins. Bottom: Expression of markers widely used to stratify major cell types into subtypes. Only the parts of the UMAPs highlighted by dashed polygons in the middle row are shown. For all data shown, bone marrow mononuclear cells from iliac crest aspirations from healthy adult donors or AML patients were used.

Since single-cell proteo-genomic approaches are not commonly performed at this level of antibody multiplexing, we designed a series of control experiments. First, we performed matched Abseq experiments in the presence or absence of antibodies to ensure that highly multiplex antibody stains do not impact on the transcriptome of single cells (Supplementary Note 3). We further performed a series of AbSeq experiments on fresh and frozen samples to demonstrate that the freeze-thawing process has no major impact on the data (Supplementary Note 3). Finally, we evaluated the sequencing requirements for optimal cell type classification in high-parametric single-cell proteo-genomic experiments (Supplementary Note 4). In the main reference data set, 70,017 high-quality BM cells were profiled with combined RNA and high-parametric surface protein information, and an average of ∼7500 surface molecules per cell were detected (Figure S3). Following data integration across experiments and measurement modalities, we identified 45 cell types and cell stages covering the vast majority of previously described hematopoietic cell types of the bone marrow and peripheral blood, including all stages of HSC differentiation in the CD34+ compartment, all T cell and NK cell populations of the CD3+ and CD56+ compartments, several dendritic cell and monocyte subpopulations from the CD33+ compartment and all major B cell differentiation states across CD10+, CD19+ and CD38^high^ compartments (Figures 1b,c Supplementary Note 5, Supplementary Table 4). In addition, poorly characterized populations, such as cytotoxic CD4+ T cells and mesenchymal stem or stromal cells (MSCs) are covered. Cells from young and aged bone marrow occupied the same cell states in all individuals, whereas cell states in AML differed (Figure 1b and see below). Importantly, the combined RNA and surface protein information provided higher resolution and revealed cell types that are not readily identified by one of the individual data layers alone (Supplementary Note 6).

Besides our main reference dataset, we have generated ‘query’ single-cell proteo-genomic datasets which are displayed in the context of the main reference (Supplementary Note 7). These include, first, the analyses of healthy BM and matched peripheral blood (PB) samples using a 197 plex antibody panel to query the expression of additional surface markers in the context of our reference (Figure S4, Supplementary Table 2). Second, the analyses of healthy BM analyzed with a 97 plex antibody panel in combination with whole transcriptome profiling to query any gene’s expression in the space defined by our reference (Supplementary Note 2). Third, the profiling of the CD34^+^CD38^-^ bone marrow compartment with a 97 plex antibody panel to provide higher resolution of immature HSPCs (see below, Figure S9c, d) and fourth, a cohort of 12 AML patients (see below, Figure 4). To make our comprehensive resource accessible, we developed the Abseq-App, a web-based application that permits visualization of gene and surface marker expression, differential expression testing and the data-driven identification of gating schemes across all datasets presented in this manuscript. A demonstration video of the app is available in the supplement (Supplemental Video S1). The Abseq-App is accessible at: https://abseqapp.shiny.embl.de/.

### Systematic association of surface markers with cell type identities, differentiation stages and biological processes

While surface markers are widely used in immunology, stem cell biology and cancer research to identify cell types, cell stages and biological processes, the exact significance of individual markers remains frequently ambiguous. To quantitatively link surface marker expression with biological processes, we assigned each cell in our data set to its respective cell type, and determined its differentiation stage, its stemness score, its cytotoxicity score, its current cell cycle phase as well as technical covariates (see Methods and below). Moreover, we included covariates representing unknown biological processes that were defined in an unsupervised manner using a factor model. Non-technical covariates were not affected by marker expression level (Figure S5a, Methods). For each surface marker, we then quantified the fraction of variance of expression that is determined by any of these processes (Figure 2a). This model identified markers that represent cell type identities or differentiation stages, as well as stemness, cytotoxicity and cell cycle properties (Figures 2b-d and S5b-f).

**Figure 2:**
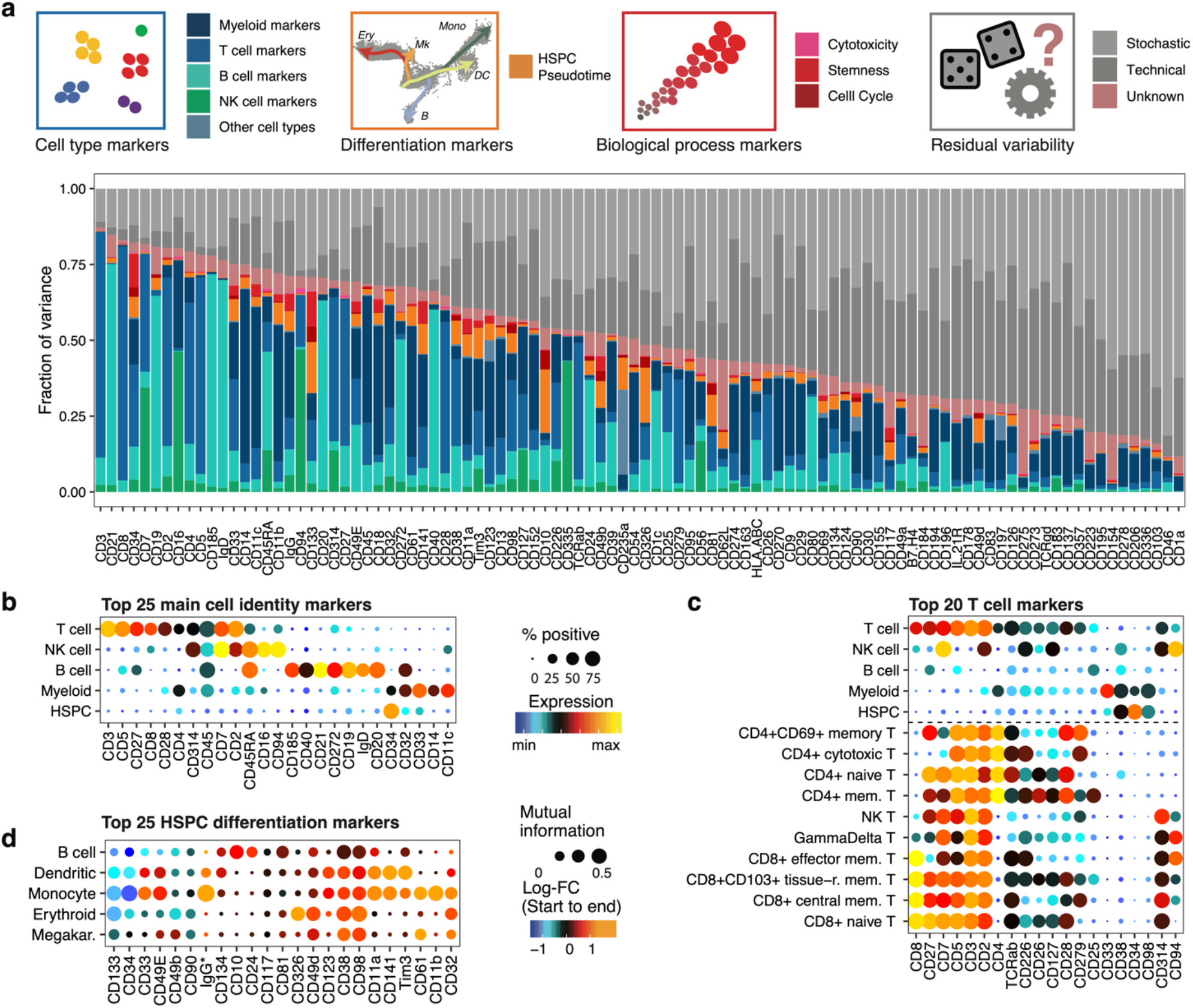
Association of surface marker expression with cell type identities, cellular differentiation, and biological processes. **a**. For each surface marker measured in our 97-plex Abseq data, the fraction of variance explained by different covariates (colored insets in top row) is displayed. For this, every single cell from healthy young individuals (n=3 samples, 28,031 single cells) was assigned to a cell type identity (blue inset, see *Figure 1b*), and cytotoxicity, stemness and cell cycle scores (red inset, see *Figure S5e*), as well as technical covariate scores were determined. Additionally, pseudotime analyses were used to assign differentiation scores to HSPCs (orange inset, see *Figure 3a*). These covariates were then used to model surface marker expression in a linear model. The fraction of variance explained by each of the processes was quantified. See Methods, section ‘*Modelling variance in surface marker expression’* for details. **b**. Cell type identity markers. Dot plot depicting the expression of the 25 surface markers with the highest fraction of variance explained by cell type across major populations. Colors indicate mean normalized expression, point size indicates the fraction of cells positive for the marker. Automatic thresholding was used to identify positive cells, see Methods, section ‘*Thresholding of surface marker expression’* for details. **c**. T cell subtype markers. The expression of the 20 surface markers with the highest fraction of variance explained by T cell subtype is displayed, legends like in *b*. **d**. HSPC differentiation markers. Dot plot depicting expression changes of markers across pseudotime in CD34+ HSPCs. Color indicates logarithmic fold change between the start and the end of each pseudotime trajectory. Point size indicates the mutual information in natural units of information (nats) between pseudotime and marker expression. The 25 surface markers with the highest fraction of variance explained by pseudotime covariates are displayed.

To characterize novel markers identified by this analysis, we initially focused on the evaluation of surface molecules that specifically mark distinct stages of HSC differentiation, since a lack of specific markers currently impedes the accurate representation of lineage commitment by flow cytometry (Notta et al., 2016; Paul et al., 2015; Pellin et al., 2019; Tusi et al., 2018; Velten et al., 2017). For this purpose, we performed pseudotime analyses within the CD34+ HSPC compartment and identified surface markers that correlate with the progression of HSCs towards erythroid, megakaryocytic, monocyte, conventional dendritic cell or B cell differentiation trajectories (Figure 2d, 3a, S5g and see Methods). Of note, the monocyte trajectory also includes neutrophil progenitor stages, but mature neutrophils are not included in the datasets due to the use of density gradient centrifugation of samples. Moreover trajectory analyses were not performed for plasmacytoid dendritic and eosinophil/basophil lineages, due to a low number of intermediate cells impeding an unanimous identification of branch points. Pseudotime analyses quantified the exact expression dynamics of many well-established markers, such as CD38 as a pan-differentiation marker, as well as CD10 and CD11c as early B cell and monocyte-dendritic cell lineage commitment marker, respectively (Figures 2d, and S6a). Importantly, our analyses revealed novel surface markers that specifically demarcate distinct stages of lineage commitment, including CD326, CD11a and Tim3 (Figure 2d and 3). To confirm the high specificity of these markers for erythroid and myeloid commitment, respectively, we used FACS-based indexing of surface markers coupled to single-cell RNAseq (“index scRNAseq”, see also Supplementary Note 8), or coupled to single-cell cultures (“index cultures”) (Figure 3b). As suggested by our proteo-genomic single-cell data, CD326 expression was associated with molecular priming and functional commitment into the erythroid lineage (Figure 3c-g and S6b, c). In contrast, Tim3 and CD11a were identified as pan-myeloid differentiation markers and were associated with transcriptomic priming and functional commitment into the myeloid lineage (Figures 3c, h-o and S6c). Finally, CD98 was identified as a novel pan-differentiation marker of HSCs, which we confirmed by classical flow cytometry (Figures 2d, and S6d-h). Beyond the progression of HSCs to lineage committed cells, we also analyzed the surface marker dynamics throughout B cell differentiation, allowing us to identify markers specific to their lineage commitment, maturation, isotype switching and final plasma cells generation (Figure S6i-p).

**Figure 3.**
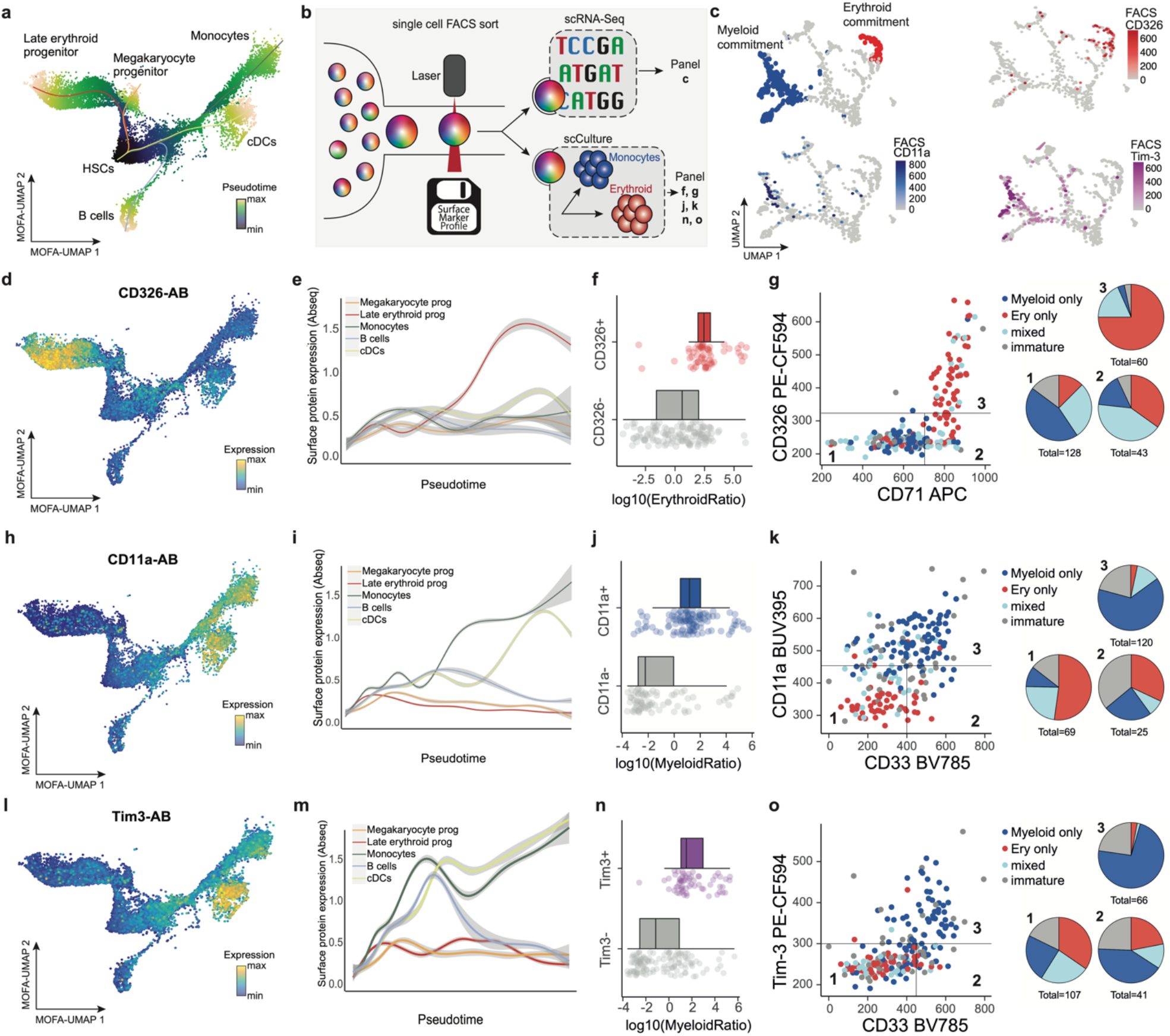
Validation of novel stage-specific HSPC differentiation markers. **a**. UMAP plot depicting CD34+ HSPCs and their pseudotime scores along five differentiation trajectories, see Methods, section “*Pseudotime analysis”*. The normalized pseudotime score across all lineages is color-coded. **b**. Scheme illustrating the experiments performed to validate the significance of selected markers. See main text and *Supplementary Note 8* for details. **c**. UMAP display of mRNA expression of n=630 CD34+ cells from a single-cell Smart-seq2 experiment where surface markers were recorded using FACS. For a detailed description of the experiment, see *Supplementary Note 8*. Upper left panel: Cells with myeloid and erythroid gene expression signatures are highlighted on the UMAP. Bottom-left and right panels: Surface protein expression (FACS data) of indicated markers is shown. **d**. UMAP display highlighting the normalized CD326 surface protein expression (Abseq data). **e**. Line plots depicting normalized CD326 surface protein expression (Abseq data) smoothened over the different pseudotime trajectories illustrated in panel a. **f**. Boxplots depicting the ratio in erythroid cells produced in single-cell cultures in relation to the CD326 expression of the founder cell (n=231 single cell derived colonies). **g**. Left panel: scatter plots depicting the differentiation potential of single founder cells in relation to their CD326 and CD71 surface expression. The founder cell potential was categorized by its ability to give rise to 1) erythroid only progeny, 2) a mix of erythroid, myeloid or any other progeny, 3) only myeloid progeny 4) remaining cells. Right panel: Founder cells were subset according to their CD326 and CD71 surface expression status and relative fractions of their respective potential are summarized as pie charts. **h-o**. Like *d-g*, except that CD11a *(h-k)* or Tim3 *(l-o)* and their relation to the formation of myeloid cells in single-cell cultures is investigated (n=214 single cell derived colonies). For all data presented, bone marrow mononuclear cells from iliac crest aspirations from healthy adult donors were used.

Together, our model provides a global and quantitative understanding of how well cell type identities, differentiation stages and biological processes are related to the expression of individual surface markers. A comprehensive overview of surface markers associated with these processes is depicted in the supplement (Supplementary Table 5, Figure S5).

### Adaptation of surface protein expression in healthy aging and cancer

To investigate the surface protein expression throughout healthy aging, we compared Abseq data of bone marrow from young and aged healthy individuals. These analyses revealed that the expression of surface molecules was highly similar across all BM populations between the age groups (Figures 4a, b, Supplementary Table 6), suggesting unexpectedly stable and highly regulated patterns of surface protein expression that are only modestly affected by aging. While cell type frequencies were also only modestly affected by aging, a significant accumulation of cytotoxic effector CD8+ T cells was observed (Figure S7a, Fagnoni et al., 1996). Moreover, the expression of several immune regulatory molecules showed age-related changes in surface presentation, including the death receptor FAS (CD95), the poliovirus receptor (CD155) and the ICOS ligand (CD275) (Figure 4b). In particular, naive CD8+ and CD4+ T cell subsets displayed an aging-associated decline of CD27 surface expression, a co-stimulatory molecule required for generation and maintenance of long-term T cell immunity (Figures 4b, c, Peters et al., 2015). Together these analyses suggest that the overall pattern of surface protein expression is widely maintained upon healthy aging, whereas specific changes, most prominently in the surface presentation of immune regulatory molecules, occur.

**Figure 4.**
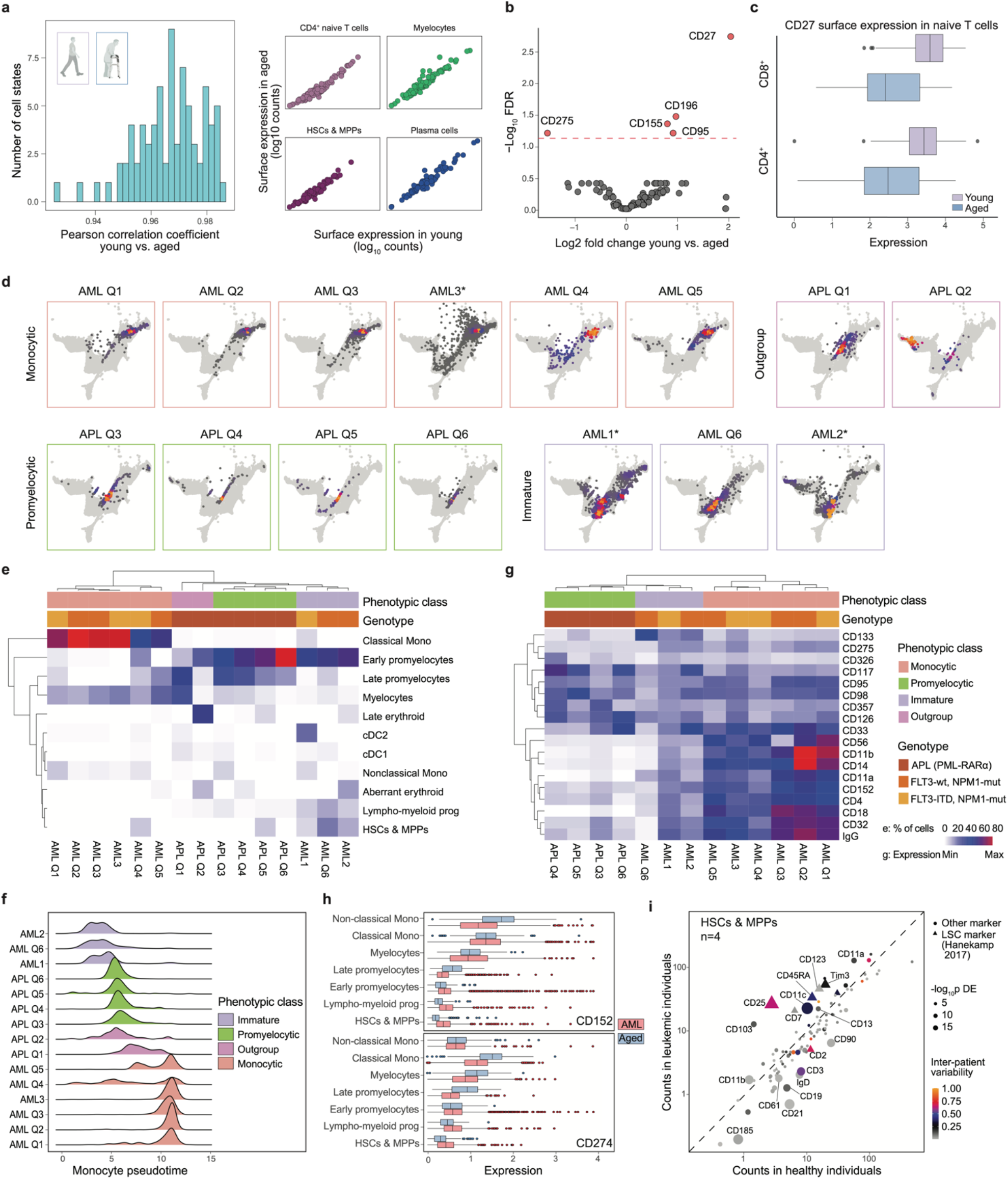
Adaptation of surface protein expression in healthy aging and cancer. **a**. Correlation of surface marker expression between matched cell types from aged and young bone marrow donors. For each cell type, mean surface marker expression across all cells was computed, separately for all ‘young’ and ‘aged’ samples, and the correlation between the two matched cell types was determined. Left panel: Histogram of Pearson correlation coefficients. Right panel: Sample scatter plots depicting the mean surface expression of all measured markers in indicated cell types, see also Supplementary Table 6. **b**. Volcano plot depicting log2 fold change and false discovery rate (FDR) for a test for differential surface marker expression between cells from young and aged individuals, while accounting for cell types as covariates. See Methods, section ‘*Differential expression testing between experimental groups and estimation of inter-patient variability’* for details. **c**. Boxplots depicting CD27 surface expression in naïve T cell populations from young and aged individuals. **d**. Projection of AML samples onto healthy reference. See Supplementary Note 7 for detail. **e**. Clustering of leukemia samples by their projected cell type composition. Lymphoid cells are excluded from the clustering. **f**. Density plots of Monocyte pseudotime, resulting from projection on the healthy reference. See Methods for details. **g**. Heatmap depicting surface markers with differential expression between the phenotypic classes defined in panel *e*. The eight markers with the most significant p values were selected for each comparison between classes. Average expression across all non-lymphoid cells is shown. **h**. Surface expression of immunotherapy targets CTLA-4 (CD152) and PD-L1 (CD274) in different myeloid compartments of healthy donors and AMLs. **i**. Scatter plot depicting the average expression of all surface markers in healthy HSCs & MPPs (x-axis) and leukemic cells projecting to the HSC & MPP cell state (y-axis). Cells from four patients where the HSC/MPP class was covered with more than 20 cells are included (AML1, AML2, AML3 and AML Q6). P-values for differential expression were computed using DESeq2 and are encoded in the symbol size. Inter-patient variability is color-coded, see Methods, section ‘*Differential expression testing between experimental groups and estimation of inter-patient variability*’ for details. See also Supplementary Table 6. For all data shown, bone marrow mononuclear cells from iliac crest aspirations from healthy adult donors or AML/APL patients were used.

We next explored surface marker remodeling in AML, a blood cancer characterized by the accumulation of immature, dysfunctional myeloid progenitors, also called blasts. While the cellular bone marrow of healthy donors displayed highly similar topologies across 6 individuals, initial analysis of 3 AML patients demonstrated that leukemic cells showed patient-specific alterations and a large degree of inter-patient variability (Figure 1b). To develop a generically applicable workflow to interpret data from hematological diseases in the context of our reference, we generated single-cell proteo-genomics datasets from a total of 15 AML patients, covering six t(15;17) translocated acute promyelocytic leukemias (APLs) and nine normal karyotype AMLs with NPM1 mutations, out of which 4 patients carried an additional FLT3 internal tandem duplication (ITD) (Supplementary Table 3). While an unsupervised integration of these data primarily highlighted patient-to-patient variability (Figure S7b), projecting cells onto our healthy reference enabled a fine-mapping of the differentiation stages of leukemia cells (Figures 4d, Supplementary Note 7). Unsupervised clustering of patients based on the relative abundancies of differentiation stages revealed three main categories: ‘monocytic AMLs’ that displayed an extensive accumulation of blasts with classical monocyte phenotype, APLs that were blocked in early and late promyelocyte states, and ‘immature AMLs’ that showed high numbers of immature blasts resembling HSC, MPP, early lympho-myeloid progenitor and early promyelocyte states (Figures 4e-f). In general, leukemic blasts retained many features reminiscent of the cell stage they were blocked in (Figures S7c-e). Accordingly, differential expression analyses revealed that many surface markers which distinguish the different AML states, also mark their corresponding healthy counterparts, such as CD133 for immature AMLs or CD14 and CD11b for monocytic AMLs (Figure 4g). This also translated into differential surface expression of potential drug targets, such as PD-L1 (CD274) and CTLA4 (CD152) (Figure 4h, S7f), suggesting that the myeloid differentiation program of the AML might be essential in the treatment choice of targeted immune therapies.

By contrast, differential analyses between AML and healthy cells from the same differentiation stage revealed markers specifically over-expressed in leukemic cells (Figure 4i, S7c, Supplementary Table 6). Interestingly, these analyses readily identified several previously described leukemia stem cell (LSC) markers, including CD25, Tim-3, CD123 and CD45RA (Hanekamp et al., 2017), supporting the validity of our approach. Quantifying the degree of inter-patient heterogeneity of each marker while accounting for cell state, revealed that many known LSC markers strongly vary in their expression between patients (Figure 4i). Taken together, this workflow of projection to a well-annotated healthy reference in combination with cell-state specific differential expression testing might become a standard in scRNA-seq analyses of hematological diseases. Our computational routines are available online at https://git.embl.de/triana/nrn.

### Data-driven isolation strategies and immunophenotypic characterization of rare bone marrow cell populations

Gating strategies for flow cytometry have evolved historically in a process of trial and error. In particular, the isolation of rare and poorly characterized cell subsets using flow cytometry remains challenging, whereas commonly used gating schemes are not necessarily optimal in purity (precision) and efficiency (recall). To tackle these problems, we explored different machine learning approaches for the data-driven definition of gating schemes. For all populations in our dataset, gating schemes defined by machine learning approaches provided higher precision (purity) if compared to classical gating schemes from literature (Figure 5a, Figure S8, Supplementary Table 7). While different machine learning methods tested achieved similar purities, gates defined by the hypergate algorithm (Becht et al., 2019) offered a higher recall (Figure 5a, Figure S8).

**Figure 5.**
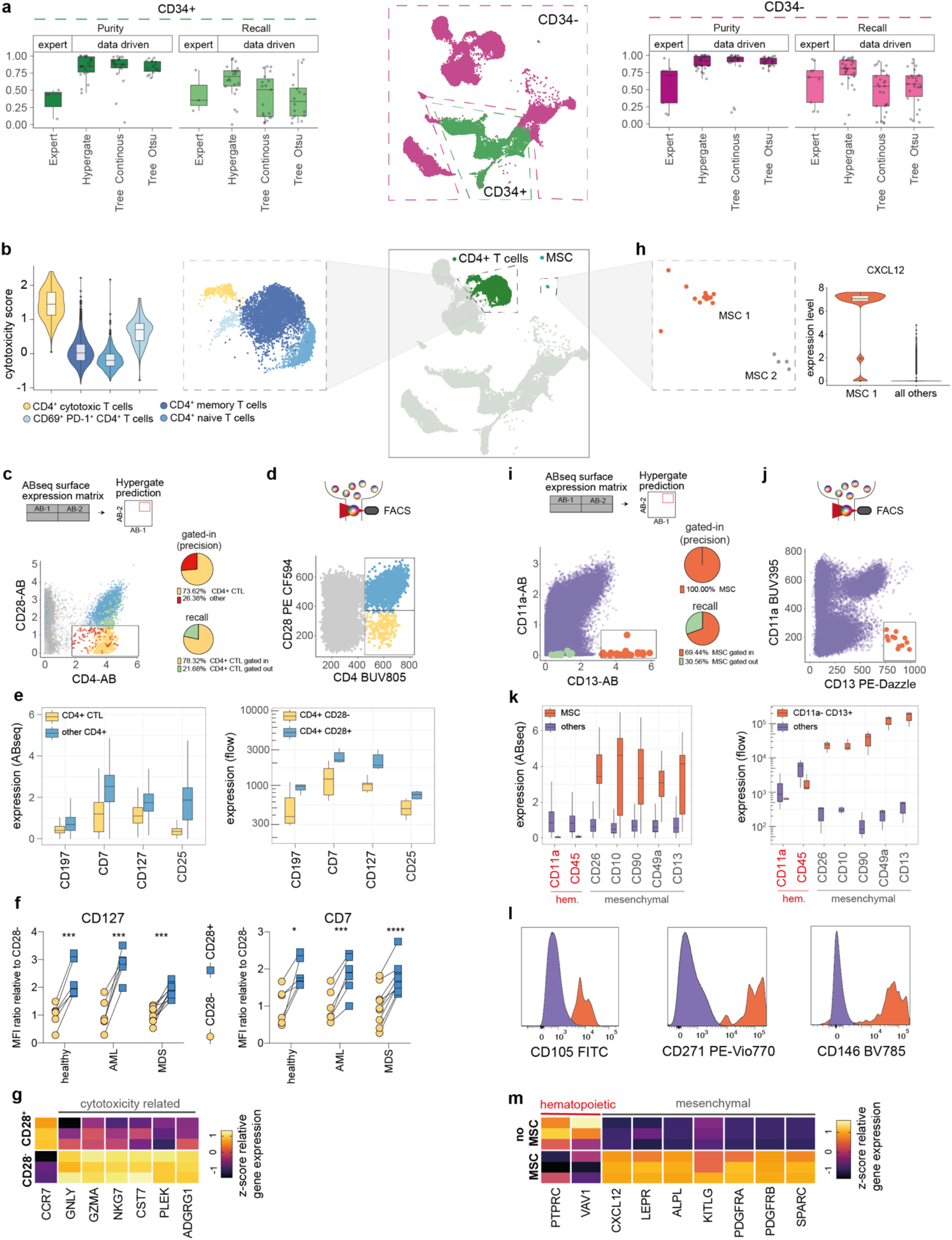
Data-driven definition of gating schemes for rare cell types. **a**. Purity and recall of published expert or data driven gating schemes for cell populations within CD34+ and CD34-compartments. **b**. Different CD4+ T cell subsets are highlighted (central and right panels) and the corresponding distributions of cytotoxicity scores for every subset are displayed (left panel). **c**. Hypergate (Becht et al., 2019) was used to identify a gating scheme for the prospective isolation of cytotoxic CD4+ T cells. The suggested gate is highlighted on a plot depicting the surface protein expression of CD4 and CD28 as identified from pre-gated CD45+CD3+ Abseq data. Yellow and green dots correspond to cytotoxic T cells located within and outside of the selected gate, respectively. Red dots correspond to other cells located inside the selected gate (false positives), blue dots correspond to other CD4+ cells located outside the gate and grey dots to other cells located outside the gate. Pie charts indicate precision and recall. **d**. FACS plot displaying the expression of CD4 and CD28 on pre-gated CD45+CD3+ cells, and respective gates (yellow dots correspond to CD4+CD28-cytotoxic T cells and blue dots to other CD4+CD28+ T cells. **e**. Boxplot depicting the expression of surface markers with differential expression between CD4+ cytotoxic T cells and other CD4+ subsets, as identified from Abseq data (left panel) and validated with FACS using the gating strategy from *d* (right panel). **f**. Paired scatter plot depicting the mean fluorescence intensities (MFI) of CD127 and CD7 in CD4+CD28-cytotoxic CD4+ T cells (yellow) and CD4+CD28+ other CD4+ T cells (blue) in bone marrow samples from healthy, AML and MDS patients. n=6, 6 and 9 patients in the respective groups. **g**. Heatmap depicting gene expression of cytotoxicity-related genes in FACS-sorted CD4+CD28- and CD4+CD28+ cells, as quantified by qPCR (n= 4 patients) **h-k**. Analogous to *b-e*, but with the identification and use of a CD11a-CD13+ gate for the isolation of CXCL12+ mesenchymal stem cells (MSC). Orange and green dots correspond to MSCs located within and outside of the selected gate, respectively. Purple dots correspond to other cells located outside the gate. **l**. Representative FACS histogram plots showing surface expression of well-known MSC surface markers, which were not contained in the original 97 antibody Abseq panel. **m**. Heatmap depicting gene expression of common hematopoietic and MSC signature genes in FACS sorted CD11a-CD13+ MSCs and cells outside the gate, as quantified by qPCR (n= 3 patients). No significance = ns, P<0.05 *, P<0.01 **, P<0.001 ***, P<0.0001 ****. CD4+CD28- and CD4+CD28+ paired cell populations within the same BM donors from different disease entities were compared using paired two-tailed t-test. For all experiments shown, bone marrow mononuclear cells from iliac crest aspirations from healthy adult donors, AML or MDS patients were used.

To validate and demonstrate this approach, we focused on determining novel gating strategies for rare and poorly characterized BM cell types, such as cytotoxic CD4+ T cells (Figure 5b and mesenchymal stem or stromal cells (MSCs) (Figure 5h). Cytotoxic CD4+ T cells represent a rare T cell population characterized by the expression of cytotoxicity genes typically observed in their well-characterized CD8+ T cell counterparts (Szabo et al., 2019). While this cell type has been suggested to be involved in several physiological and pathophysiological processes, no coherent gating strategy for their prospective isolation exists (Takeuchi and Saito, 2017). Hypergate suggested that cytotoxic CD4+ T cells display an immunophenotype of CD4+CD28-, and differential expression analyses of surface markers revealed that cytotoxic CD4+ T cells express significantly lower levels of CD7, CD25, CD127 and CD197 if compared to other CD4+ T cell subsets (Figure 5b-e). Flow cytometric analyses of CD4+CD28-T cells confirmed the expected immunophenotype in BM from healthy donors and patients with different hematological cancers, suggesting a robust and efficient prospective isolation of this rare cell type (Figure 5d-f). Finally, FACS-based sorting of CD4+CD28-T cells followed by gene expression analysis confirmed the expression of cytotoxicity genes in this population (Figure 5g).

MSCs constitute a rare and heterogeneous group of cells in the bone marrow (Al-Sabah et al., 2020; Frenette et al., 2013). While *ex vivo*-expanded MSCs have been phenotyped extensively, primary human MSCs remain poorly characterized, in particular due to their extremely low frequency. In our dataset, we captured a small number of heterogeneous MSCs, with one subset (MSC-1) expressing high levels of the key bone marrow-homing cytokine CXCL12 (Figure 5h). Hypergate suggested CXCL12-expressing MSCs to be most efficiently isolated by expression of CD13 and absence of CD11a (Figure 5i). Indeed, flow cytometric analyses of CD13+CD11a-MSCs validated the immunophenotype suggested by our Abseq data and confirmed known and novel MSC surface markers identified by our approach (Figure 5j-l). Moreover, FACS-based isolation of CD13+CD11a-cells followed by transcriptomic analyses revealed a high enrichment of CXCL12 and other key MSC signature genes (Figure 5m).

Together, these analyses demonstrate the utility of our approach for deriving gating schemes from data and mapping the surface marker expression of poorly characterized populations. The Abseq-App in combination with our single-cell proteo-genomic reference map allows users to define new data-driven gating schemes for any population of interest.

### A fully data-driven gating scheme reflects the molecular routes of human hematopoiesis

Gating schemes for complex biological systems, such as the hematopoietic stem and progenitor cell (HSPC) compartment, are steadily improving. However, there is strong evidence from single-cell transcriptomics (Giladi et al., 2018; Paul et al., 2015; Tusi et al., 2018; Velten et al., 2017), lineage tracing (Perié et al., 2015; Rodriguez-Fraticelli et al., 2018) and single-cell functional experiments (Notta et al., 2016) that even the most advanced gating schemes do not recapitulate the molecular and cellular heterogeneity observed by single-cell genomics approaches. This has contributed to several misconceptions in the understanding of the hematopoietic system, most notably, incorrect assumptions on the purity of cell populations and inconsistent views on lineage commitment hierarchies (Haas et al., 2018; Jacobsen and Nerlov, 2019; Laurenti and Göttgens, 2018; Loughran et al., 2020).

In order to generate flow cytometric gating schemes that most adequately reflect the transcriptomic states associated with hematopoietic stem cell differentiation, we used the Abseq-dataset of CD34+ cells from one BM sample (‘Young1’) to train a decision tree. Thereby, we obtained a gating scheme that uses 12 surface markers to define 14 leaves representing molecularly defined cell states with high precision (Figure 6a-c). The data-derived scheme excelled in the identification of lineage committed progenitors, a major shortcoming of many current gating strategies (Figure 6a-c) (Notta et al., 2016; Paul et al., 2015; Perié et al., 2015; Velten et al., 2017). Importantly, cell populations defined by the data-defined gating scheme were transcriptionally more homogenous, compared to a widely used gating scheme (Figure 6d, e; Doulatov et al., 2010), a state-of-the-art gating scheme focusing on lymph-myeloid differentiation (Figure 6e, S9a-d; Karamitros et al., 2018) and a ‘consensus gating’ scheme generated *in silico* to combine the latter with a scheme focusing on erythroid-myeloid differentiation (Figure 6e, S9b; Psaila et al., 2016). Of note, individual populations from the data-defined scheme displayed a functional output comparable to populations of the ‘consensus gating’ scheme, while the data-defined scheme overall provided a higher level of information on functional lineage commitment (Figure S9e, f).

**Figure 6.**
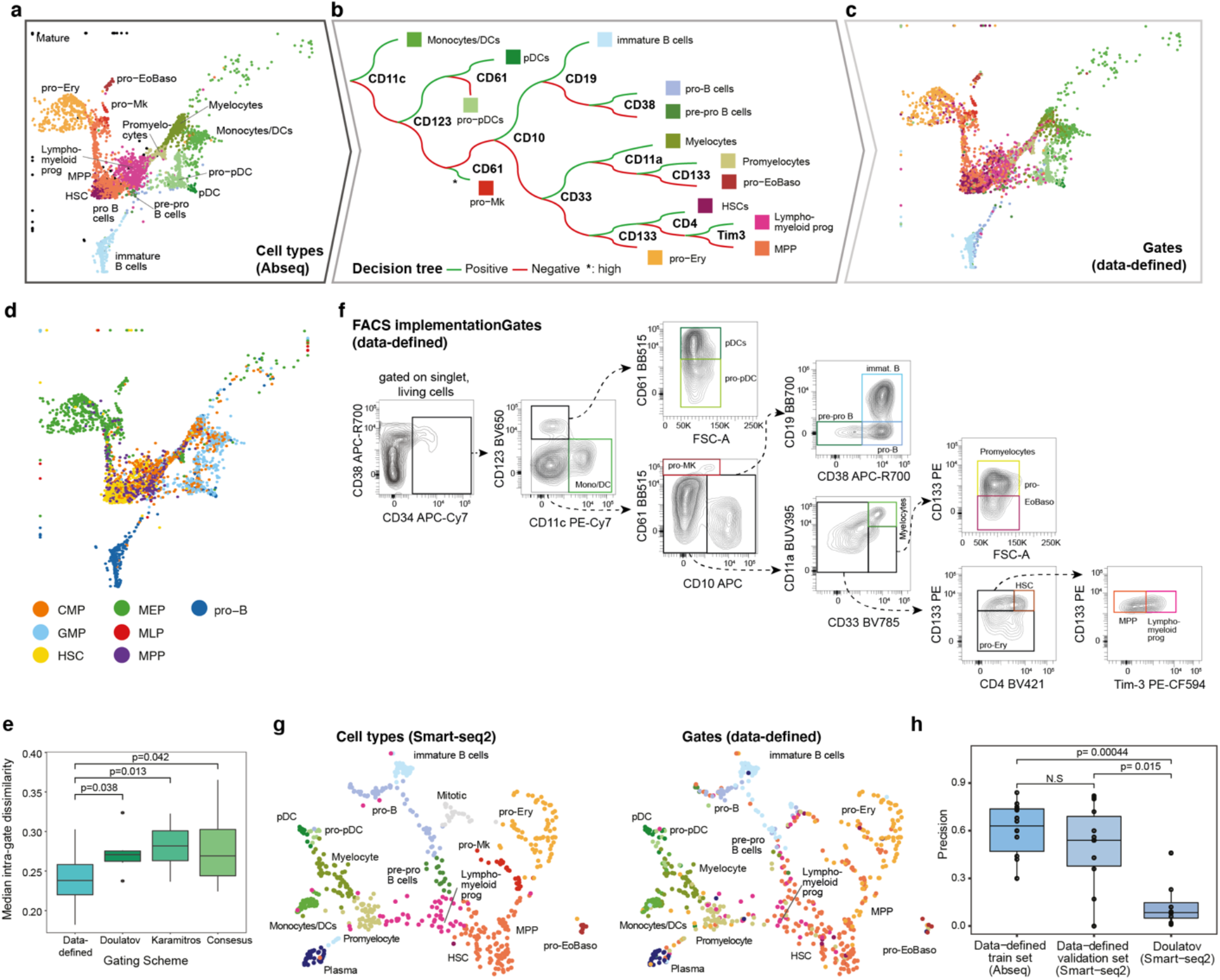
Data-driven definition of gating schemes for hematopoietic stem and progenitor cells. **a**. UMAP depicting all CD34+ HSPCs cells from one healthy young individual. Clustering and cluster annotation were performed exclusively on this individual to achieve a higher subtype resolution of stem cells (‘HSCs’), immature progenitors with lymphoid/myeloid transcriptomic priming (‘Lympho-myeloid progenitros’) and immature progenitors with erythroid/megakaryocytic transcriptomic priming (‘MPP’). See panel *b* for color scheme. **b**. Decision tree using surface marker expression from the Abseq data in order to classify cells into cell types. See Methods, section ‘*Data-driven identification of gating schemes’* and main text for details. **c**. UMAP highlighting cell type classification obtained from the decision tree. Please take note that colors now correspond to putative ‘gates’ applied to the expression levels of the 12 markers shown in panel b, and not to cell types defined from single-cell multi-omics data. **d**. UMAP highlighting classification obtained from a decision tree recapitulating the classical gating scheme used in the field (Doulatov et al., 2010), i.e. HSC: CD34+CD38-CD45RA-CD90+; MPP: CD34+CD38-CD45RA-CD90-; MLP: CD34+CD38-CD45RA+; CMP: CD34+CD38+CD10-CD45RA-*Flt3*+; MEP: CD34+CD38+CD10-CD45RA-*Flt3*-; GMP: CD34+CD38+CD10-CD45RA+*Flt3*+; pro-B: CD34+CD38+CD10+. Since CD135 was not part of the Abseq panel, the expression of *Flt3* was smoothened using MAGIC (van Dijk et al., 2018) for this purpose. Automatic thresholding was used to identify marker-positive cells, see Methods, section ‘*Thresholding of surface marker expression’* for details. **e**. Boxplot depicting the intra-gate dissimilarity for cell classification with panels from Doulatov et al., 2010 (panel *d*), the gating scheme from Karamitros et al., 2018 (*Figure S9*), the in-silico created ‘consensus gating’ scheme combing Doulatov et al., 2010, Karamitros et al., 2018 and Psaila et al., 2016 (*Figure S9*) and the data-driven gating scheme (panel *c*). Intra-gate dissimilarity is defined as one minus the average Pearson correlation of normalized gene and surface antigen expression values of all cells within the gate. P-values are from a two-sided Wilcoxon test. **f**. Implementation of FACS gating scheme suggested by the decision tree from panel *b*. **g**. UMAP display of mRNA expression of n=630 CD34+ HSPCs from an indexed single-cell Smart-seq2 experiment where the expression of the 12 surface markers (for the data-defined gating) was recorded using FACS. Left panel: Clusters are highlighted based on gene expression, see Supplementary Note 8 for details. Right panel: Classification of the cells based on FACS markers using the data-defined gates shown in panel *f*. **h**. Precision of the classification scheme shown in panel *b*, computed on the training data (i.e. the Abseq dataset) and the test data (i.e. the Smart-seq2 dataset). Precision was computed per gate as the fraction of correctly classified cells. For comparison with the Doulatov gating scheme, the dataset from Velten et al., 2017 was used. P-values are from a two-sided Wilcoxon test. For all data shown, bone marrow mononuclear cells from iliac crest aspirations from healthy adult donors were used.

To validate this new gating scheme, we implemented the suggested surface marker panel in a classical flow cytometry setup and performed Smart-seq2 based single-cell RNA-sequencing while simultaneously recording surface marker expression (index-scRNAseq) (Figure 6f, g, Supplementary Note 8). This approach demonstrated that the new gating strategy efficiently separated molecularly defined cell states (Figure 6g). Quantitatively, the data-defined gating scheme performed equally well at resolving molecularly defined cell states on the Abseq training data as on the Smart-seq2 validation data, and significantly outperformed the expert-defined gating scheme (Figure 6h). A limitation of the low cellular throughput of the Smart-seq2 analysis is that the signature-based identification might result in the “over-identification” of certain cell states. Together, our results demonstrate that high-content single-cell proteo-genomic maps can be used to derive data-defined cytometry panels that describe the molecular states of complex biological systems with high accuracy. Moreover, our gating scheme permits a faithful identification and prospective isolation of transcriptomically defined progenitor states in the human hematopoietic hierarchy using cost-effective flow cytometry.

### Systematic integration of single-cell genomics, flow cytometry and functional data via NRN

While classical FACS gating strategies are of great use for the prospective isolation and characterization of populations, single-cell genomics studies revealed that differentiation processes, including the first steps of hematopoiesis, are most accurately represented by a continuous process (Macaulay et al., 2016; Nestorowa et al., 2016; Pellin et al., 2019; Tusi et al., 2018; Velten et al., 2017). To complement the approach based on discrete gates, we here propose that high-dimensional flow cytometry data can be used to place single cells into the continuous space of hematopoietic differentiation spanned by single-cell proteo-genomics exploiting shared surface markers (Figure 7a). Based on the observation that surface marker expressions in flow cytometry and Abseq follow similar distributions (Figure S10a), we developed a new projection algorithm termed nearest rank neighbors (NRN: https://git.embl.de/triana/nrn/, see Methods). Given an identical starting population, NRN employs sample ranks to transform surface marker expression of FACS and Abseq data to the same scale, followed by k-nearest neighbors-based projection into a space defined by the proteo-genomic single-cell data. We tested NRN on FACS indexed Smart-seq2 datasets using the classification panel developed in Figure 6 (12 markers) and a semi-automated panel based on our Abseq data to better resolve erythro-myeloid lineages (11 markers, Supplementary Note 8). We evaluated the performance of NRN using a variety of methods. First, cell types molecularly defined by Smart-seq2 were placed correctly on the Abseq UMAP (Figure 7b). For most molecularly defined cell types, the accuracy of the projection using the flow cytometry data was close to the performance of data integration using whole transcriptome data with a state-of-the-art algorithm (Figure S10b-d). Most importantly, the projections closely reflected the gradual progression of cells through pseudotime, as confirmed by the expression dynamics of key lineage genes from our FACS indexed Smart-seq2 data (Figure 7c). This suggests that NRN, in combination with high quality reference datasets, can be used to study the continuous nature of cellular differentiation processes by flow cytometry.

**Figure 7.**
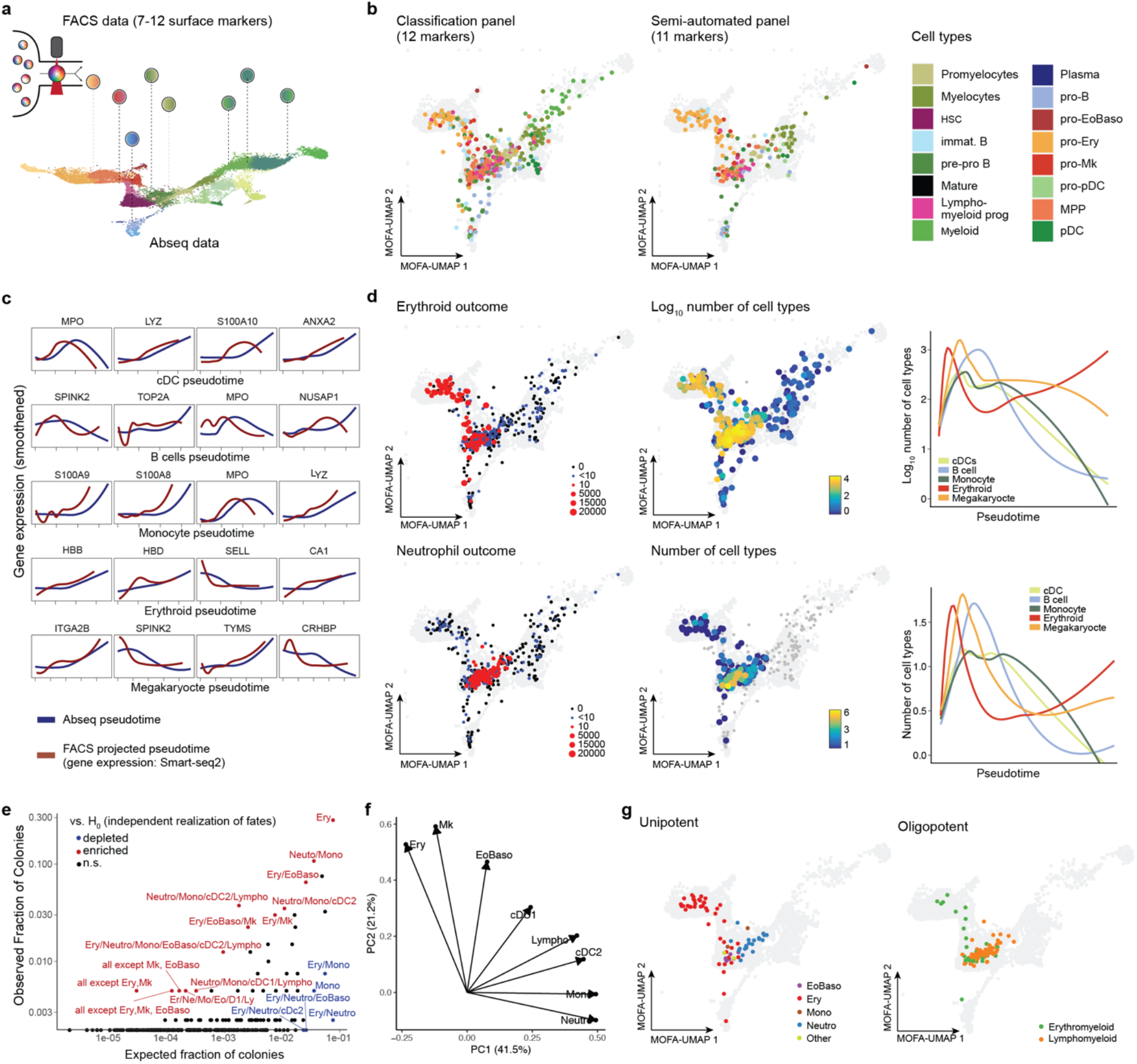
Systematic integration of single-cell genomics, flow cytometry and functional data via NRN. **a**. Illustration of the concept. See main text and methods for details. **b**. Projection of indexed Smart-seq2 data onto a reference UMAP. Single cells with recorded (‘indexed’) FACS measurements of surface markers (data-defined classification panel or semi-automated panel) were subjected to Smart-Seq2 based scRNA-seq. The commonly used surface markers were used to project cells via NRN onto the Abseq UMAP (see Methods, section ‘*The NRN algorithm for integrating FACS and single cell genomics data’* for details). Take note that only FACS data was used for the projection in UMAP space, whereas colors depict cell types identified from RNA expression. **c**. Projection of indexed Smart-seq2 data onto reference pseudotime trajectories. The same single cells were projected onto the differentiation trajectories shown in Figure 3a using FACS measurements only. The expression of differentiation markers was then determined from available Smart-seq2 data and smoothened over projected pseudotime values (red lines). For comparison, the expression values of the same genes were determined from Abseq data and smoothened over the reference pseudotime values (blue lines). The selected genes correspond to the five genes with the strongest statistical association to the respective trajectory. **d**. Projection of indexed single-cell culture data onto a reference UMAP. Single cells with available FACS measurements of 12 surface markers (data-defined classification panel from Figure 5) were projected onto the UMAP defined by Abseq via NRN. Single cells were seeded into culture medium supporting the formation of erythroid, megakaryocytic and distinct myeloid cell types, see Methods, section *‘Single-cell index cultures’* for details. The ability of single cells to give rise to erythroid cells and neutrophils were highlighted on the UMAPs. Colony size as well as the total number of cell types per colony are highlighted both on the UMAP and on projected pseudotime. **e**. Analysis of cell type combinations in n=397 colonies. For any combination of Erythroid (Ery), Neutrophil (Neutro), Monocytic (Mono), Eosinophil or Basophil (EoBaso), Lymphoid (Lympho), Megakaryocytic (Mk) and Dendritic (cDC1 and cDC2) potential, the scatter plot depicts the fraction of colonies containing this exact combination of cell types (y-axis) and the theoretical fraction of colonies containing this exact combination of cell types under the assumption that cell fates are independently realized with the same marginal probabilities (x-axis). Significance was calculated from a binomial test and is color-coded. These analyses do not exclude that other combinations of fates are not biologically selected as well, i.e. absence of evidence does not constitute evidence for absence. **f**. PCA analysis of colony compositions. **g**. Distribution of colonies with frequent combinations of cells types in the projected UMAP space. Erythromyeloid: Only containing EoBaso, Mk and/or Ery cells. Lymphomyeloid: All other combinations. For all data shown, bone marrow mononuclear cells from iliac crest aspirations from healthy adult donors were used.

A key limitation of single-cell genomics remains the lack of insights into functional differentiation capacities of cells. We therefore evaluated whether NRN can be used to interpret functional single-cell data in the context of single-cell genomic reference maps. For this purpose, we performed single-cell culture assays, while recording surface markers of our data-defined gating scheme from Figure 6, followed by data integration using our Abseq data via NRN. As expected, cells with the highest proliferative capacity and lineage potency were placed in the phenotypic HSC and MPP compartments, and HSPCs placed along the transcriptomically defined differentiation trajectories continuously increased the relative generation of cells of the respective lineage (Figure 7d). Functionally unipotent progenitors cells were observed along the respective transcriptomic trajectories, but were also present in the phenotypic HSC/MPP compartment (Figure 7d, g), in line with previous findings on early lineage commitment of HSPCs (Notta et al., 2016; Paul et al., 2015; Velten et al., 2017). In contrast, oligopotent cells with distinct combinations of cell fates were specifically enriched in the HSC/MPP compartment (Figure 7d, g). Some of these fate combinations, in particular combinations of erythroid, megakaryocytic and eosinophilic/basophilic fates, and combinations of lymphoid, neutrophilic, monocytic, and dendritic fates, co-occurred more frequently than expected by chance (Figure 7e, f), in line with most recent findings on routes of lineage segregation (Drissen et al., 2019; Görgens et al., 2014; Tusi et al., 2018; Velten et al., 2017). Despite strong associations between surface phenotype, transcriptome and function, cells with a highly similar phenotype can give rise to different combinations of lineages (Figure 7g). This observation suggests a role of stochasticity in the process of lineage commitment, or hints towards layers of cell fate regulation not observed in the transcriptome. Taken together, our observations confirm that hematopoietic lineage commitment predominantly occurs continuously along the routes predicted by the transcriptome, with an early primary erythro-myeloid versus lympho-myeloid split (Drissen et al., 2019; Görgens et al., 2014; Notta et al., 2016; Paul et al., 2015; Tusi et al., 2018; Velten et al., 2017) and might help reconciling discrepancies in the interpretation of previous studies.

In sum, our data resource alongside the NRN algorithm enables accurate integration of flow data with single-cell genomics data. This permits the charting of continuous processes by flow cytometry and the mapping of single-cell functional data into the single-cell genomics space.

## DISCUSSION

In this study, we have demonstrated the power of single-cell proteo-genomic reference maps for the design and analysis of cytometry experiments. We have introduced a map of human blood and bone marrow spanning the expression of 97-197 surface markers across 45 cell types and stages of hematopoietic stem cell differentiation, healthy ageing, and leukemia. Our dataset is carefully annotated and will serve as a key resource for hematology and immunology.

While cytometry experiments remain the working horse of immunology, stem cell biology and hematology, recent single-cell atlas projects have revealed that current cytometry setups do not accurately reflect the full complexity of biological systems (Papalexi and Satija, 2018; Paul et al., 2015). For the first time, we have exploited single-cell proteo-genomic data to systematically design and interpret flow cytometry experiments that mirror most accurately the cellular heterogeneity observed by single-cell transcriptomics. Unlike approaches based on index sorting (Baron et al., 2019; Paul et al., 2015; Velten et al., 2017; Wilson et al., 2008), single-cell proteo-genomics has a sufficient throughput to enable the profiling of entire tissues or organs, and at the same time covers up to several hundred of surface markers. Unlike single-cell RNA-seq data, antibody tag counts reflect the true distributions of surface marker expression, enabling a quantitative integration of cell atlas data with FACS. Building on these unique properties of our reference map, we have automated the design of gating schemes for the isolation of rare cell types, we have devised a gating strategy that reflects the molecular routes of hematopoietic stem cell differentiation, and we have demonstrated the direct interpretation of flow cytometry data in the context of our reference.

These advances enable a functional characterization of molecularly defined cell states and thereby directly impact on hematopoietic stem cell research. There is a growing consensus in the field that lineage commitment occurs early from primed HSCs, that not all progenitor cells in the classical MEP/GMP gates are functionally oligopotent, and that the main branches of the hematopoietic system are a GATA2-positive branch of erythroid, megakaryocytic and eosinophil/basophil/mast cell progenitors, as well as a GATA2-negative branch of lympho-myeloid progenitors, including monocytes, neutrophils and dendritic cells (Drissen et al., 2019; Giladi et al., 2018; Görgens et al., 2014; Pellin et al., 2019; Tusi et al., 2018; Velten et al., 2017; Zheng et al., 2018). Due to a lack of better alternatives, many functional studies still use the classical gating scheme alongside the outdated concept of ‘common myeloid progenitors’(Akashi et al., 2000; Kondo et al., 1997; Pei et al., 2017). Here, we introduce and validate a flow cytometry scheme that allows the prospective isolation of molecularly homogeneous progenitor populations. We have used this scheme to show that transcriptional lineage priming impacts on cellular fate *in vitro* (Notta et al., 2016; Velten et al., 2017), thereby contributing further evidence for the revised model of hematopoiesis. In the future, a wider use of this scheme has the potential to avoid conflicting results stemming from imprecisely defined populations.

Furthermore, these advances enable the rapid profiling of blood formation and other bone marrow phenotypes while offering a resolution comparable to single-cell genomics. Recently, bone marrow phenotypes of diseases, ranging from sickle cell disease (Hua et al., 2019) to leukemia (van Galen et al., 2019) have been investigated using scRNA-seq. However, due to economic and experimental hurdles, the throughput of these studies has remained restricted to maximally tens of patients. Accordingly, the ability to associate patient genotypes with phenotypes is thereby highly limited, and these assays have not been translated to diagnostic routines. Our new gating schemes and analytical strategies are widely applicable to profile aberrations encountered in disease, both in research, and ultimately in clinical diagnostics.

While we have demonstrated the implementation of data-driven design and analysis strategies for cytometry assays in the context of bone marrow, conceptually the approach presented here can be applied to any organ of interest. Thereby, it has the potential to enable the precise isolation and routine profiling of the myriad of cell types discovered by recent single-cell atlas projects.

## SUPPLEMENTAL MATERIAL

- Methods
- Supplemental Tables S1-S9
- Supplemental Figures S1-S10
- Supplemental Video S1
- Supplementary Notes 1-8

## ACKNOWLEDGEMENTS

We thank Vadir Lopez-Salmeron, Vishnu Ramani, Edyta Kowalczyk and Wieland Keilholz from BD Biosciences / Multiomics for providing oligo-labelled antibodies and their support in the implementation of the Rhapsody platform. We would like to thank members of the Haas, Velten, Trumpp and Steinmetz labs for helpful discussions. Moreover, we thank members from the DKFZ flow cytometry and the EMBL genomics core facility for support. This work was financially supported by the Emerson foundation grant 643577 (to LV), grant PID2019-108082GA-I00 by the Spanish Ministry of Science, Innovation and Universities (MCIU/AEI/FEDER, UE), the German Bundesministerium für Bildung und Forschung (BMBF) through the Juniorverbund in der Systemmedizin ‘LeukoSyStem’ (FKZ 01ZX1911D to LV, SH and SR), SFB873, FOR2674 and FOR2033 funded by the Deutsche Forschungsgemeinschaft (DFG), the SyTASC consortium (Deutsche Krebshilfe), the Dietmar Hopp Foundation (all to AT); and the José Carreras Foundation for Leukemia Research (grant no. DCJLS 20R/2017 to LV, AT, and SH). LV acknowledges support of the Spanish Ministry of Science and Innovation to the EMBL partnership, the Centro de Excelencia Severo Ochoa and ‘the CERCA Programme / Generalitat de Catalunya. DN is an endowed professor of the Deutsche José 641Carreras Leukämie Stiftung (DJCLS H 03/01). Contributions by DN, JCJ, WKH and TB were supported by the Gutermuth Foundation, the H.W. & J. Hector fund, Baden-Württemberg.

## AUTHOR CONTRIBUTIONS

S.H., L.V. and M.P. conceived the study with help from D.H., A.T., and V.B.. D.V., S.T. and M.P. performed the single-cell proteo-genomics experiments with help from D.L. and V.B.. D.V. performed the experimental validations, established new experimental gating schemes and performed functional experiments with help from M.A. and P.H-M.. S.T., L.J-S. and L.V. performed bioinformatics analyses with conceptional input from D.V., M.P. and S.H.. S.T. developed the AbSeq-App. S.T. and L.V. established the NRN algorithm. S.H. supervised the experimental work with conceptional input from L.V.. L.V. supervised the bioinformatics analyses with conceptional input from S.H. T.A. co-supervised S.T.. M.P., D.O-R. and B.R. provided assistance in cell sorting and single-cell work-flows. S.R., R.L., T.B., J-C.J, D.N., W-K.H. and C.M-T. provided clinical samples and conceptional input on data interpretation. S.H., L.V., S.T., L.J-S. and D.V. wrote the manuscript and prepared figures. All authors have carefully read the manuscript.

## CONFLICT OF INTEREST

The oligo-coupled antibodies used in this study were a gift from BD Biosciences. The authors declare no other relevant conflicts of interest.

## CODE AVAILABILITY

The implementation of the NRN algorithm is available at https://git.embl.de/triana/nrn

## DATA AVAILABILITY

Data is available for interactive browsing at https://abseqapp.shiny.embl.de. Datasets including raw and integrated gene expression data, cell type annotation, metadata and dimensionality reduction are available as Seurat v3 objects through figshare: https://figshare.com/projects/Single-cell_proteo-genomic_reference_maps_of_the_human_hematopoietic_system/94469

## SUPPLEMENTARY FIGURES

**Figure S1.**
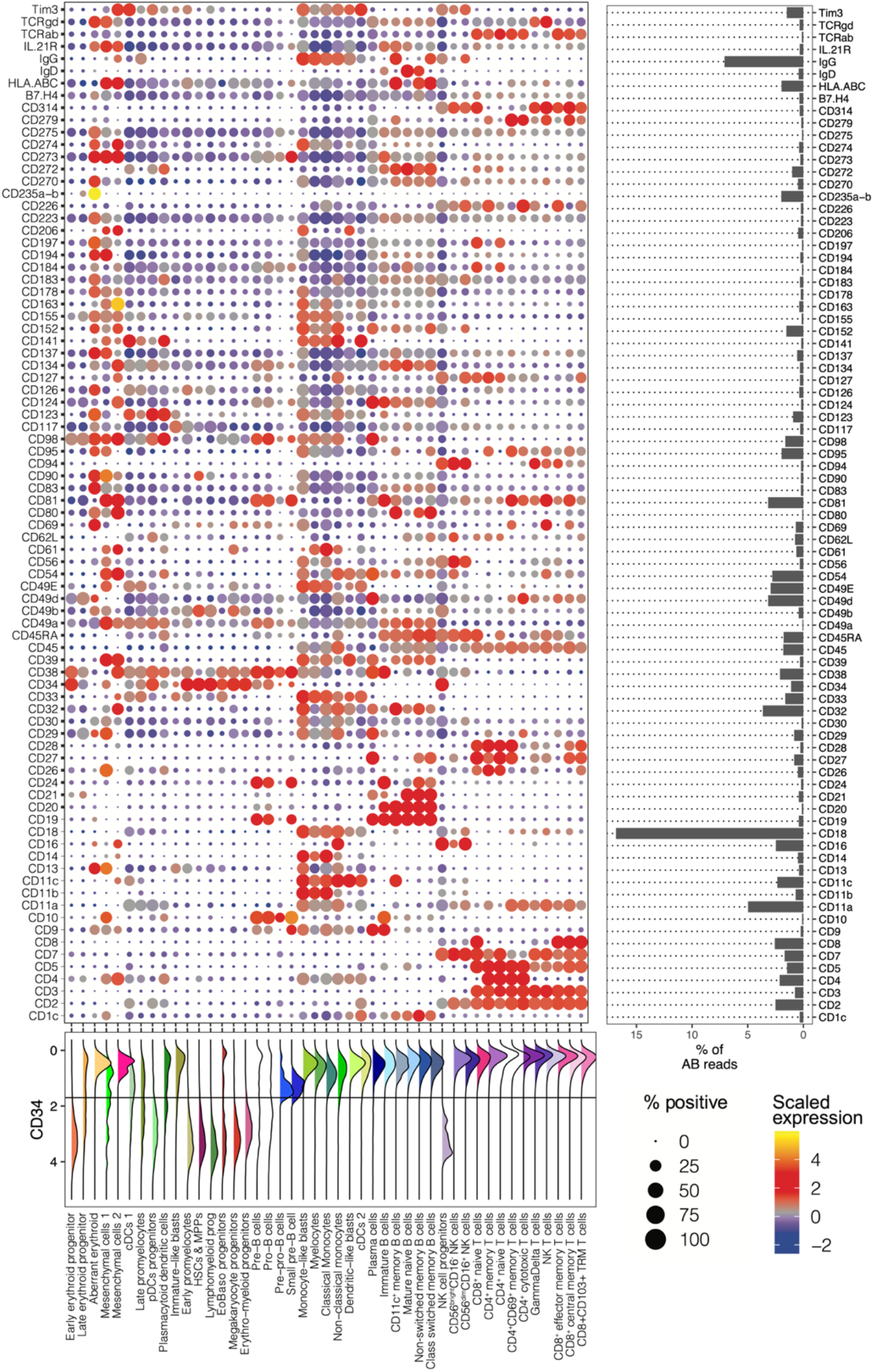
A proteo-genomic single-cell map of 97 surface markers in human bone marrow. Related to *Figure 1*. Dot plot depicting the expression of all surface markers by cell type. Color indicates mean normalized expression, point size indicates the fraction of cells positive for the marker. Automatic thresholding was used to identify positive cells, see Methods, section ‘*Thresholding of surface marker expression’* for details. The panel on the right depicts the fraction of total reads obtained for each marker as a proxy for absolute expression levels. Bottom panel illustrates the distribution of CD34+ expression across populations, similar plots can be generated for any marker using the Abseq-App.

**Figure S2.**
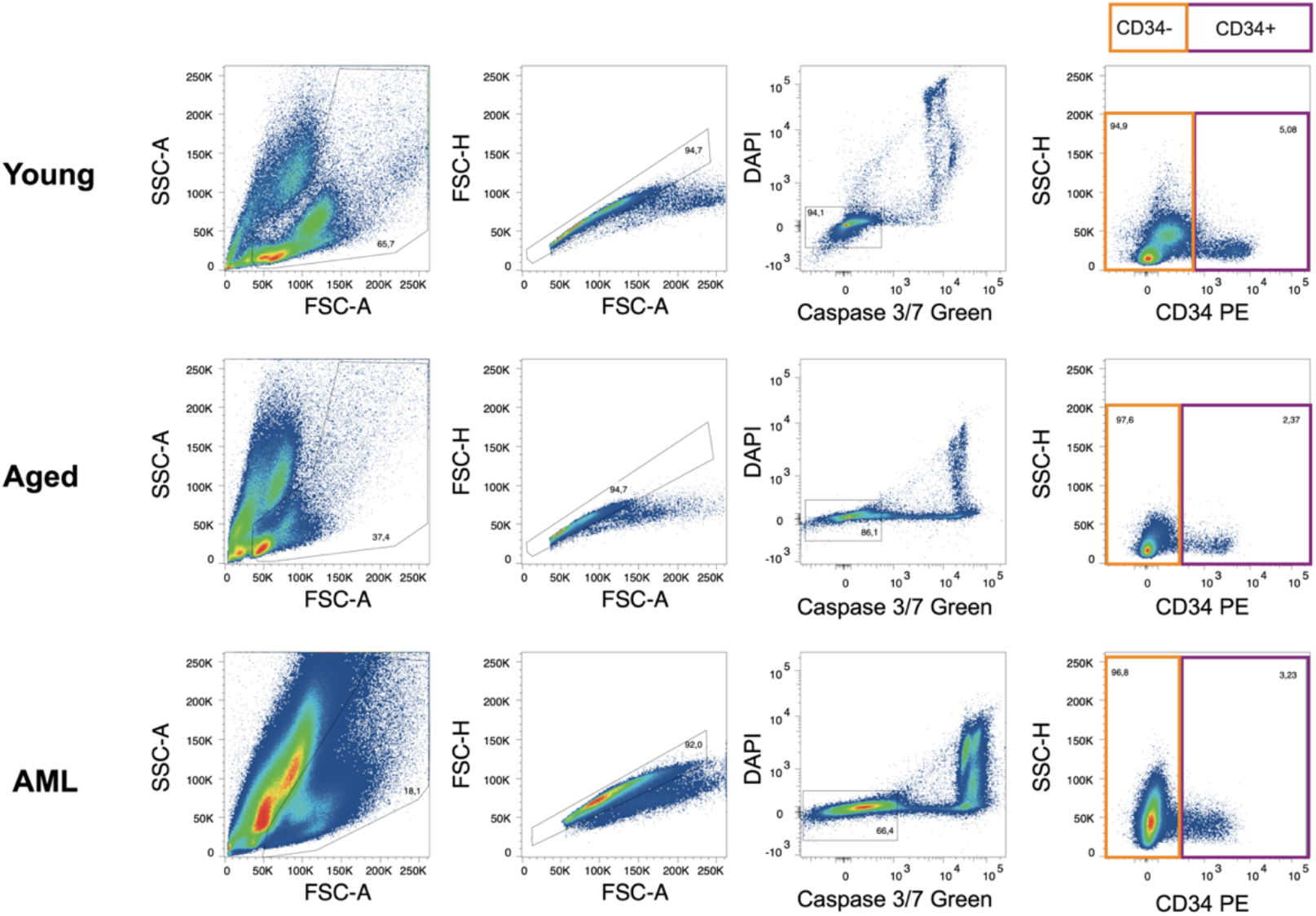
Representative gating schemes used for the enrichment of CD34+ cells. Related to *Figure 1*. For additional information on cell sorting setups, see Methods, section *‘Cell sorting for Abseq’*.

**Figure S3.**
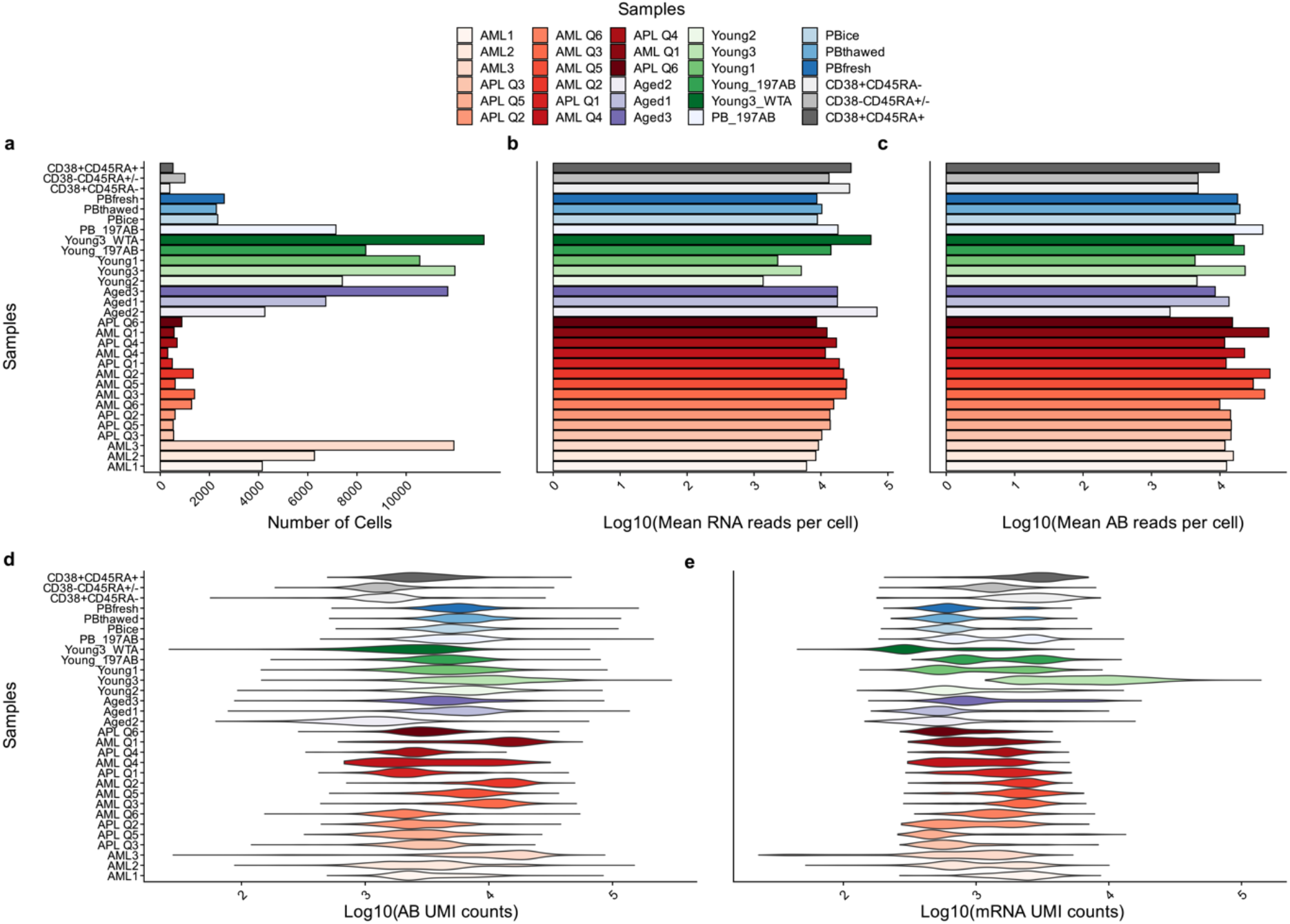
Sequencing statistics. Related to *Figure 1*. Plots depict **a**. the number of cells passing filters. Note that samples AML Q1-Q6 and APQ1-6 were multiplexed (hashed) into one experiment. **b-c**. the sequencing depth on the surface and mRNA level and **d-e**. the number of surface and mRNA molecules per cell observed. Note that targeted mRNA sequencing was performed as described in the main text.

**Figure S4.**
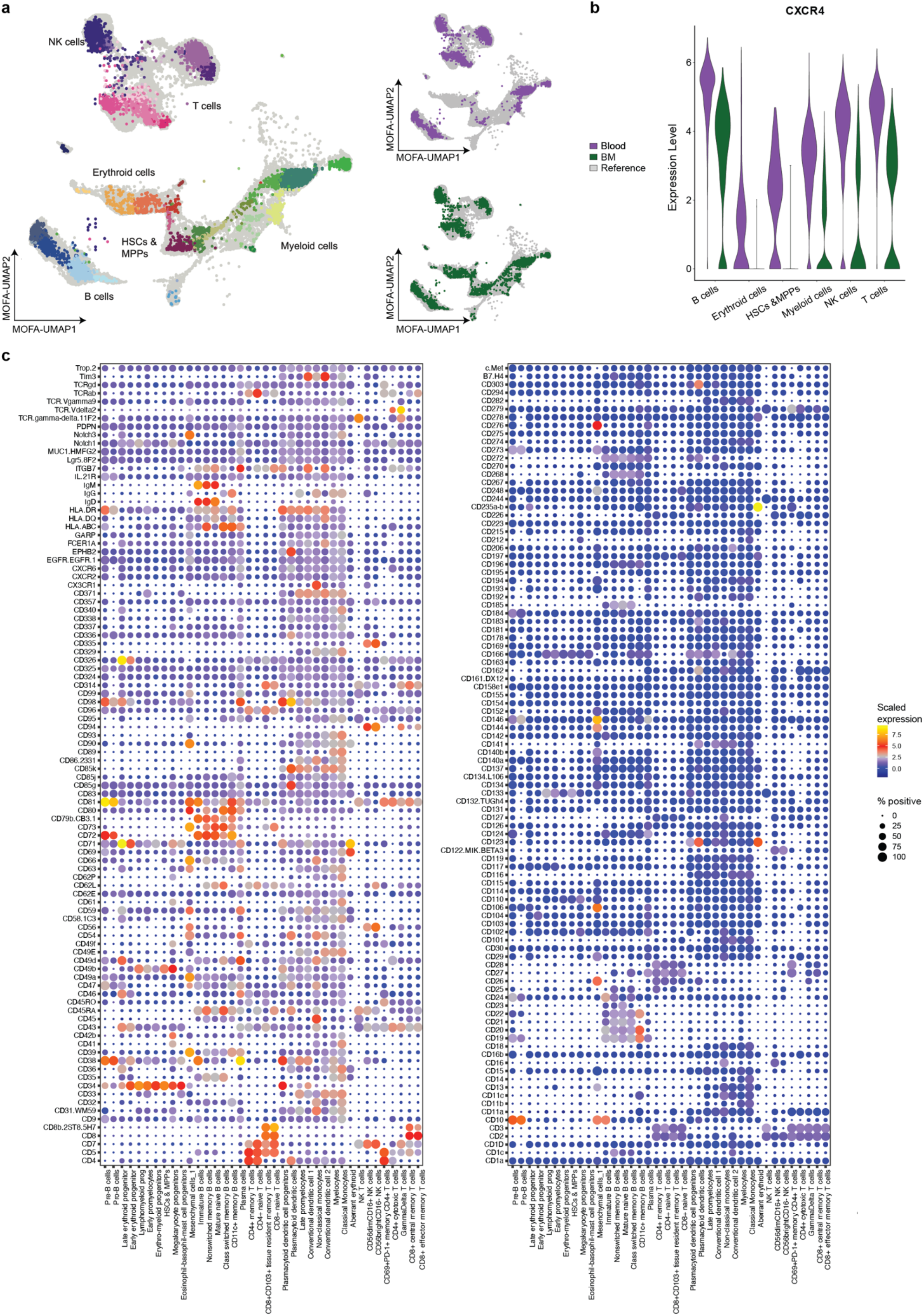
A single-cell proteo-genomic map of 197 surface markers in human bone marrow and blood. Related to *Figure 1*. **a**. UMAP projection on the original coordinate system from the healthy dataset (see *Supplementary Note 7*). Cells are colored by the mapped cell type. **b**. UMAP colored by sample origin (blood and bone marrow). **c**. Violin plot depicting the expression of the bone marrow homing receptor CXCR4 on matching cell types of the blood and bone marrow. **d**. Dot plot depicting the expression of all surface markers by cell type. Color indicates mean normalized expression, point size indicates the fraction of cells positive for the marker. Automatic thresholding was used to identify positive cells, see Methods, section ‘*Thresholding of surface marker expression’* for detail. For all data shown, bone marrow mononuclear cells from iliac crest aspirations or peripheral blood mononuclear cells from healthy adult donors were used.

**Figure S5.**
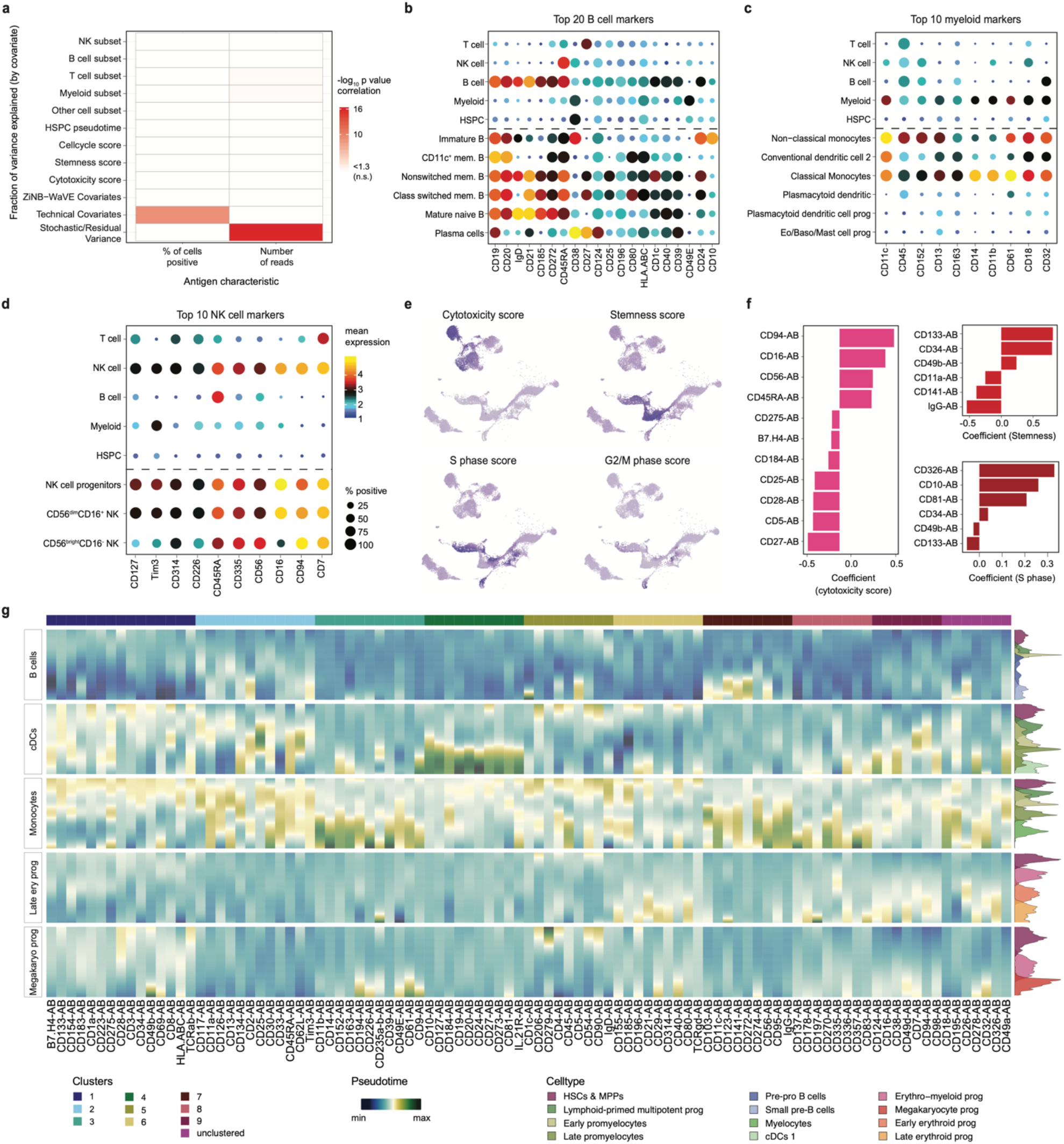
Markers of cell types and biological processes. Related to *Figure 2*. **a**. Heatmap investigating if the fraction of variance explained by the different covariates is correlated to antigen-level technical covariates. P values were calculated from Pearson correlation using a t-distribution. **b-d**. Dot plot depicting the expression of the 10-20 surface markers with the highest fraction of variance explained by B cell subtype *(b)*, myeloid subtype *(c)* and NK cell subtype *(d)*. Color indicates mean normalized expression, point size indicates the fraction of cells positive for the marker. Automatic thresholding was used to identify positive cells, see Methods, section ‘*Thresholding of surface marker expression’* for details. **e**. UMAPs highlighting the scores for various biological processes, as computed using the gene lists from Supplementary Table 9. **f**. Bar charts depicting the markers with the highest fraction of variance explained by cytotoxicity score (pink), stemness score (red) and S-phase score (dark red), and the corresponding model coefficients. See *Supplementary Table 9* for the gene lists used for calculating these scores. **g**. Pseudotime of all 97 surface proteins for the five trajectories (B cells, cDCs, Monocytes, Late erythroid progenitor and Megakaryocyte progenitor). Markers were clustered according to their expression pattern using tradeseq (van den Berge, 2020). The density plots indicate the differentiation stages along the pseudotime.

**Figure S6.**
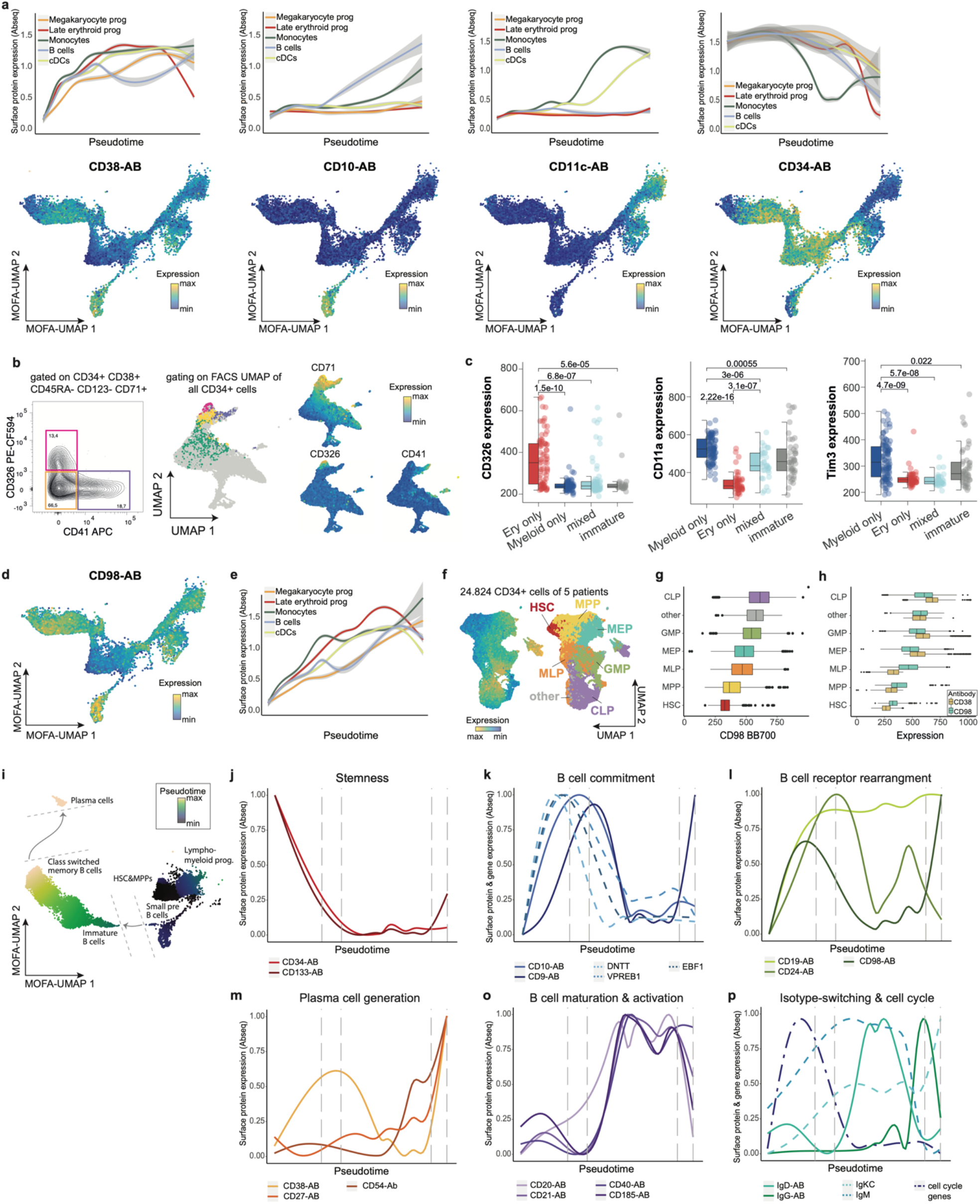
Surface markers associated with HSC and B cell differentiation. Related to *Figure 2* and *3*. **a**. top panels: Line plots depicting normalized CD38, CD10, CD11c and CD34 surface protein expression (Abseq data) smoothened over the different pseudotime trajectories illustrated in *Figure 3a*. bottom panels: UMAP display of CD34+ HSPCs, highlighting the surface expression of each corresponding marker. **b**. Left panel: gating strategy for subsetting CD71+ erythroid/megakaryocytic HSPCs into CD41+ megakaryocyte progenitors and CD326+ erythroid progenitors. Right panel: UMAP display of CD34+ cells from a healthy donor analyzed with a 12-color FACS panel focused erythroid/megakaryocytic differentiation (see *Supplemental Table S8*). Surface expression values were used as input for UMAP dimensionality reduction. Feature plots of CD71, CD326 and CD41 expression highlight the bifurcation within CD71+ HSPCs. **c**. Culture outcome categories described in Figure 3g were analyzed with regards to their CD326, CD11a or Tim3 surface expression. Wilcoxon rank sum test was used for comparison of individual groups and significance levels between groups are depicted. **d**-**e**. Like *Figure 3 d-e*, except that CD98 expression is shown. **f**. UMAP display of CD34+ cells from five healthy donors analyzed with a 12-color FACS stem and progenitor panel (see Supplemental Table S8). Surface expression values were used as input for UMAP dimensionality reduction. Left panel shows CD98 surface expression, right panel shows assignment of individual gates to the UMAP according to the following gating strategy; HSC: CD34+CD38-CD45RA-CD90+; MPP: CD34+CD38-CD45RA-CD90-; MLP: CD34+CD38-CD45RA+; MEP: CD34+CD38+CD10-CD45RA-; GMP: CD34+CD38+CD10-CD45RA+; CLP: CD34+CD38+CD10+CD45RA+; other: cells that did not fall into any of the mentioned gates. **g**. Boxplots showing CD98 expression in individual cell populations mentioned in *f*. **h**. Boxplots showing co-expression of CD98 and CD38 surface markers in respective cell populations. **i**. Like Figure 3a, UMAP plot depicting the pseudotime score along the B cell differentiation trajectory emanating from CD34+ HSCs & MPPs and Lympho-myeloid progenitors. **j-p**. Line plots depicting surface protein expression (Abseq data) representative for different indicated biological processes smoothened over the B cell pseudotime trajectory. For all experiments shown, human adult bone marrow mononuclear cells from iliac crest aspirations were used.

**Figure S7.**
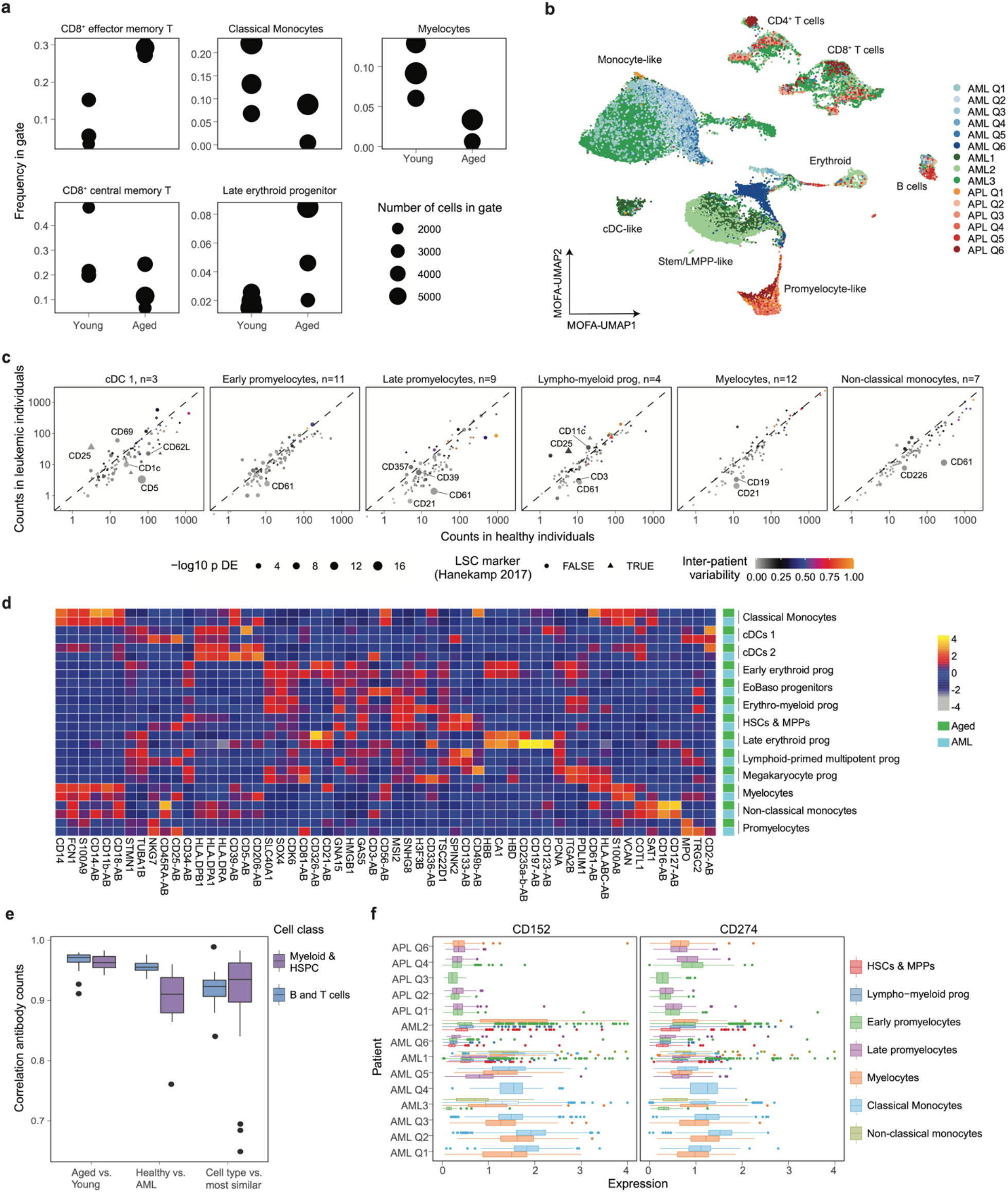
Changes in surface protein expression and cell type abundance induced by ageing and leukemia. Related to *Figure 4*. **a**. Frequency of selected cell types in young and aged individuals. Only the cell types with the most significant changes are shown, see Methods, section ‘*Changes in cell type abundance between experimental groups*’. **b**. UMAP display of all AML patients. Data were integrated using scanorama and MOFA, as for the main dataset (see Method ‘Data analysis of Abseq data’ and ‘MOFA integration, Clustering, and identification of cell type markers’). **c**. For every myeloid cell state with sufficient representation of at least 20 cells in at least three patients, surface marker expression in AML (x-axis) is compared to surface marker expression in healthy individuals (y-axis). AML cell types were defined using a projection as in main *Figure 4d, e*. P-values for differential expression were computed using DESeq2 and are encoded in the symbol size. Inter-patient variability is color-coded, see Methods, section ‘*Differential expression testing between experimental groups and estimation of inter-patient variability’* for details (n=indicates the number of patients included). See also Supplementary Table 6. **d**. Heatmap depicting cell state specific gene expression in leukemic and healthy individuals. Five most significantly overexpressed markers were identified for each cell state, using only leukemic cells. The expression of all markers selected is shown and compared to their expression in the corresponding healthy cell states. **e**. Correlation in surface marker expression between cells from aged, young and leukemic individuals, similar to main *Figure 4a*. Correlations are shown for matching cell types from young versus aged individuals, from healthy individuals versus AML patients, as well as for cell types versus the transcriptomically most similar cell type available in the dataset. **f**. Boxplot depicting the expression of CD152 and CD274 in different cell states from different patients. Only populations covered with at least 50 cells in a given patient are included. See also main *Figure 4h*. For all data shown, bone marrow mononuclear cells from iliac crest aspirations from healthy adult donors or AML/APL patients were used.

**Figure S8.**
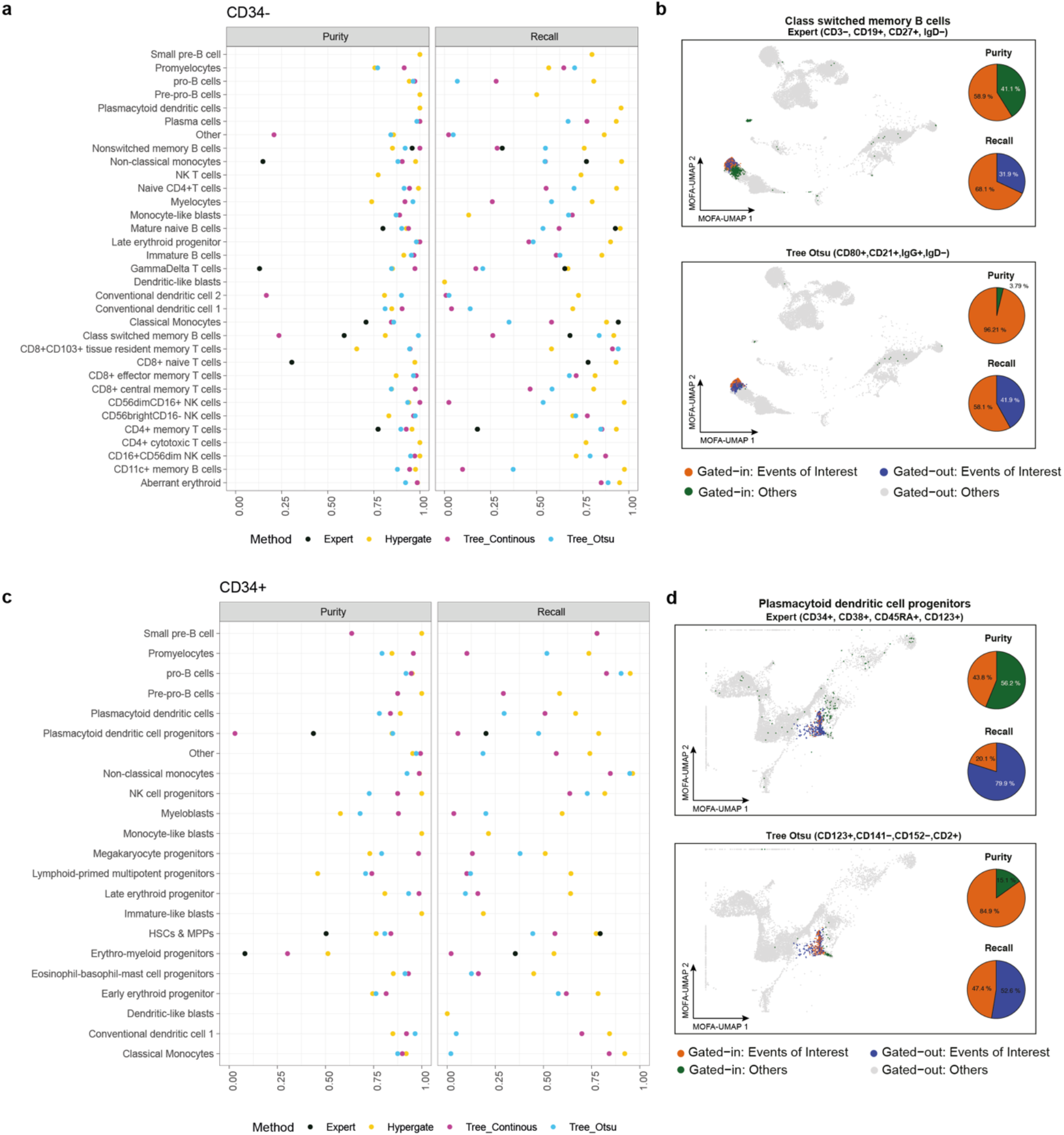
Comparison of data-defined and state-of-the-art (expert-defined) gating schemes. Related to *Figure 5*. **a**. Performance of different methods used for the definition of gates of CD34-populations. Gates for each cell type were defined from CD34-Abseq data using the following strategies: Black dots correspond to gates that were manually set by an expert based on the current state of the art for purifying the cell type of interest (*Supplementary Table 7*). Yellow dots correspond to gate that were set using the hypergate algorithm (Becht et al., 2019). Violet dots correspond to gates that were set using a decision tree. light-blue dots correspond to gates that were set using a decision tree with pre-defined thresholds, see Methods, section ‘*Data-driven identification of gating schemes’*. For each gating scheme, precision (purity) and recall were calculated. **b**. Illustration of the calculation of precision (purity) and recall for class switched memory B cells. Orange and blue dots on the UMAP correspond to class switched memory B cells located within and outside of the selected gate, respectively. Green dots correspond to other cells located inside the selected gate (false positives) and grey dots to other cells located outside the gate (true negatives). Pie charts indicate precision (purity) and recall. Top panel: An expert defined state of the art gating scheme (CD3-CD19+CD27+IgD-) is shown. Bottom panel: A data defined gating scheme (CD80+CD21+IgG+IgD-) is shown. **c**. Like *a*, except that CD34+ populations are shown. **d**. Like *b*, except that gating schemes to define plasmacytoid dendritic cell progenitors are shown.

**Figure S9:**
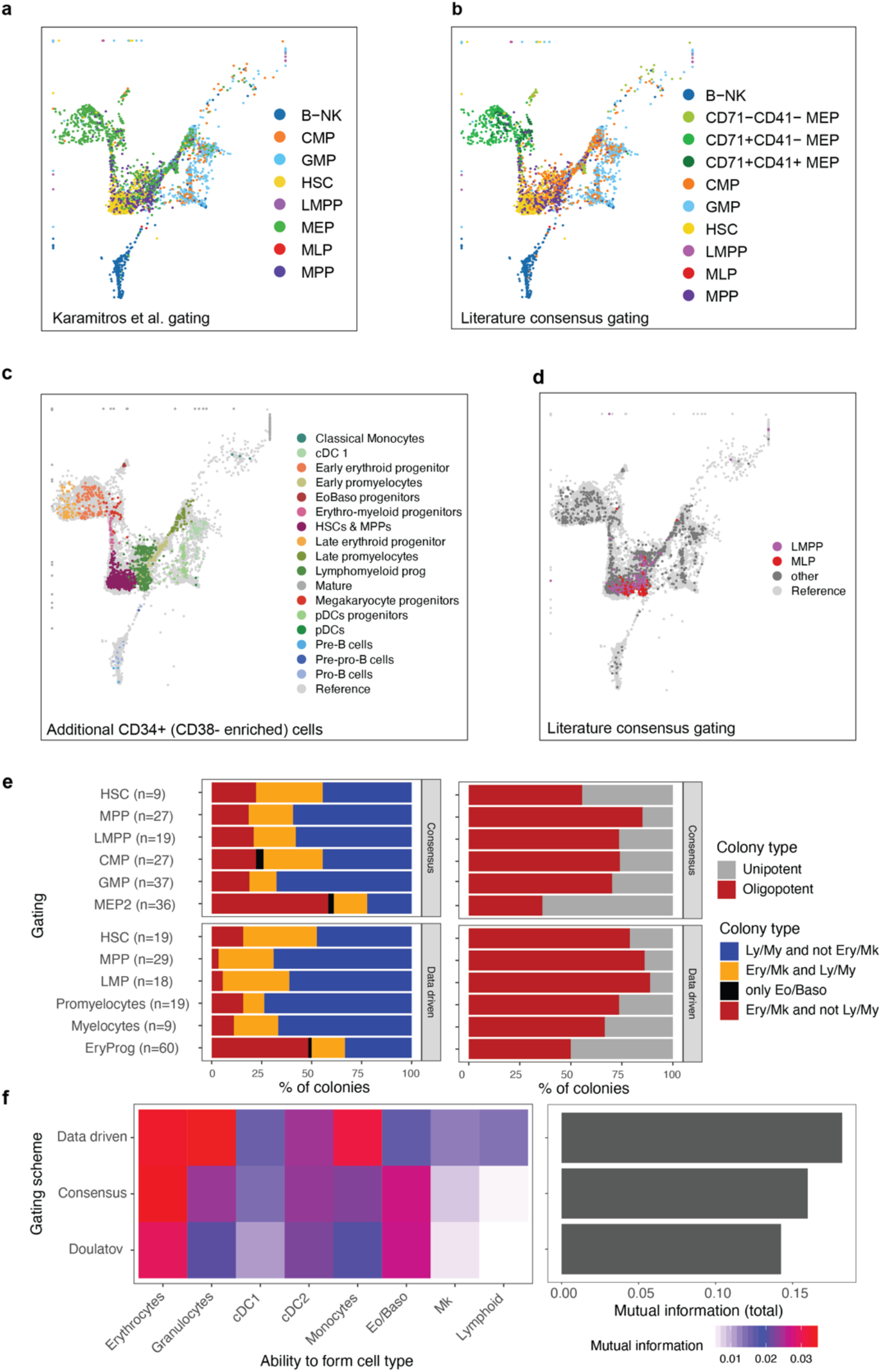
Evaluation of different gating schemes. Related to *Figure 6*. **a**. UMAP highlighting classification obtained from the gating scheme described by Karamitros et al., 2018, i.e. HSC: CD34+CD38-CD10-CD45RA-CD90+; MPP: CD34+CD38-CD10-CD45RA-CD90-; LMPP:CD34+CD38-CD10-CD45RA+; MLP: CD34+CD38-CD10+; MEP: CD34+CD38+CD10-CD45RA-CD123-; CMP: CD34+CD38+CD10-CD45RA-CD123+; GMP: CD34+CD38+CD10-CD45RA+CD123+; B-NK: CD34+CD38+CD10+. **b**. UMAP highlighting classification obtained from a consensus scheme combining the schemes of Doulatov et al., Karamitros et al. and Psaila et al., HSC: CD34+CD38-CD10-CD45RA-CD90+; MPP:CD34+CD38-CD10-CD45RA-CD90-; LMPP:CD34+CD38-CD10-CD45RA+; MLP: CD34+CD38-CD10+; CD71-CD41-MEP: CD34+CD38+CD10-CD45RA-*FLT3*-*ITGA2B*-*TFRC-*; CD71+CD41-MEP: CD34+CD38+CD10-CD45RA-*FLT3*-*ITGA2B*-*TFRC*+; CD71+CD41+ MEP: CD34+CD38+CD10-CD45RA-*FLT3*-*ITGA2B*+; CMP: CD34+CD38+CD10-CD45RA-*FLT3*+; GMP: CD34+CD38+CD10-CD45RA+; B-NK: CD34+CD38+CD10+. The marker CD135, CD41, CD71 were not part of the 97 Abseq panel. The expression of the corresponding genes, *FLT3, ITGA2B* and *TFRC*, were smoothened using MAGIC respectively (van Dijk et al., 2018). **c**. UMAP of additional CD34+ cells with specific enrichment of CD34+ CD38-cells, projected on the original coordinate system, colored by mapped cell types **d**. Same as *c* but colored by immunophenotypic classification obtained from a consensus scheme recapitulating the scheme of Karamitros et al. and and Psaila et al. (see above). **e**. Separation of functional potential by the data driven and the literature ‘consensus gating’ scheme. Single cells were sorted according to the two gating schemes and cultured for 19 days. Colonies were scored as Ery/Mk if they contained at least 5 erythrtoid or megakaryocytic cells, and as Ly/My if they contained at least 5 cells of types Neutrophil, cDC, Monocyte, or B/NK. Unipotent: Only one of these cell types was formed with at least 5 cells; oligopotent: At least two of these cell types were formed. Only gates for which at least 9 colonies were observed are shown. **f**. Mutual information (in nats) between the gate identity and the ability to form any of the cell types, or the total mutual information across all cell types.

**Figure S10.**
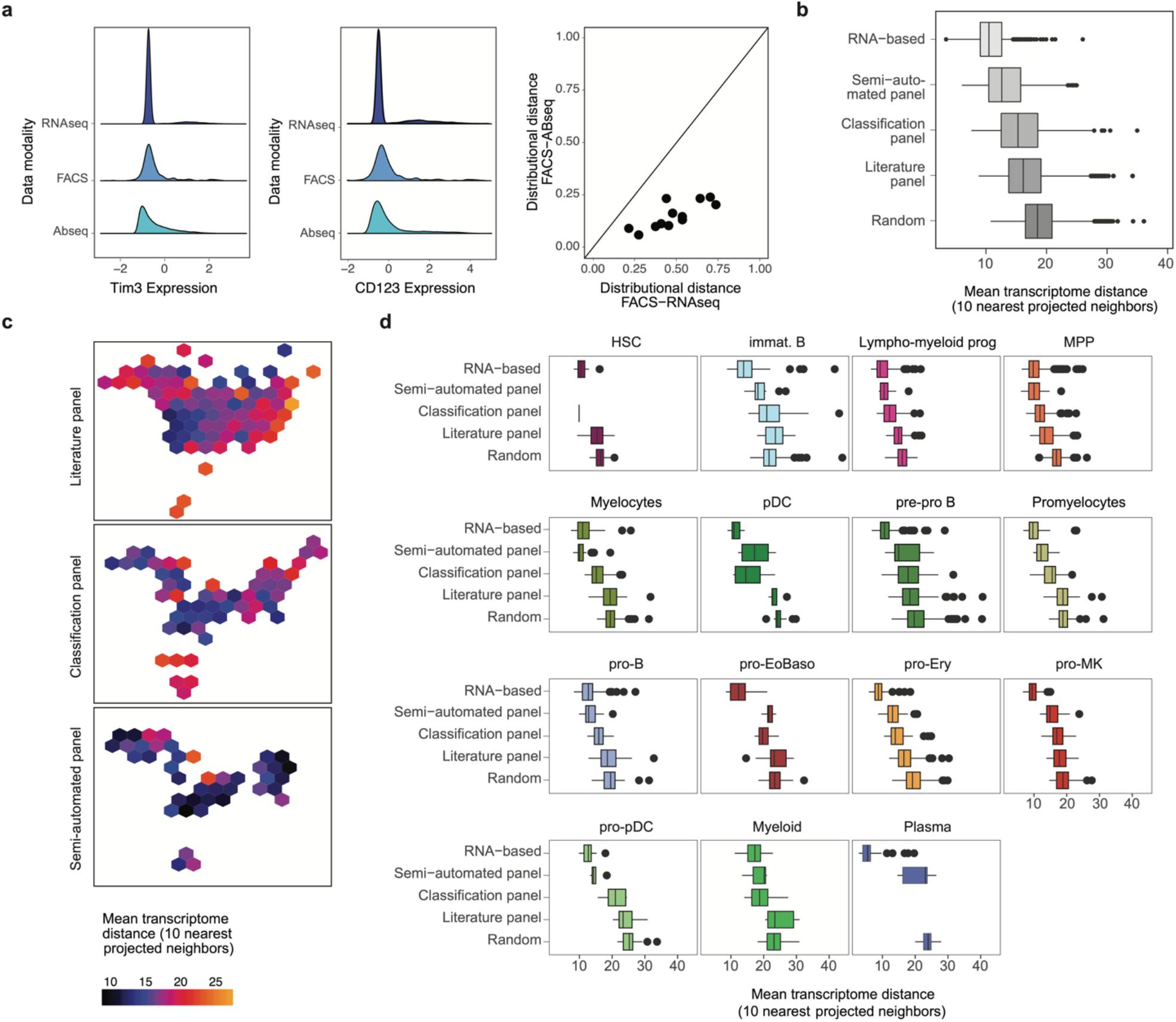
Projection and classification of cytometry data using a single-cell proteo-genomic reference. Related to *Figure 7*. **a**. Distribution of normalized, scaled expression values of Tim3 (left panel) and CD123 (central panel) measured by scRNA-seq, Abseq, and FACS. Right panel: Scatter plot depicts the dissimilarity between the distribution of expression values measured by FACS, and the distribution measured by scRNA-seq (x-axis) or Abseq (y-axis) as quantified using Kolmogorov-Smirnov distance. Data for all markers included in the panel from main *Figure 6f* is shown. **b-d**. Comparison of data integration strategies. Smart-seq2 data and Abseq data were integrated with five different strategies. RNA-based: Integration by Seurat v3, based on gene expression (transcriptome). Random: Random selection of ten nearest neighbors. Others: Surface marker-based integration using NRN, using defined sets of surface markers (Classification panel, Semi-automated panel: see *Table S8*. Literature panel: CD34, CD38, CD45RA, CD90, CD10, CD135/*Flt3*, CD49f.). For every cell projected on the UMAP, the ten nearest neighbors in projected UMAP space were identified. Subsequently, the mean Euclidean distance between their location in a gene expression-based PCA space (Smart-seq2) was computed. **b**. Boxplot summarizing the distance across data integration strategies. **c**. Hexagonal plot summarizing the projection accuracy for different regions of the UMAP. **d**. Boxplots stratified by cell type demonstrate that projection using the semi-automated panel performs close to an RNA-based integration in most cases.

## Notes

https://abseqapp.shiny.embl.de

https://figshare.com/projects/Single-cell_proteo-genomic_reference_maps_of_the_human_hematopoietic_system/94469

### Competing Interest Statement

The oligo-coupled antibodies used in this study were a gift from BD Biosciences. Besides this, the authors declare no relevant conflicts of interest.

### Summary of Updates

Dataset has been updated to include a broader representation of AML patients (now, total of 15 patients). A reference-based analysis strategy for AML patient data is introduced. Figure 4 has been completely revised accordingly. More specific analyses on functional potential were added to figure 3, 7 and the supplement.

## REFERENCES

Akashi, K., Traver, D., Miyamoto, T., and Weissman, I.L. (2000). A clonogenic common myeloid progenitor that gives rise to all myeloid lineages. Nature 404, 193–197.

Al-Sabah, J., Baccin, C., and Haas, S. (2020). Single-cell and spatial transcriptomics approaches of the bone marrow microenvironment. Curr. Opin. Oncol. 32, 146–153.

Baccin, C., Al-Sabah, J., Velten, L., Helbling, P.M., Grünschläger, F., Hernández-Malmierca, P., Nombela-Arrieta, C., Steinmetz, L.M., Trumpp, A., and Haas, S. (2020). Combined single-cell and spatial transcriptomics reveal the molecular, cellular and spatial bone marrow niche organization. Nat. Cell Biol. 22, 38–48.

Baron, C.S., Barve, A., Muraro, M.J., van der Linden, R., Dharmadhikari, G., Lyubimova, A., de Koning, E.J.P., and van Oudenaarden, A. (2019). Cell Type Purification by Single-Cell Transcriptome-Trained Sorting. Cell 179, 527-542.e19.

Becht, E., Simoni, Y., Coustan-Smith, E., Evrard, M., Cheng, Y., Ng, L.G., Campana, D., and Newell, E.W. (2019). Reverse-engineering flow-cytometry gating strategies for phenotypic labelling and high-performance cell sorting. Bioinformatics 35, 301–308.

van Dijk, D., Sharma, R., Nainys, J., Yim, K., Kathail, P., Carr, A.J., Burdziak, C., Moon, K.R., Chaffer, C.L., Pattabiraman, D., et al. (2018). Recovering Gene Interactions from Single-Cell Data Using Data Diffusion. Cell 174, 716-729.e27.

Van Dongen, J.J.M., Lhermitte, L., Böttcher, S., Almeida, J., Van Der Velden, V.H.J., Flores-Montero, J., Rawstron, A., Asnafi, V., Lécrevisse, Q., Lucio, P., et al. (2012). EuroFlow antibody panels for standardized n-dimensional flow cytometric immunophenotyping of normal, reactive and malignant leukocytes. Leukemia 26, 1908–1975.

Doulatov, S., Notta, F., Eppert, K., Nguyen, L.T., Ohashi, P.S., and Dick, J.E. (2010). Revised map of the human progenitor hierarchy shows the origin of macrophages and dendritic cells in early lymphoid development. Nat. Immunol. 11, 585–593.

Drissen, R., Thongjuea, S., Theilgaard-Mönch, K., and Nerlov, C. (2019). Identification of two distinct pathways of human myelopoiesis. Sci. Immunol. 4.

Fagnoni, F.F., Vescovini, R., Mazzola, M., Bologna, G., Nigro, E., Lavagetto, G., Franceschi, C., Passeri, M., and Sansoni, P. (1996). Expansion of cytotoxic CD8+ CD28-T cells in healthy ageing people, including centenarians. Immunology 88, 501–507.

Frenette, P.S., Pinho, S., Lucas, D., and Scheiermann, C. (2013). Mesenchymal stem cell: Keystone of the hematopoietic stem cell niche and a stepping-stone for regenerative medicine. Annu. Rev. Immunol. 31, 285–316.

van Galen, P., Hovestadt, V., Wadsworth, M.H., Hughes, T.K., Griffin, G.K., Battaglia, S., Verga, J.A., Stephansky, J., Pastika, T.J., Lombardi Story, J., et al. (2019). Single-Cell RNA-Seq Reveals AML Hierarchies Relevant to Disease Progression and Immunity. Cell 176, 1265-1281.e24.

Giladi, A., and Amit, I. (2018). Single-Cell Genomics: A Stepping Stone for Future Immunology Discoveries. Cell 172, 14–21.

Giladi, A., Paul, F., Herzog, Y., Lubling, Y., Weiner, A., Yofe, I., Jaitin, D., Cabezas-Wallscheid, N., Dress, R., Ginhoux, F., et al. (2018). Single-cell characterization of haematopoietic progenitors and their trajectories in homeostasis and perturbed haematopoiesis. Nat. Cell Biol. 20, 836–846.

Görgens, A., Ludwig, A.K., Möllmann, M., Krawczyk, A., Dürig, J., Hanenberg, H., Horn, P.A., and Giebel, B. (2014). Multipotent hematopoietic progenitors divide asymmetrically to create progenitors of the lymphomyeloid and erythromyeloid lineages. Stem Cell Reports 3, 1058–1072.

Haas, S. (2020). Hematopoietic Stem Cells in Health and Disease—Insights from Single-Cell Multi-omic Approaches. Curr. Stem Cell Reports 6, 67–76.

Haas, S., Trumpp, A., and Milsom, M.D. (2018). Causes and Consequences of Hematopoietic Stem Cell Heterogeneity. Cell Stem Cell 22, 627–638.

Han, X., Wang, R., Zhou, Y., Fei, L., Sun, H., Lai, S., Saadatpour, A., Zhou, Z., Chen, H., Ye, F., et al. (2018). Mapping the Mouse Cell Atlas by Microwell-Seq. Cell 172, 1091-1107.e17.

Han, X., Zhou, Z., Fei, L., Sun, H., Wang, R., Chen, Y., Chen, H., Wang, J., Tang, H., Ge, W., et al. (2020). Construction of a human cell landscape at single-cell level. Nature 581, 303–309.

Hanekamp, D., Cloos, J., and Schuurhuis, G.J. (2017). Leukemic stem cells: identification and clinical application. Int. J. Hematol. 105, 549–557.

Hua, P., Roy, N., de la Fuente, J., Wang, G., Thongjuea, S., Clark, K., Roy, A., Psaila, B., Ashley, N., Harrington, Y., et al. (2019). Single-cell analysis of bone marrow–derived CD34+ cells from children with sickle cell disease and thalassemia. Blood 134, 2111–2115.

Jacobsen, S.E.W., and Nerlov, C. (2019). Haematopoiesis in the era of advanced single-cell technologies. Nat. Cell Biol. 21, 2–8.

Karamitros, D., Stoilova, B., Aboukhalil, Z., Hamey, F., Reinisch, A., Samitsch, M., Quek, L., Otto, G., Repapi, E., Doondeea, J., et al. (2018). Single-cell analysis reveals the continuum of human lympho-myeloid progenitor cells article. Nat. Immunol. 19, 85–97.

Kondo, M., Weissman, I.L., and Akashi, K. (1997). Identification of clonogenic common lymphoid progenitors in mouse bone marrow. Cell 91, 661–672.

Laurenti, E., and Göttgens, B. (2018). From haematopoietic stem cells to complex differentiation landscapes. Nature 553, 418–426.

Loughran, S.J., Haas, S., Wilkinson, A.C., Klein, A.M., and Brand, M. (2020). Lineage commitment of hematopoietic stem cells and progenitors: insights from recent single cell and lineage tracing technologies. Exp. Hematol. 88, 1–6.

Macaulay, I.C., Svensson, V., Labalette, C., Ferreira, L., Hamey, F., Voet, T., Teichmann, S.A., and Cvejic, A. (2016). Single-Cell RNA-Sequencing Reveals a Continuous Spectrum of Differentiation in Hematopoietic Cells. Cell Rep. 14, 966–977.

Nestorowa, S., Hamey, F.K., Pijuan Sala, B., Diamanti, E., Shepherd, M., Laurenti, E., Wilson, N.K., Kent, D.G., and Göttgens, B. (2016). A single-cell resolution map of mouse hematopoietic stem and progenitor cell differentiation. Blood 128, e20–e31.

Notta, F., Zandi, S., Takayama, N., Dobson, S., Gan, O.I., Wilson, G., Kaufmann, K.B., McLeod, J., Laurenti, E., Dunant, C.F., et al. (2016). Distinct routes of lineage development reshape the human blood hierarchy across ontogeny. Science (80-.). 351.

Papalexi, E., and Satija, R. (2018). Single-cell RNA sequencing to explore immune cell heterogeneity. Nat. Rev. Immunol. 18, 35–45.

Paul, F., Arkin, Y., Giladi, A., Jaitin, D.A., Kenigsberg, E., Keren-Shaul, H., Winter, D., Lara-Astiaso, D., Gury, M., Weiner, A., et al. (2015). Transcriptional Heterogeneity and Lineage Commitment in Myeloid Progenitors. Cell 163, 1663–1677.

Pei, W., Feyerabend, T.B., Rössler, J., Wang, X., Postrach, D., Busch, K., Rode, I., Klapproth, K., Dietlein, N., Quedenau, C., et al. (2017). Polylox barcoding reveals haematopoietic stem cell fates realized in vivo. Nature 548, 456–460.

Pellin, D., Loperfido, M., Baricordi, C., Wolock, S.L., Montepeloso, A., Weinberg, O.K., Biffi, A., Klein, A.M., and Biasco, L. (2019). A comprehensive single cell transcriptional landscape of human hematopoietic progenitors. Nat. Commun. 10.

Perié, L., Duffy, K.R., Kok, L., De Boer, R.J., and Schumacher, T.N. (2015). The Branching Point in Erythro-Myeloid Differentiation. Cell 163, 1655–1662.

Peters, M.J., Joehanes, R., Pilling, L.C., Schurmann, C., Conneely, K.N., Powell, J., Reinmaa, E., Sutphin, G.L., Zhernakova, A., Schramm, K., et al. (2015). The transcriptional landscape of age in human peripheral blood. Nat. Commun. 6.

Psaila, B., Barkas, N., Iskander, D., Roy, A., Anderson, S., Ashley, N., Caputo, V.S., Lichtenberg, J., Loaiza, S., Bodine, D.M., et al. (2016). Single-cell profiling of human megakaryocyte-erythroid progenitors identifies distinct megakaryocyte and erythroid differentiation pathways. Genome Biol. 17.

Rodriguez-Fraticelli, A.E., Wolock, S.L., Weinreb, C.S., Panero, R., Patel, S.H., Jankovic, M., Sun, J., Calogero, R.A., Klein, A.M., and Camargo, F.D. (2018). Clonal analysis of lineage fate in native haematopoiesis. Nature 553, 212–216.

Schaum, N., Karkanias, J., Neff, N.F., May, A.P., Quake, S.R., Wyss-Coray, T., Darmanis, S., Batson, J., Botvinnik, O., Chen, M.B., et al. (2018). Single-cell transcriptomics of 20 mouse organs creates a Tabula Muris. Nature 562, 367–372.

Shahi, P., Kim, S.C., Haliburton, J.R., Gartner, Z.J., and Abate, A.R. (2017). Abseq: Ultrahigh-throughput single cell protein profiling with droplet microfluidic barcoding. Sci. Rep. 7, 1–12.

Stoeckius, M., Hafemeister, C., Stephenson, W., Houck-Loomis, B., Chattopadhyay, P.K., Swerdlow, H., Satija, R., and Smibert, P. (2017). Simultaneous epitope and transcriptome measurement in single cells. Nat. Methods 14, 865–868.

Stuart, T., and Satija, R. (2019). Integrative single-cell analysis. Nat. Rev. Genet. 20, 257–272.

Szabo, P.A., Levitin, H.M., Miron, M., Snyder, M.E., Senda, T., Yuan, J., Cheng, Y.L., Bush, E.C., Dogra, P., Thapa, P., et al. (2019). Single-cell transcriptomics of human T cells reveals tissue and activation signatures in health and disease. Nat. Commun. 10.

Takeuchi, A., and Saito, T. (2017). CD4 CTL, a cytotoxic subset of CD4+ T cells, their differentiation and function. Front. Immunol. 8.

Tanay, A., and Regev, A. (2017). Scaling single-cell genomics from phenomenology to mechanism. Nature 541, 331–338.

Tusi, B.K., Wolock, S.L., Weinreb, C., Hwang, Y., Hidalgo, D., Zilionis, R., Waisman, A., Huh, J.R., Klein, A.M., and Socolovsky, M. (2018). Population snapshots predict early haematopoietic and erythroid hierarchies. Nature 555, 54–60.

Velten, L., Haas, S.F., Raffel, S., Blaszkiewicz, S., Islam, S., Hennig, B.P., Hirche, C., Lutz, C., Buss, E.C., Nowak, D., et al. (2017). Human haematopoietic stem cell lineage commitment is a continuous process. Nat. Cell Biol. 19, 271–281.

Wilson, A., Laurenti, E., Oser, G., van der Wath, R.C., Blanco-Bose, W., Jaworski, M., Offner, S., Dunant, C.F., Eshkind, L., Bockamp, E., et al. (2008). Hematopoietic Stem Cells Reversibly Switch from Dormancy to Self-Renewal during Homeostasis and Repair. Cell 135, 1118–1129.

Zheng, S., Papalexi, E., Butler, A., Stephenson, W., and Satija, R. (2018). Molecular transitions in early progenitors during human cord blood hematopoiesis. Mol. Syst. Biol. 14.

