## Supplementary Notes for "Single-cell proteo-genomic reference maps of the hematopoietic system enable the purification and massive profiling of precisely defined cell states"

#### Supplementary Note 1. Panel design for targeted transcriptomics

In order to establish a comprehensive targeted transcriptomics approach in the human bone marrow, we designed a panel that covers all cell types and differentiation stages of this organ. For this purpose, we used one of the individuals from the human bone marrow dataset released by the Human Cell Atlas project (Data obtained from Data Portal preview site: https://data.humancellatlas.org/explore/projects/cc95ff89-2e68-4a08-a234-480eca21ce79 ). We filtered out any cell for which fewer than 500 genes and more than 10% mitochondrial counts were detected. We performed unsupervised clustering and UMAP-based dimensionality reduction on the 33.000 cells that passed the quality filter. Subsequently, we annotated the cell types based on canonical markers from the human bone marrow. We obtained 17 cell types covering all the main lineages in the hematopoietic system (Figure N1a).

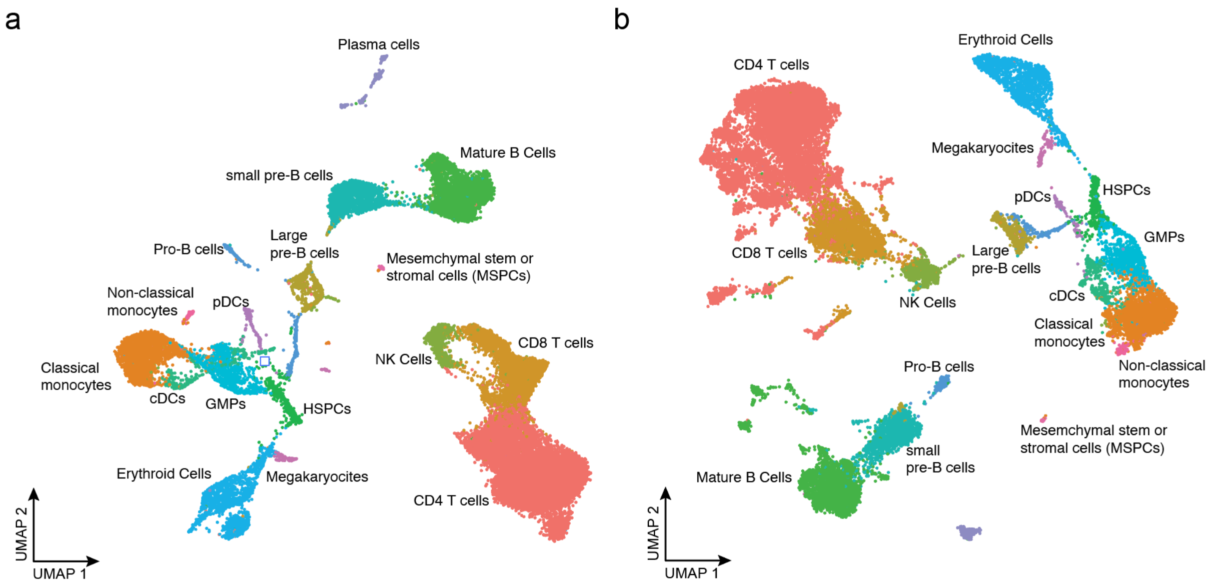

**Figure N1. Panel design for targeted single-cell transcriptomics of the human bone marrow.** Bone marrow scRNA-seq data from a healthy individual was obtained from the Human cell atlas, the data was processed and annotated. **a.** UMAP visualization of the whole transcriptome data highlighting the result of unsupervised clustering. **b.** UMAP based on the 462 gene panel selected for Abseq highlighting the clusters obtained in *a*.

In order to identify a sparse, yet maximally informative set of marker genes, we followed the approach for target selection in targeted single-cell transcriptomics developed previously (Schraivogel et al., 2020). In short, we determined the genes that were differentially expressed between cell types and used these as input to train a generalized linear model of cell type identity while applying a LASSO regularization in order to select a maximally sparse set of features. Regularization parameters were determined using 10-fold cross-validation. A total 257 genes were selected with this method. In addition to these genes, we included 83 cell cycle markers (Kowalczyk et al., 2015), 88 genes corresponding to the Abseq antibodies, and 75 genes with high variability in single-cell datasets of AML patients (Velten et al., 2018). Due to a poor representation of hematopoietic stem and progenitor cells in the cell atlas dataset, we complemented our targeted panel with 45 stem and progenitor cell markers previously identified using a similar approach (Velten et al., 2017). All selected genes and primer sequences are included in Supplementary Table S1. For primer design, a proprietary routine was followed (BD Bioscience).

The selected panel allowed us to classify the cell types of the human cell atlas dataset with a 99,92% of agreement and Cohen's kappa coefficient (κ) of 0.932. Moreover, we performed unsupervised clustering and dimensionality reduction using only the selected genes for the panel (Figure N1b) and obtained a partitioning highly similar to the clustering obtained when using whole-transcriptome data, as quantified by an adjusted Rand index of 0.98. The performance of the panel was further validated by comparing whole-transcriptome and targeted Abseq data (see Supplementary Note 2).

### Supplementary Note 2: Whole transcriptome sequencing validates performance of the targeted panel

In order to exclude that the targeted assay leads to biases in clustering or cell type annotation, we performed whole transcriptome sequencing (WTA) together with profiling of the same 97 antibodies on a sample from a healthy individual (Young3). For this, we processed 14,378 cells using the whole transcriptome protocol for the BD Rhapsody system. On average we sequenced ~60,000 reads (i.e. 7x deeper than with the targeted approach) for the RNA layer and 18,000 reads for the antibody layer per cell. After normalizing the counts by the library size, the top 3,000 highly variable genes and all 97 antibodies were used as input for the MOFA dimensionality reduction, unsupervised clustering, and UMAP calculations as described in the methods section. Thereby, 34 distinct clusters were identified (Figure N2a) Subsequently, we utilized the label transfer approach from Seurat v3 (Stuart et al., 2019) to predict the cell type identities using the targeted datasets as reference. We calculated the mutual overlap of the predicted labels with the unsupervised cluster (Figure N2b), and also projected the whole transcriptome data into the original reference space for comparison (Figure N2c, d and see also Supplementary Note 7). In the majority of cases, a 1:1 correspondence between clusters from the targeted approach and clusters from the whole transcriptome approach was observed. Some cell types were not covered in the WTA approach due to low cellular coverage. Together, these data suggest that our targeted panel resolves cell types equally well as the WTA approach at strongly reduced costs.

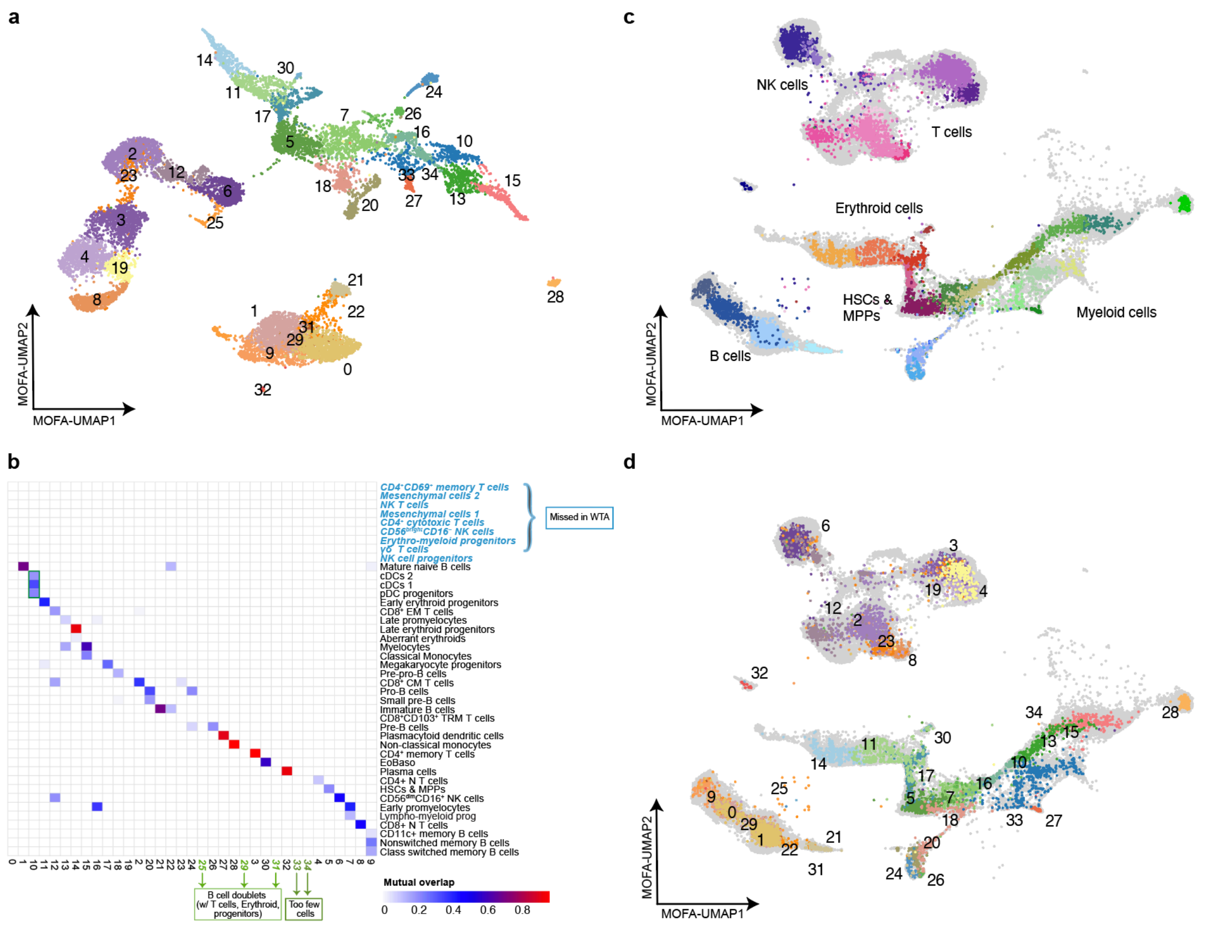

**Figure N2. Whole-transcriptome single-cell proteo-genomic sequencing. a.** MOFA-UMAP visualization of the whole transcriptome dataset colored by the unsupervised clusters. **b.** Mutuality analysis between the unsupervised clusters from the WTA and the identities predicted from the targeted datasets. Mutual overlap is defined as the product between the % of cells from a targeted cluster that form part of a given WTA cluster, and the % of cells from a WTA cluster that form part of a given targeted cluster. A mutual overlap of 1 indicates perfect overlap. **c.** Projection of the WTA samples in the healthy reference MOFA-UMAP space colored by predicted cell types **d.** and unsupervised clusters from *a*.

### Supplementary Note 3: Effects of ultra-high plex antibody stainings and freeze-thaw cycles on gene expression

In order to exclude that staining live, primary cells with 97 antibodies affects gene expression, we performed a control experiment. For this purpose, we used a sample from a healthy donor (Young1) and proceeded as described in the Methods section 'Cell sorting for Abseq'. Half of the sample was then incubated with 97 Abseq antibodies, while the other half was left on ice. Finally, the BD Rhapsody protocol was performed as described (see Methods 'Abseq surface labeling, single-cell capture and library preparation'). In order to compare both samples, only the transcriptomic information was considered. We normalized the datasets, performed unsupervised clustering, and annotated the clusters into major cell types from the bone marrow based on canonical markers (Figure N3a, b). All cell types were present in both conditions without considerable batch effects driving any type of separation. The correlation between the average expression of each gene between both samples was 0.996 (Figure N3c). To account for possible slight variations in cell type abundance, we further investigated the correlation between experiments at the level of individual cell types, and consistently found very high correlations (see Figure N3d for representative examples). Only the erythroid-specific gene HBB, and to a lesser extent HBD, were more abundantly observed in most cell types of the non-stained experiment, suggesting that there is an elevated HBB background signal in this experiment. HBB is frequently observed as "background" in different single-cell RNA-seq studies (Young & Behjati, 2020). HBB represents 96% of the RNA present in red blood cells and 80% of RNA in blood extracellular vesicles (Kerkelä et al., 2019). Hence, cellular debris present in prepared samples mostly contains HBB, explaining the abundance of this specific gene in the background signal. Here, the additional washes included during the antibody staining apparently have helped to reduce the background signal.

The datasets established in this study was obtained from cryopreserved bone marrow cells. To evaluate the effect of freeze-thaw cycles on gene- and surface antigen expression, we performed three control experiments with peripheral blood mononuclear cells (PBMCs). Blood was drawn, subjected to Ficoll density gradient centrifugation and PBMCs were frozen or left on ice for 6 hours; after this time interval, blood was drawn again from the same donor and subjected to Ficoll density gradient centrifugation. All three samples (PB fresh, PB on ice, PB freeze-thaw) were then processed together and living cells were FACS-sorted and stained with 97 surface antibodies before single cell capture. Globally, correlations in gene and surface antigen expression were very high (freeze-thaw vs. fresh = 0.994, freeze-thaw vs. on ice = 0.994 and fresh vs. on ice = 0.997, Figure N3j). UMAP visualization of the data revealed that cell types were unaffected by freezing except for the monocytes, showing a minor shift upon freeze-thawing (Figure N3g, h). As a result of the freeze-thaw process, monocytes upregulated the T cell costimulatory molecules CD275 (ICOS ligand) and the immediate early gene JUN, while downregulated the homing receptor CD62L (SELL), indicative of a stress response (Figure N3i). Global gene expression patterns remained unaffected. All in all, the freeze-thaw process had only a minor impact on our data. Specific changes associated with freeze-thawing (such as for monocytes) apply equally for samples that have been processed using the same pipeline.

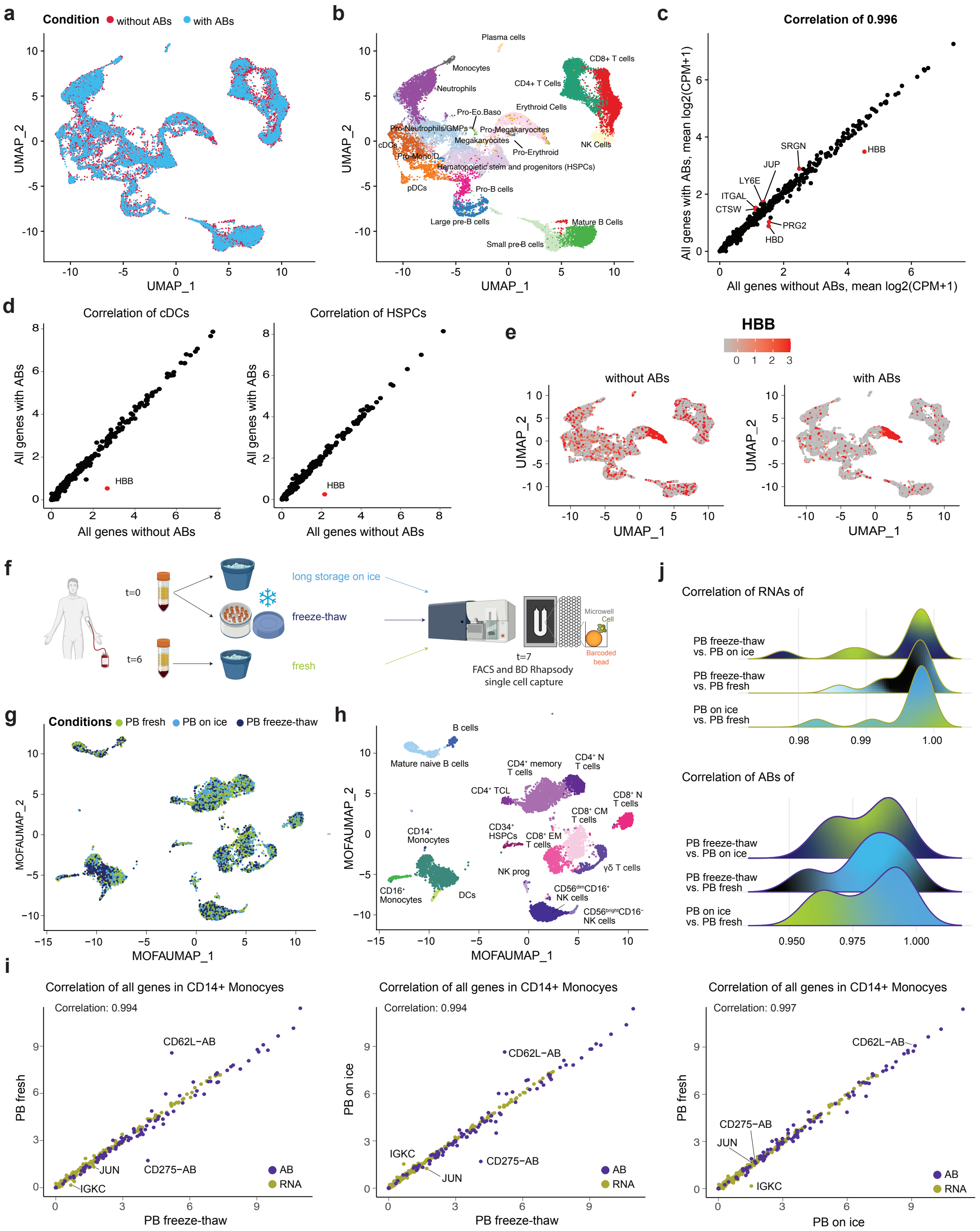

**Figure N3. Effects of ultra-high plex antibody stainings and freeze-thaw cycles on gene expression**. **a-e.** Comparison of a healthy bone marrow sample incubated with and without Abseq antibodies. **a, b.** A UMAP was generated on the targeted RNA single-cell profiles. The cells from either condition *(a)* and the annotated cell types *(b)* are highlighted. **c.** Correlation between the average expression of every gene in each sample; top differentially expressed genes are highlighted in red. **d.** Correlation between the average expression of every gene in two representative cell types (cDCs and HSPCs); HBB as most differentially expressed gene is highlighted in red **e.** UMAP visualization colored by the HBB expression in positive and negative samples. **f-j.** Comparison of fresh, frozen/thawed and iced PBMC samples. **f.**

Overview of the experimental setup **g, h.** UMAPs were generated on the fresh, iced and freeze-thawed PBMC samples sequenced with the 462 RNAs and 97 ABs targets. The cells are colored by their condition *(g)* and cell type *(h)*, respectively. **i.** Correlation between the average expression of each gene per condition. Average expression of JUN, IGKC, CD276-AB and CD62L-AB are highlighted. **j.** Ridge plots display the correlation coefficient of correlation between the average expression of every gene per cell type and condition. Cell types which were represented by less than 50 cells were excluded from the analysis.

### Supplementary Note 4: Analysis of sequencing requirements

In order to establish the minimal sequencing requirements necessary to obtain an accurate cell type classification, we down-sampled the RNA and AB counts of a healthy bone marrow sample (Young1) and determined if this would yield a similar cluster structure using the same unsupervised clustering pipeline as used for the non-down-sampled datasets. For illustration, data of all nine samples with an average UMI count of 10,126 per cell (Figure 1b) were down-sampled to an average of 300 UMIs per cell (60 UMIs antibodies, 240 UMIs RNA) (Figure N4a) and subsequently a UMAP was calculated. This revealed that even much lower read numbers are sufficient to distinguish cell types and differentiation states. While the total read/UMI counts may appear low compared to whole transcriptome single-cell studies, it is in fact very deep for a targeted approach, as analyzed in detail in (Schraivogel et al., 2020). To provide a more quantitative analysis, we performed down-sampling to different depths and determined the overlap in cluster structure using the adjusted Rand index (Figure N4c). Additionally, we used label transfers from the non-down-sampled dataset on the down-sampled dataset to determine if cell types could be correctly classified. The agreement between the true labels and the learnt labels was determined using Cohen's Kappa (Figure N4c). Both metric showed that the RNA counts were more important for the correct cell type identification and that we reached saturation with around 70% of the RNA reads, i.e. an average read depth of 4,000 per cell. All samples were sequenced to an average of ~8,500 reads for the mRNA libraries and ~12,000 reads per cell for the antibody libraries. A more specific analysis on the benefit of profiling both surface markers and RNA for cell type annotation, e.g. in the context of T cell populations, is presented in Supplementary Note 6.

We observed that some antibodies (e.g. CD18) occupy high percentages of reads whereas others are lowly represented (e.g. CD20, see Figure S1). The large variation in reads per antibody is not correlated with the fraction of cells positive for the antibody (e.g. both CD18 and CD20 are expressed in 19% of cells, but CD18 has 63 times more reads), indicating that it is primarily related to the abundance of the antigen on the surface, properties of the antibody, or properties of the panel. In future, unlabeled antibodies of the same clone (also referred to as cold competitors) could be included for highly represented antibodies to decrease the fraction of reads consumed by these antibodies and thereby reduce the overall sequencing requirements. The data shown in Figure S1 (also included in Table S5) can be useful for selecting these antibodies in future studies.

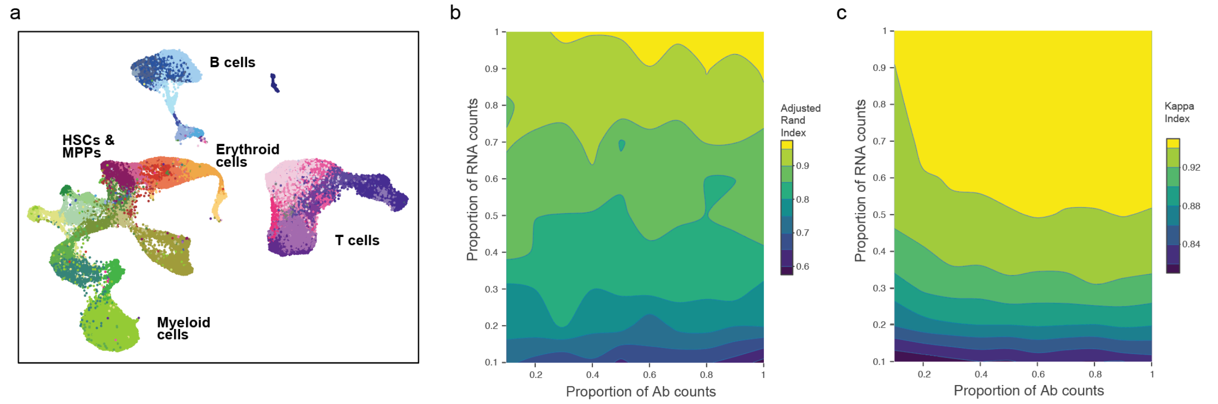

**Figure N4. Sequencing requirements. a.** UMAP of the nine bone marrow samples down-sampled to an average of 300 UMIs (60 UMIs antibodies and 240 UMIs RNA). Cells are colored corresponding to *Figure 1b*. **b, c**. Effect of down-sampling RNA or antibody reads on cell type classification by *(b)* unsupervised clustering or *(c)* label transfer.

### Supplementary Note 5: Cell type annotation

For cell type annotation, we used both information of the mRNA and cell surface antibody readouts. Especially for inter-connected clusters within the hematopoietic differentiation hierarchy and their branches along myeloid and lymphoid trajectories, fine gradients of surface marker expression and mRNA expression exist, which result in slight expression changes from one annotated cluster to the next. Supplementary Table S4 summarizes the surface markers and mRNAs used for annotation of the 45 clusters. For additional comparisons and analysis of differential mRNA and surface marker expression between any individual cluster of choice, usage of the Abseq-App and Supplementary Table S5 is highly recommended.

Hematopoietic stem cells and multipotent progenitors (HSCs & MPP) are defined via surface expression of CD34, CD133 and absence/lower levels of CD38, CD45RA and Tim3 expression. HSCs can be further characterized by robust CD90 and substantial CD4 expression. On the mRNA level, HSCs and MPPs were defined by high expression of CRHBP, NPR3, MEIS1, PROM1 and THY1 (Figure N5a). Erythro-myeloid progenitors (EMP) are emanating directly from the HSC and MPP cluster, maintain CD34 expression but lack CD133, CD38, CD4, CD11a or CD45RA surface expression. They are characterized via high expression of CPA3, FCER1A, DEPTOR and TESPA1 on the mRNA level (Figure N5a). Downstream of EMPs, early erythroid progenitors and late erythroid progenitors can be characterized by presence of CD34 and CD38 surface expression. Moreover, CD98 and CD326 start to be expressed at marked levels in early erythroid progenitors and further increase in late erythroid progenitors (Figure N5b). At the stage of the late erythroid progenitor cluster, surface expression of CD235 can be observed (Figure N5b). In addition, SOX4, APOC1, IL1B, CASP3 and SLC40A1 are highly expressed in early erythroid progenitors, whereas late erythroid progenitors highly express hemoglobins (HBB, HDD), CA1, BLVRB, AHSP and in later stages GYPA, encoding Glycophorin A. Of note, TFRC mRNA (encodes for transferrin receptor or CD71) starts to be highly expressed in EMPs and is constitutively expressed throughout erythroid differentiation. An additional erythroid cluster with enrichment in old and leukemic individuals could be observed next to late erythroid progenitors, termed aberrant erythroid cells, which was positive for CD235a, CD123, CD197 and CD90 surface and high levels of HBB, HBD, CA1, HEMGN, RHAG mRNA expression (Figure N5b). Megakaryocyte progenitors were defined by high surface expression of CD61 and CD49b and mRNA expression of PPBP, ITGA2B, PLEK, MPL, FERMT3 and SLC37A1 (Figure N5b). As shown previously, Eosinophil/Basophil progenitors showed similarities in gene expression with other cell types along the erythroid differentiation trajectory, including high expression of CPA3 and FCER1A. Other genes specifically expressed in this cell type are CLC, HDC, PRG2. Interestingly, RNASE2 and CPA3 are also markedly expressed in Eosinophil/Basophil progenitors, but also expressed at high levels in cell clusters along the myeloid differentiation trajectory (Figure N5b). Lympho-myeloid progenitors emanate from the HSC & MPP cluster and were defined by their expression of CD34 and CD133, absence/low levels of CD38 and dim expression of CD45RA, CD49b, Tim3, CD117 and in some cells also CD10. On the transcriptomic level, cells within the Lympho-myeloid progenitor cluster expressed high levels of FLT3, SPINK2, MZB1, ITM2C and MDK (Figure N5a).

Directly adjacent to the Lympho-myeloid progenitor cluster, early promyelocytes, that represent an intermediate myeloid differentiation stage can be characterized by their surface expression of CD34, CD38, CD133, CD117 and CD33 (Figure N5a and N5i). Signature genes highly expressed in the early promyelocytes cluster comprise MPO, AZU1, CPA3, RNASE3, MGST and CEBPA. Following the early promyelocytes stage, the subsequent myeloid differentiation stages are referred to as late promyelocyte and myelocyte stages. In our dataset, cells within the late promyelocyte cluster gradually lose CD34, CD133 and CD117 expression, whereas CD38 is increased and CD33 expression remains at levels comparable to the early promyelocytes stage. Late promyelocytes can be further characterized by their high transcriptomic expression of LYZ, CSTA and RNASE2, and slightly lower MPO and AZU1 expression if compared to the early promyelocytes stage (Figure N5i). Myelocytes represent the next transient myeloid differentiation stage, which lost their CD34 and CD133 surface expression, and can be clearly distinguished from late promyelocytes by the emergence of CD11b expression on their surface. Moreover, they highly express CD33 and CD11a and upregulate CD49e expression (Figure N5i). They can be further characterized by high mRNA expression of S100A8, CSTA and S100A9. Classical monocytes show overlap in their surface marker expression to myelocytes, however they express additional markers like CD14, CD61 and CD32, and express higher levels of CD13 than their progenitors (Figure N5i). On the transcriptomic level, classical monocytes highly express CD14, VCAN, CTSS and FCN1. Non-classical monocytes appear to be the terminal cluster within the myeloid differentiation trajectory and can be distinguished from non-classical monocytes by high expression of surface CD16 and CD127, as well as FCGR3A, LST1 and MS4A1 on the transcriptomic level (Figure N5i).

Next to the manifold myeloid differentiation stages, plasmacytoid dendritic cell progenitors (pDC prog) and mature plasmacytoid dendritic cells (pDC), as well as both conventional dendritic cells of type 1 (cDC1) and type 2 (cDC2) are mapped (Figure N5j). On their surface, pDC progenitors express CD34 and CD133 and in addition high levels of CD38 and CD123. Moreover, they can be further characterized by dim expression of CD33, CD45RA and Tim3, as well as high CD98 expression. Signature genes highly expressed in pDC progenitors encompass IRF8, IL3RA, SHD and DNTT. Mature pDCs gradually lose surface CD34 expression, remain positive for CD38 and also express higher levels of CD45RA than their progenitors. Besides high CD123 expression, these cells can also be further characterized by high surface expression of CD61, CD98 and CD4 as well as dim CD10 expression. On the transcriptomic level, IRF8, IL3RA, CCDC50, CD4, TCF4 and DERL3 are highly expressed in pDCs. Conventional dendritic cells of type 1 show dim levels of CD34 and CD133 surface expression, and can be defined by high expression levels of CD33, CD11c, CD141 and Tim3. S100A10, FCER1A, HAVCR2 and CD2 showed high mRNA expression levels in cDC1 cells. The second type of conventional dendritic cells, cDC2, can be delineated by their absence of surface CD34 and CD133 expression, lower CD38, CD45RA and CD141 levels than their cDC1 counterparts and high expression of CD11c, Tim3 and additionally CD1c. Transcriptionally cDC2 are similar to cDC1, however can be further separated by higher expression of FCER1A, CD1C and HLA-DQB1.

Emanating from the Lympho-myeloid progenitor cluster, B cell commitment takes place, which is accompanied by gradual elevation of CD10 surface expression. Cells within the first B cell developmental stage, i.e. pre-pro B cells, express CD34 and dim levels of CD133 and CD38. CD10 expression is present, but to a significantly lower degree compared to later B cell differentiation stages. Moreover, surface markers that are gradually upregulated in later stages, like CD19, CD81, CD24 or CD9 are absent in this early differentiation stage. Pre-pro B cells and overall B cell commitment can be further characterized by high expression of MME, VPREB1, CD79A and DNTT (Figure N5c). Moreover, the transcription factor PAX5 is highly expressed at this differentiation stage (see WTA expression data in Abseq-App). After the pre-pro B cell stage, cells enter and pass sequential stages, i.e. the pro-B cell, and pre-B cell stages. Of note, our clustering splits pro-B cells and pre-B cells according to distinct cell cycle states, which cells undergo being in early and late pro-B cell or pre-B cell stages. Therefore, we labelled these clusters cycling pre-B and pro-B cells and non-cycling pre-B and pro B cells (Figure N5c). In more detail, cells forming the former cluster are in a highly proliferative state (high S-phase score, low G2/M score), whereas the latter are in a halted pre-mitotic, quiescent state, in which DNA-rearrangements known as heavy and light chain rearrangements take place. In pro-B cells, the latter state corresponds to cells that already underwent successful DJ-recombination at the Igm locus, after which the above-mentioned proliferative state is entered, finally leading to VDJ rearrangement that is accompanied by Rag2 expression (see WTA dataset in the Abseq-app). Successful heavy chain rearrangement also marks the transition into the pre-B cell stage and to the expression of the so-called pre-BCR, which corresponds to surface IgM coupled to surrogate light chains. IGHM mRNA expression is increased upon entry into pre-B cell stage, accompanied by high proliferation, which terminates as the cells enter the small pre-B cell stage in which Igk light chain rearrangement takes place. Successful completion of all BCR rearrangements and surface expression of a functional BCR renders B cell entry into the immature B cell stage.

Immature B cells are thought to exit the BM and start to transit through secondary lymphoid organs, where final maturation events like isotype switching and somatic hypermutation occur. However, these processes also occur in the bone marrow in germinal center-like structures (Cariappa, Chase, Liu, Russell, & Pillai, 2007). B cells in that stage can be defined via CD9 surface expression, which is highest in immature B cells, as well as expression of CD81, and in late stages CD272, CD20, and IgD, which marks the start of isotype switching (Figure N5d). Importantly, immature B cells lack CD185 expression, which plays an important role for germinal center organization and is a signature surface marker for subsequent maturation stages. CD9, CD79B, CD24 and TCL1A are highly expressed on the transcriptional level at this stage. As expected, in late stages of the immature B cell cluster, isotype switching from IgM to IgD can be observed, marking the transition to mature naïve B cells. Mature naïve B cells express higher levels of surface CD5 than other B cell developmental stages, and temporarily lower their CD24 surface expression. Cells at this stage also highly express IL4R and IGHD mRNAs. Non-IgG-class switched B cells succeed mature naïve B cells and this stage can be characterized by elevated surface expression of CD24, CD25, CD27, CD1c and IgD in the absence of IgG expression (Figure N5d). High expression levels of CD82, CD27, JCHAIN and CD1C mRNAs are transcriptomically characteristic for this stage. After B cells switched their isotype from IgD to IgG, high surface IgG expression and absence of IgD expression can be observed (Figure N5d). Moreover, transcriptionally class switched cells express elevated CD82, ITGB1 and ITGAM mRNAs. Adjacent to class-switched B cells, a small cluster of CD11c+ memory B cells is apparent, whose exact biological function remains elusive (Figure N5d). They have been described as putative progenitors of antibody secreting plasma cells and have also been studied in the context of autoimmune diseases (Golinski et al., 2020; Karnell et al., 2017). This cell subset is characterized by their high surface levels of CD11c, IgG, dim levels of CD279 and lack of CD185 (CXCR5). Lack of CD185 expression indicates that cells of this cluster are not present in follicular structures any longer. The final stage of B cell development is represented by plasma cells, which express high surface levels of CD38, CD27 and CD54. On the transcriptomic level, high expression of MZB1, DERL3, FKBP11 and SEC11C is observed in this cluster.

Besides B cells, CD3+ positive T cells, which passed several differentiation stages in the thymus, re-locate to the bone marrow. One can generally distinguish between alpha-beta T cells and gamma-delta T cells, which describes an intrinsic difference in the respective T cell receptor (TCR) composition. Alpha-beta T cells can be further separated in CD4+ and CD8+ T cells, that are either CD45+, CD3+, TCRab+, CD4+ or CD45+, CD3+, TCRab+, CD8+ respectively. Of general note, inclusion of the surface marker information greatly aids T cell annotation, as their RNA content is low and many gene signatures are shared either between individual CD4+ and CD8+ T cell clusters or shared with other cytotoxic cells, like natural killer cells.

Within the former, naïve CD4+ T cells can be further characterized by high expression of CD45RA, CD28 and CD197 (CCR7) (Figure N5e). They are devoid of surface CD95, CD279 and CD25 and further express high levels of CD4 mRNA, as well as CCR7, SELL and AIF1. Upon antigen encounter, naïve T cells give rise to memory CD4+ T cells, which is accompanied by changes in their surface proteome and transcriptome. Memory CD4+ T cells express CD28, don't express surface CD45RA or CD197, and upregulate CD25 expression (Figure N5e). Transcriptionally, KLRB1, CCR6 and CD82 expression levels are elevated in this cell cluster. Besides memory CD4+ T cells, two more CD4 positive clusters adopted a memory phenotype, but differed quite significantly to the former. We found one cluster, which we termed cytotoxic CD4+ T cells, to be highly positive for cytotoxicity-related mRNAs like GNLY, GZMA, NK7, and ADGRG1 and devoid of any CD28 expression (Figure N5e). The second cluster, CD69+ CD4+ T cells, highly expressed CD69 and CD279 on its surface, and CD81, CST7, TGFBI and HLA-DPBI mRNAs were elevated (Figure N5e). These cells likely constitute CD4 tissue resident T cells, since both CD69 and CD279 are considered tissue resident memory T cell markers (Kumar et al., 2017).

Naïve CD8+ T cells are transcriptionally highly similar to naïve CD4+ T cells, express high levels of surface CD197 and, in contrast to their CD4+ T cell counterparts, express CD314. Furthermore, naïve CD8+ T cells are lacking CD94, CD279 and CD103 surface protein expression (Figure N5f). Transcriptionally, cells within this cluster highly express CCR7, SELL, and AIF1 mRNAs. Upon antigen encounter, naïve CD8+ T cells adopt a central memory T cell phenotype, and are henceforth termed CD8+ central memory T cells, which downregulate CD45RA surface protein and lack CD197, CD103 and CD94 expression. Moreover, surface CD279 and CD81 is elevated in this cluster compared to other bone marrow CD8+ T cell subsets (Figure N5f). Targeted mRNA readouts indicated high expression of CD8A, DUSP1, CD74 and CCR5. A second memory phenotype cluster, termed CD8+ effector memory T cells, could be observed within CD8+ T cells, which showed substantial expression of surface CD45RA, CD226, and CD94 in the absence of CD103, CD279, CD27, CD28 expression (Figure N5f). Cells within this cluster express high levels of genes involved in cytotoxicity related processes. Another prominent cluster showing a memory T cell phenotype were CD8+ CD103+ tissue resident memory T cells. They can be characterized by high surface expression of CD103, CD25 and CD26 in the absence of CD45RA and CD197 expression (Figure N5f). On the transcriptional level, these cells robustly express IL7R, ITM2C and DPP4 mRNAs.

Besides CD4+ and CD8+ $\alpha\beta$-T cells, $\gamma\delta$-T cells, are apparent in our dataset. Within that $\gamma\delta$-T cells cluster, one subset was highly positive for surface CD226, CD26, CD94 and negative for CD45RA expression (Figure N5g). In the second subset, CD26, CD226 and CD94 expression is absent or only dim, but TCR-$\gamma\delta$ (clone B1) is highly expressed (Figure N5g). Interestingly, TCR-$\gamma\delta$ antibody clone B1, which is present in the 97 Ab panel seems to be less efficient in $\gamma\delta$-T cell detection than clone 11F2, which was present in the 197 Ab panel (Figure S4). The anti-Vd2 TCR antibody (clone B6) uniquely labels the CD226+ CD26+ CD94+ CD45RA- subset and leaves the other subset unstained. We therefore recommend using both anti-Vd2 (B6) and anti TCR-$\gamma\delta$ (11F2) for improved $\gamma\delta$-T cell detection. On the transcriptional level, KLRB1, TRDC, TRGC and DPP4 were highly expressed in this cluster.

The second CD3+ TCRαβ-negative cluster are putative natural killer T cells (NKT cells), that can be characterized by elevated CD69 and CD314 surface expression, which were accompanied by dim binding of the TCRγδ antibody (clone B1).

Next to CD3+ T cells, two well-known natural killer (NK) cell clusters, namely CD56$^{dim}$ CD16+ and CD56$^{bright}$ CD16- NK cells, are present in our dataset. Both subsets are readily identified via high surface expression of CD45, CD7, CD94 and CD45RA (Figure N5h). CD56$^{bright}$ CD16- NK cells express higher levels of CD56 and CD335, as well as lower levels of CD16 on their surface. In contrast, CD56$^{dim}$ CD16+ NK cells have higher surface levels of CD16, CD127 and CD152. The latter are thought to be more cytotoxic, which is also reflected in the transcriptional differences between the two subsets. Compared to other clusters in the dataset, generally cytotoxic mRNAs are highly expressed in these two clusters. In addition, a small cluster located in proximity of HSCs and MPPs was particularly interesting, as it expressed mature NK surface markers like CD16, CD56 and CD7 as well as surface markers specific for immature progenitor cells like CD34 and CD133 ((Figure N5a and N5h). A similar phenotype was observed at the mRNA level, as these cells both expressed mature NK mRNAs like NKG7 or KLRK1 and stem and progenitor specific mRNA like CRHBP, CD34 and NPR3. We therefore named this cluster NK cell progenitor. Besides healthy hematopoietic cells, at least one mesenchymal stromal cell (MSC) cluster is present in our dataset (Figure N5k). MSC cluster 1 was characterized by high surface expression of CD10, CD13, CD26 and CD49a, which was accompanied by typical MSC gene expression like CXCL12 and SPARC. Putative MSC cluster 2 expressed antibodies found in scavenger cells like macrophages, such as CD206, CD141, CD163 and CD16, but also expressed CXCL12 suggesting a mixed composition and some degree of heterogeneity.

All clusters described above were consistently identified in six healthy BM donors. In the reference AML patients (n=3, Figure 1b), we were able to determine three additional cell clusters. Some of these were either specific for individual AML samples or a mix of cells from different AML patients. Regarding the latter, the cluster annotated as immature blasts is a mixture of cells from all three patients. Cells within this cluster have heterogeneous surface phenotypes and share similarities with different healthy cell types (Figure N5l).

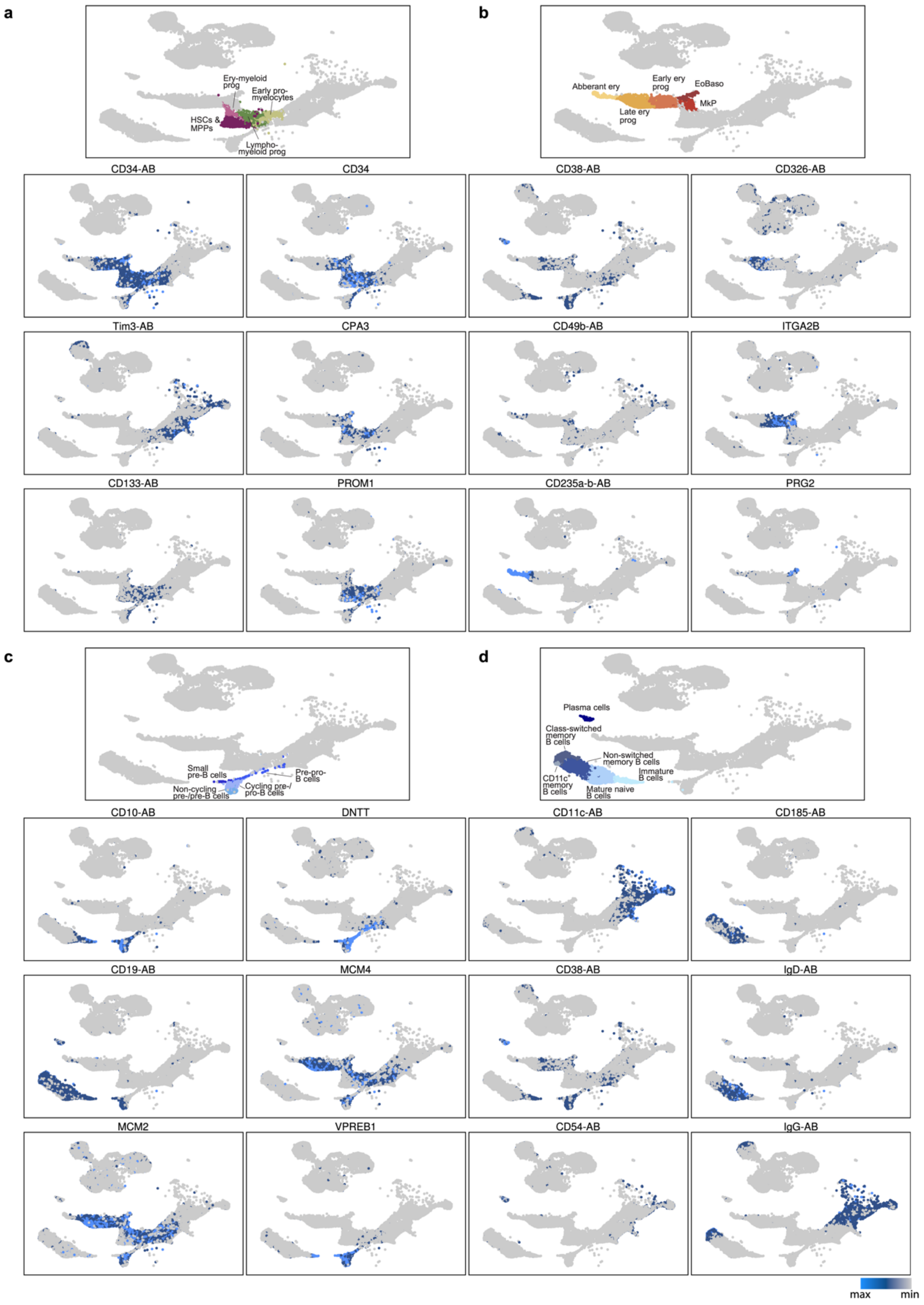
a
Ery-myeloid prog
Early pro-myelocytes
HSCs & MPPs
Lympho-myeloid prog
b
Abberant ery
Early ery prog
EoBaso
MkP
Late ery prog
CD34-AB
CD34
CD38-AB
CD326-AB
Tim3-AB
CPA3
CD49b-AB
ITGA2B
CD133-AB
PROM1
CD235a-b-AB
PRG2
c
Small pre-B cells
Pre-pro-B cells
Non-cycling pre-/pre-B cells
Cycling pre-/pro-B cells
d
Plasma cells
Class-switched memory B cells
Non-switched memory B cells
Immature B cells
CD11c+ memory B cells
Mature naive B cells
CD10-AB
DNTT
CD11c-AB
CD185-AB
CD19-AB
MCM4
CD38-AB
IgD-AB
MCM2
VPREB1
CD54-AB
IgG-AB
max
min

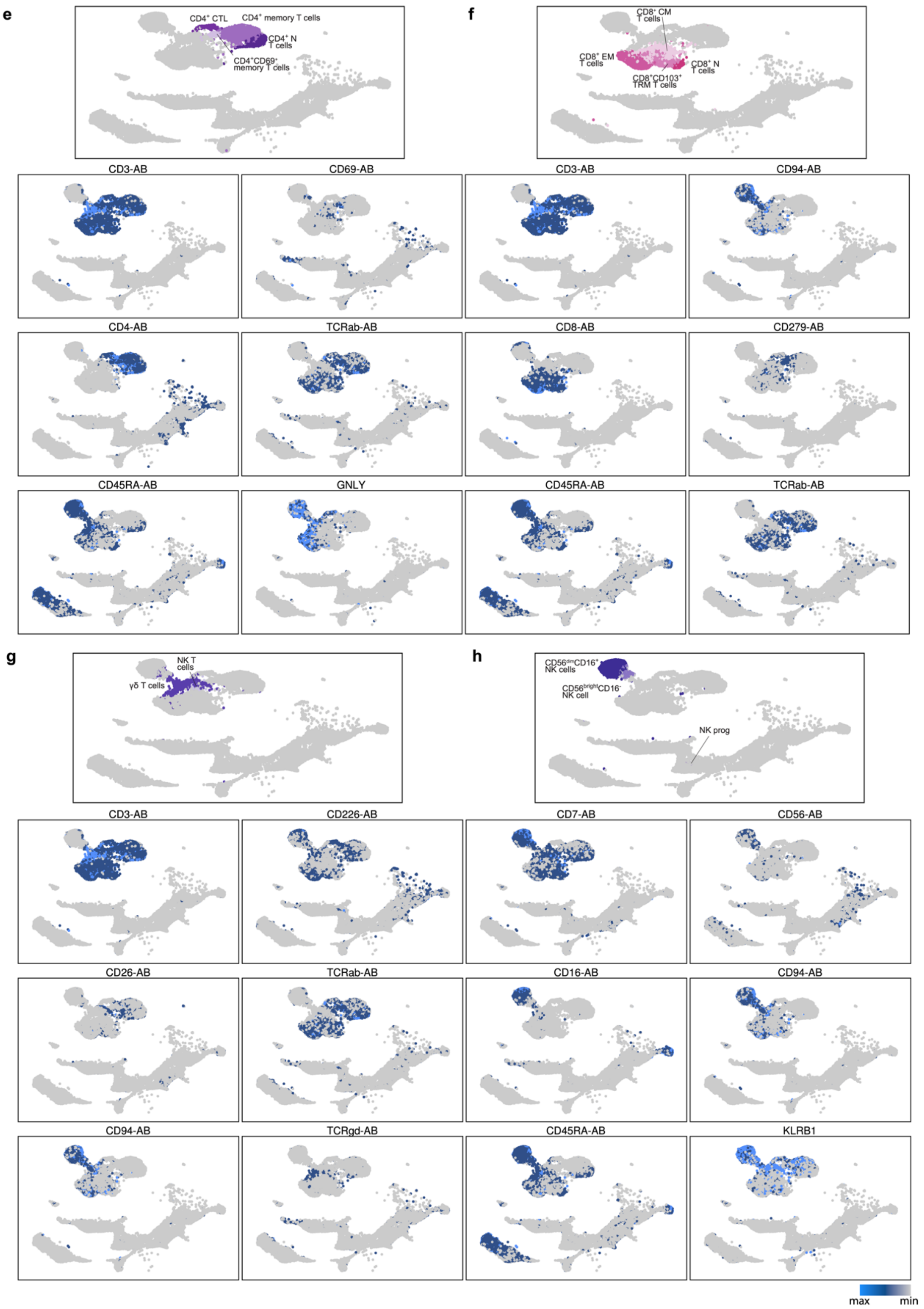

e
CD4+ CTL
CD4+ memory T cells
CD4+ N T cells
CD4+CD69+ memory T cells
f
CD8+ CM T cells
CD8+ EM T cells
CD8+ N T cells
CD8+CD103+ TRM T cells
CD3-AB
CD69-AB
CD3-AB
CD94-AB
CD4-AB
TCRab-AB
CD8-AB
CD279-AB
CD45RA-AB
GNLY
CD45RA-AB
TCRab-AB
g
NK T cells
γδ T cells
h
CD56dimCD16+ NK cells
CD56brightCD16- NK cell
NK prog
CD3-AB
CD226-AB
CD7-AB
CD56-AB
CD26-AB
TCRab-AB
CD16-AB
CD94-AB
CD94-AB
TCRgd-AB
CD45RA-AB
KLRB1
max
min

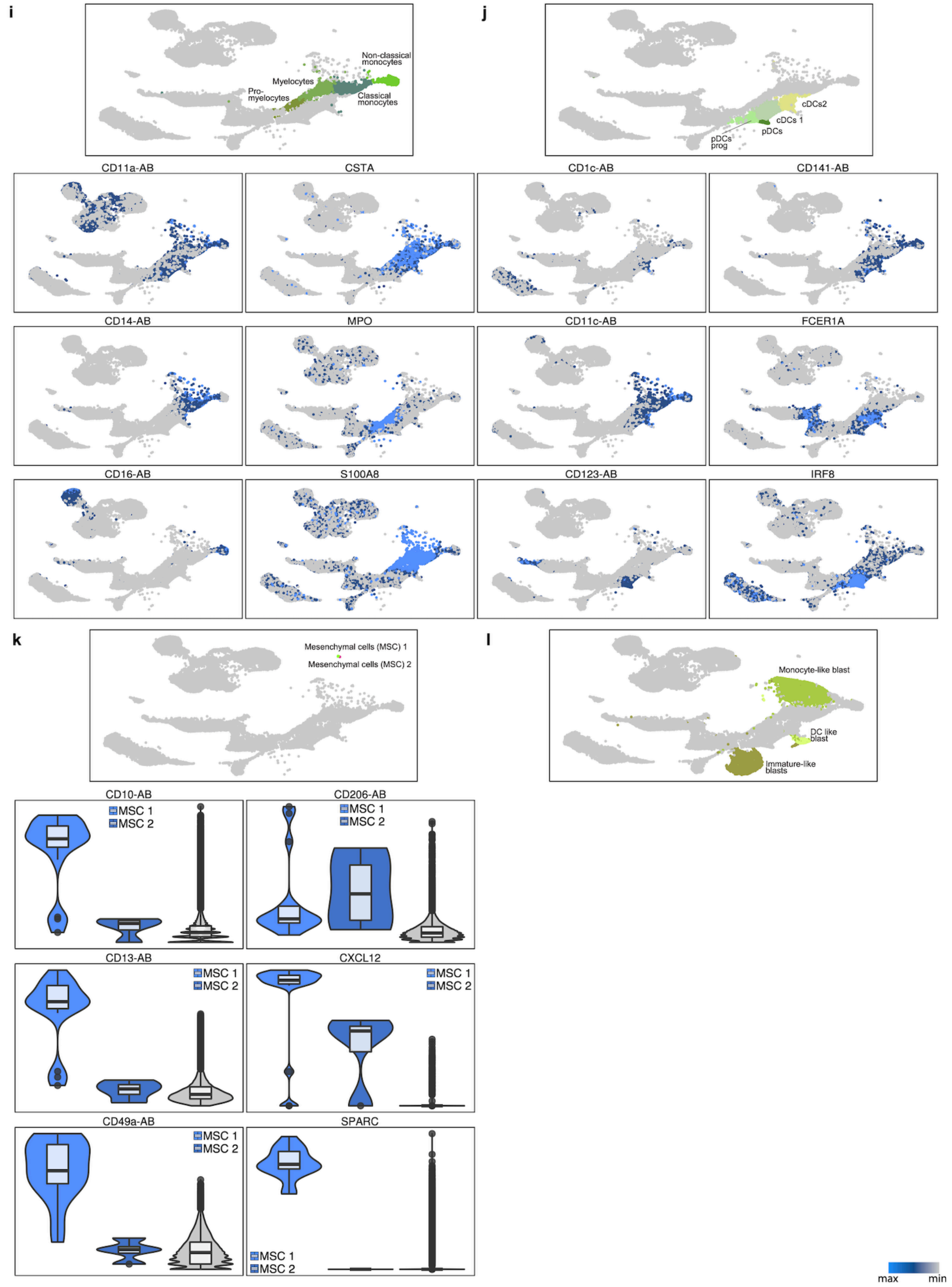

**Figure N5. Cell type annotation.** Both mRNA and surface protein expression were considered for annotation, and several exemplary plots for either mRNA or surface protein expression are shown. Cell types depicted in the UMAPs visualizations were grouped into biological subgroups.

**Supplementary Note 6: Inclusion of surface marker data improves cell type classification**

In order to determine the utility of RNA and surface marker expression for resolving cell types and cell stages, we performed UMAP visualizations using either RNA or surface marker information (Figure N6) or UMAP visualization after MOFA-UMAP multi-omic integration of RNA and surface markers (Figure 1b). Interestingly, dependent on the cell type, RNA or surface marker information were more powerful in separating cell types and cell stages (Figure N6). For example, for the HSPC compartment, RNA was superior over surface markers in resolving cell states, while B cell stages and the myeloid compartment were more efficiently separated using surface markers (Figure N6). Most importantly, the combined information of both layers provided the highest resolution not achieved by any of the individual layers alone (compare to Figure 1b). Interestingly, NK cell progenitors, a subset that we found to contain both features of mature NK cells (surface: CD56, CD7, mRNA: NKG7, KLRK1) and hematopoietic stem and progenitor cells (surface: CD34, CD133, mRNA: CD34, CRHBP, NPR3) was grouped with the HSPC compartment when only using RNA markers and with the NK cell compartment when only using the surface markers. Only in the joined view (Figure 1b), these cells formed an independent cluster, identifying a cell population that has hitherto remained hidden. Similarly, T cell subpopulations were resolved much better when using information from both omics layers. For example, populations such as γδ-T cells and NK T cells did not form an independent group in either of the individual layers and information from both layers was required to resolve them. In addition, usage of surface markers improves the separation between cytotoxic T cells and NK cells.

Our results imply that during stem cell differentiation, mRNA expression is a relatively early step in the process of commitment, compared to surface protein expression. In line with that, we and others have consistently observed lineage priming signatures in cells that surface phenotypically appear immature (see Figure 6d, Figure S9a, b and see also Paul et al., 2015; Velten et al., 2017). By contrast, in mature cell stages, cellular identity is firmly established and reflected both in the transcriptome, and on surface protein expression. In these mature cell types, antigen expression adds information to mRNA expression alone for three reasons: First, especially in T cells, mRNA measurements are often noisy due to the low RNA content of the cells. Second, in T cells and B cells, the annotation of cell types has historically been performed using surface antigens. Relatively similar cell states may have therefore been classified as functionally different based on the expression of a single marker, as in the case of class-switched vs. non-switched memory B cells, that mostly differ in the expression of surface immunoglobulins (IgM, IgD vs. IgG, IgA) while maintaining a very similar transcriptome. Third, a technical reason for our observations may be that the antibody panel (97 antibodies selected based on availability) can be biased towards providing higher resolution in specific cell types, whereas the mRNA panel was designed more systematically (see Supplementary Note 1).

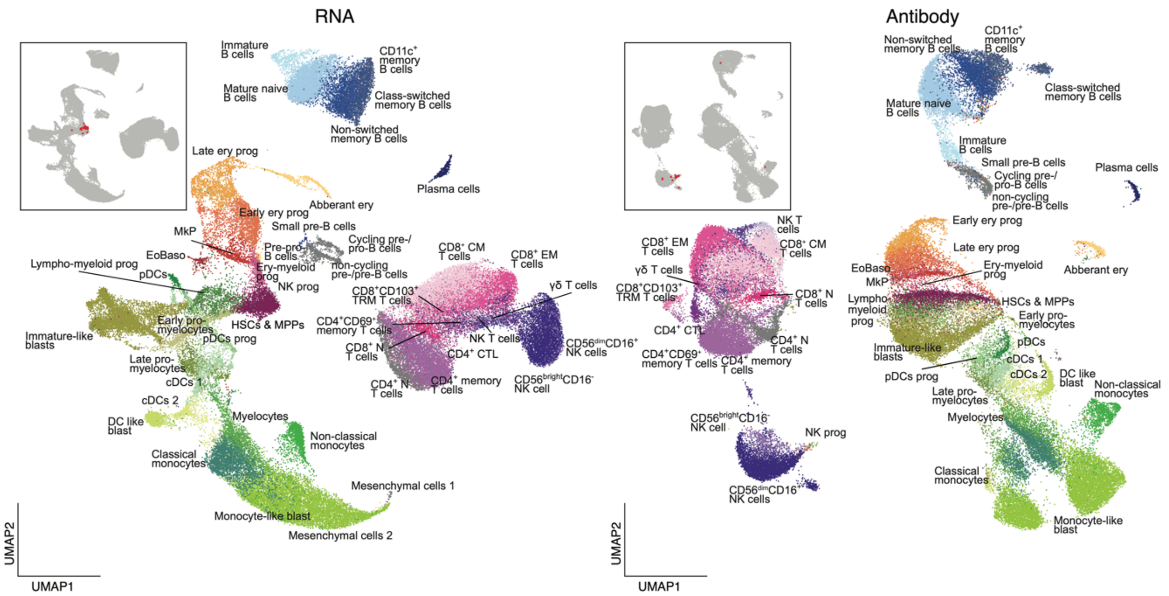

**Figure N6. Inclusion of surface marker data improves cell type classification.** UMAP visualization was calculated using either only RNA, or only surface marker expression. NK progenitor cells are highlighted in red. A UMAP visualization containing the combined information of RNA and surface markers is displayed in the main *Figure 1b*.

**Supplementary Note 7: Validation of query datasets projections**

To project new "query" single cell RNA-seq data on the reference atlas, the reference was subset to include only cells from the healthy individuals. For every query cell, we then determined the five nearest neighbors using scmap (Kiselev, Yiu, & Hemberg, 2018). The median UMAP position and cell type label of the five neighbors was then computed. For projecting on pseudotime, the average pseudotime value of the neighbors was computed if they were part of a given trajectory, i.e. not classified NA. The high accuracy of this mapping strategy was confirmed using both the 200-antibody experiment and the whole transcriptome experiment (Supplementary Note 2); in both cases, projected cell types had the same marker gene expression pattern as the cell types used in the original annotation.

For a more quantitative analysis, we evaluated the effectivity of this approach by projecting the healthy reference against itself. For a randomly selected set of cells, we showed that cells consistently projected very close to their original location (Figure N7a); the only exception were cells falling into the very heterogeneous and small class of mesenchymal stromal cell 2. We then calculated the precision of the projection as the proportion of correctly assigned cell type labels (Figure N7b). This showed that for most cell types the projection and mapping of the cell type had a precision higher than 0.8, with just a few populations having a lower precision, likely due to projection to a very similar cell state (Figure N7c).

The entire workflow of projection on the reference, differential expression testing and estimation of inter-patient variability is available at https://git.embl.de/triana/nrn

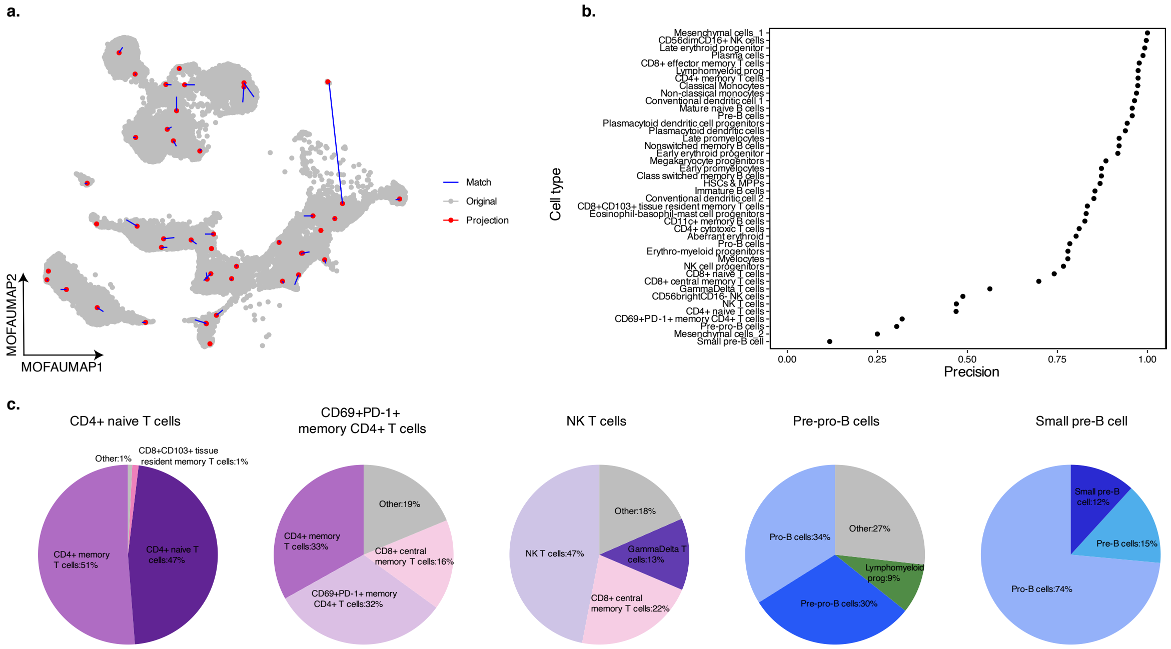

**Figure N7. Evaluation of self-projection on the healthy reference. a**. Precision of the cell type assignment base on the projection of the healthy reference against itself using scMAP. **b.** UMAP projection of one cell of each cell type (red) against the whole reference (gray) and the connection between the original coordinate and the projected one (blue). **c.** Distribution of the cell type projection in representative cell types with low assigment precision.

### Supplementary Note 8: Smart-seq2 validation

To validate markers identified in Figure 2 and 3, and gating panels described in Figure 6, we performed two experiments that couple FACS-based indexing of surface markers to single-cell RNAseq ("index scRNAseq"). For this purpose, we used Smart-seq2 and successfully sequenced 630 single cells that were stained with the data-defined classification panel established in Figure 6, and 330 single cells that were stained with a semi-automated panel aiming at resolving the erythro-myeloid differentiation of HSCs. The semi-automated panel included the backbone from the classification panel, but additionally contained the well-known erythroid priming marker CD71 (encoded by TFRC), the newly identified erythroid commitment marker CD326 (Figure 3), the eosinophil-basophil progenitor marker FcER1A, as well as CD49b, a marker highly expressed in megakaryocyte progenitors (see Supplementary Table S7). Additionally, we included 1035 cells from a previously published HSPCs dataset (Velten et al., 2017) with indexed single-cell RNAseq using the classical Doulatov gating scheme (Doulatov et al., 2010) into the analysis. To visualize our data, we performed dimensional reduction by Principal Components Analysis (PCA) and UMAP. We identified groups of similar cells using Shared Nearest Neighbor (SNN) clustering followed by data integration using canonical correlation analysis and mutual nearest neighbors as implemented in Seurat v3. A UMAP representation of the integrated dataset demonstrated a successful data integration (Figure N8a). Unsupervised clustering revealed the presence of multiple cell subpopulations adequately representing HSPC differentiation (Figure N8b). For all clusters we identified markers defined as those with the highest classification power as measured by the area under the receiver operating characteristics curve (AUC). The top five markers per cell type are shown in Figure N8d. Cell type labels (Figure N8c) were determined using the same marker genes as for the main Abseq data set (see Table S4 and Supplementary Note 5).

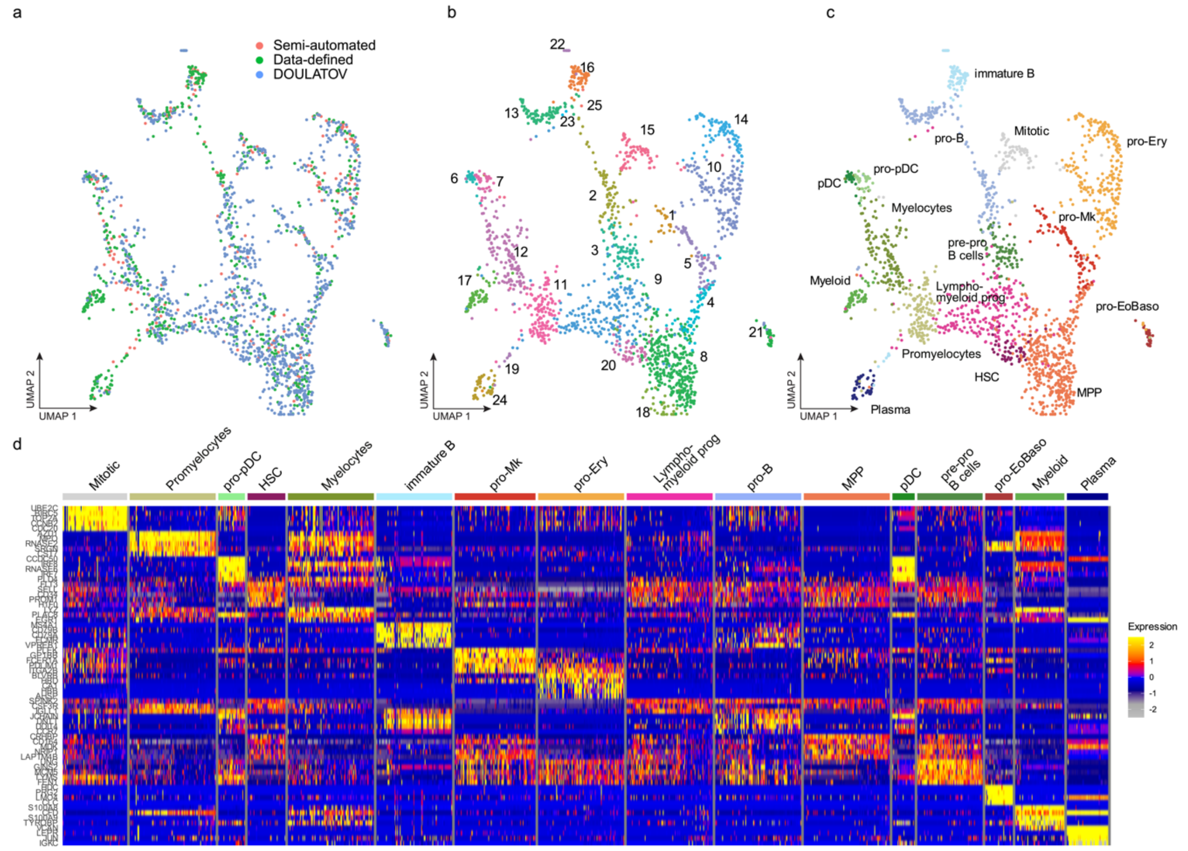

**Figure N8. Annotation of the Smart-seq2 dataset. a-c**. UMAP visualization of different “index scRNAseq” datasets colored by surface markers recorded (“indexed”) in the experiment *(a)* (Data-defined: Surface markers from data-defined panel from *Figure 6*; Semi-automated: Markers defined in the text above; DOULATOV: Markers from Velten et al., 2017), unsupervised clustering *(b)* and cell type annotation *(c)*. **d.** Heatmap of top marker genes for each cluster / cell type.

### References

Cariappa, A., Chase, C., Liu, H., Russell, P., & Pillai, S. (2007). Naive recirculating B cells mature simultaneously in the spleen and bone marrow. *Blood*, *109*(6), 2339–2345. https://doi.org/10.1182/blood-2006-05-021089

Doulatov, S., Notta, F., Eppert, K., Nguyen, L. T., Ohashi, P. S., & Dick, J. E. (2010). Revised map of the human progenitor hierarchy shows the origin of macrophages and dendritic cells in early lymphoid development. *Nature Immunology*, *11*(7), 585–593. https://doi.org/10.1038/ni.1889

Golinski, M. L., Demeules, M., Derambure, C., Riou, G., Maho-Vaillant, M., Boyer, O., … Calbo, S. (2020). CD11c+ B Cells Are Mainly Memory Cells, Precursors of Antibody Secreting Cells in Healthy Donors. *Frontiers in Immunology*, *11*. https://doi.org/10.3389/fimmu.2020.00032

Karnell, J. L., Kumar, V., Wang, J., Wang, S., Voynova, E., & Ettinger, R. (2017, November 1). Role of CD11c+ T-bet+ B cells in human health and disease. *Cellular Immunology*, Vol. 321, pp. 40–45. https://doi.org/10.1016/j.cellimm.2017.05.008

Kerkelä, E., Lahtela, J., Larjo, A., Impola, U., Mäenpää, L., & Mattila, P. (2019). *Exploring transcriptomic landscapes in red blood cells, in their extracellular vesicles and on a single-cell level*. https://doi.org/10.21203/rs.2.14503/v1

Kiselev, V. Y., Yiu, A., & Hemberg, M. (2018). Scmap: Projection of single-cell RNA-seq data across data sets. *Nature Methods*, *15*(5), 359–362. https://doi.org/10.1038/nmeth.4644

Kowalczyk, M. S., Tirosh, I., Heckl, D., Rao, T. N., Dixit, A., Haas, B. J., … Regev, A. (2015). Single-cell RNA-seq reveals changes in cell cycle and differentiation programs upon aging of hematopoietic stem cells. *Genome Research*, *25*(12), 1860–1872. https://doi.org/10.1101/gr.192237.115

Kumar, B. V., Ma, W., Miron, M., Granot, T., Guyer, R. S., Carpenter, D. J., … Farber, D. L. (2017). Human Tissue-Resident Memory T Cells Are Defined by Core Transcriptional and Functional Signatures in Lymphoid and Mucosal Sites. *Cell Reports*, *20*(12), 2921–2934. https://doi.org/10.1016/j.celrep.2017.08.078

Paul, F., Arkin, Y., Giladi, A., Jaitin, D. A., Kenigsberg, E., Keren-Shaul, H., … Amit, I. (2015). Transcriptional Heterogeneity and Lineage Commitment in Myeloid Progenitors. *Cell*, *163*(7), 1663–1677. https://doi.org/10.1016/j.cell.2015.11.013

Schraivogel, D., Gschwind, A. R., Milbank, J. H., Leonce, D. R., Jakob, P., Mathur, L., … Steinmetz, L. M. (2020). Targeted Perturb-seq enables genome-scale genetic screens in single cells. *Nature Methods*, *17*(6), 629–635. https://doi.org/10.1038/s41592-020-0837-5

Stuart, T., Butler, A., Hoffman, P., Hafemeister, C., Papalexi, E., Mauck, W. M., … Satija, R. (2019). Comprehensive Integration of Single-Cell Data. *Cell*, *177*(7), 1888-1902.e21. https://doi.org/10.1016/j.cell.2019.05.031

Velten, L., Haas, S. F., Raffel, S., Blaszkiewicz, S., Islam, S., Hennig, B. P., … Steinmetz, L. M. (2017). Human haematopoietic stem cell lineage commitment is a continuous process. *Nature Cell Biology*, *19*(4), 271–281. https://doi.org/10.1038/ncb3493

Velten, L., Story, B. A., Hernandez-Malmierca, P., Milbank, J., Paulsen, M., Lutz, C., … Steinmetz, L. M. (2018, December 21). MutaSeq reveals the transcriptomic consequences of clonal evolution in acute myeloid leukemia. *BioRxiv*, p. 500108. https://doi.org/10.1101/500108

Young, M. D., & Behjati, S. (2020). SoupX removes ambient RNA contamination from droplet-based single-cell RNA sequencing data. *GigaScience*, *9*(12). https://doi.org/10.1093/gigascience/giaa151
